# Whole-genome sequencing of rare disease patients in a national healthcare system

**DOI:** 10.1101/507244

**Authors:** Willem H Ouwehand, on behalf of the NIHR BioResource and the 100,000 Genomes Project

**Author notes:** A full list of authors and their affiliations appears at the end of the manuscript and a list of collaborators and their affiliations appears at the end of the Supplementary Information.

## Abstract

Most patients with rare diseases do not receive a molecular diagnosis and the aetiological variants and mediating genes for more than half such disorders remain to be discovered. We implemented whole-genome sequencing (WGS) in a national healthcare system to streamline diagnosis and to discover unknown aetiological variants, in the coding and non-coding regions of the genome. In a pilot study for the 100,000 Genomes Project, we generated WGS data for 13,037 participants, of whom 9,802 had a rare disease, and provided a genetic diagnosis to 1,138 of the 7,065 patients with detailed phenotypic data. We identified 95 Mendelian associations between genes and rare diseases, of which 11 have been discovered since 2015 and at least 79 are confirmed aetiological. Using WGS of UK Biobank^1^, we showed that rare alleles can explain the presence of some individuals in the tails of a quantitative red blood cell (RBC) trait. Finally, we reported 4 novel non-coding variants which cause disease through the disruption of transcription of *ARPC1B*, *GATA1*, *LRBA* and *MPL*. Our study demonstrates a synergy by using WGS for diagnosis and aetiological discovery in routine healthcare.

---

Rare diseases affect approximately 1 in 20 people, but only a minority of patients receive a genetic diagnosis^2^. Approximately 10,000 rare diseases are known, but fewer than half have a resolved genetic aetiology^3^. Even when the aetiology is known, the prospects for diagnosis are severely diminished by a fragmentary approach to phenotyping and the restriction of genetic testing to a disease-specific panel of genes. On average, a molecular cause is determined after three misdiagnoses and 16 physician visits over a “diagnostic odyssey” lasting more than two years^4^. However, recent developments in WGS technology mean it is now possible to perform comprehensive genetic testing systematically in an integrated national healthcare system. The large-scale implementation of WGS for diagnosis will also enable the discovery of new genetic aetiologies, through the identification of novel causal mutations in the coding and non-coding parts of the genome.

In a pilot study for the 100,000 Genomes Project supported by the National Institute for Health Research (NIHR), we have performed WGS of 13,037 individuals enrolled at 57 National Health Service (NHS) hospitals in the United Kingdom and 26 hospitals in other countries (Fig. 1a, Extended Data Fig. 1a, Supplementary Table 1) in three batches, to clinical standard (Fig. 1b). The participants were distributed approximately equally between the sexes (Supplementary Table 1) and their distribution across ethnic groups closely matched that reported in the UK census (Fig. 1c; https://www.ons.gov.uk/census/2011census). In total, 9,802 participants (75%) were affected with a rare disease or had an extreme measurement of a quantitative trait, of which 9,024 were probands and 778 were affected relatives. Each participant was assigned to one of 18 domains (Table 1): 7,388 individuals to one of 15 rare disease groups, 50 individuals to a control group, 4,835 individuals to a Genomics England Limited (GEL) group and 764 individuals to a group of UK Biobank participants with extreme red blood cell indices (Supplementary Information). The rare disease domains covered pathologies of a wide range of organ systems and each had pre-specified inclusion and exclusion criteria (Supplementary Information, Supplementary Table 1, Extended Data Fig. 1b). The variation in sample size across domains was primarily due to differences in recruitment rate, which limited the efficiency of the experimental design. We subsequently collected detailed phenotypic information, through web-based data capture applications, in the form of Human Phenotype Ontology (HPO) terms for 13 of the rare disease domains (Fig. 2a,b, Extended Data Fig. 1c). Patients with diverse diagnoses were enrolled to the GEL domain, together with healthy family members, but only the affection status of these participants were available for this study. In addition, HPO-coded phenotypes were not collected for Leber Hereditary Optic Neuropathy (LHON) and Ehler-Danlos and Ehler-Danlos-like Syndromes (EDS) patients. In total, 19,605 HPO terms were selected to describe patient phenotypes. Quantitative data were transcribed to HPO terms using domain-specific rules, while free text was transcribed manually.

**Table 1.**
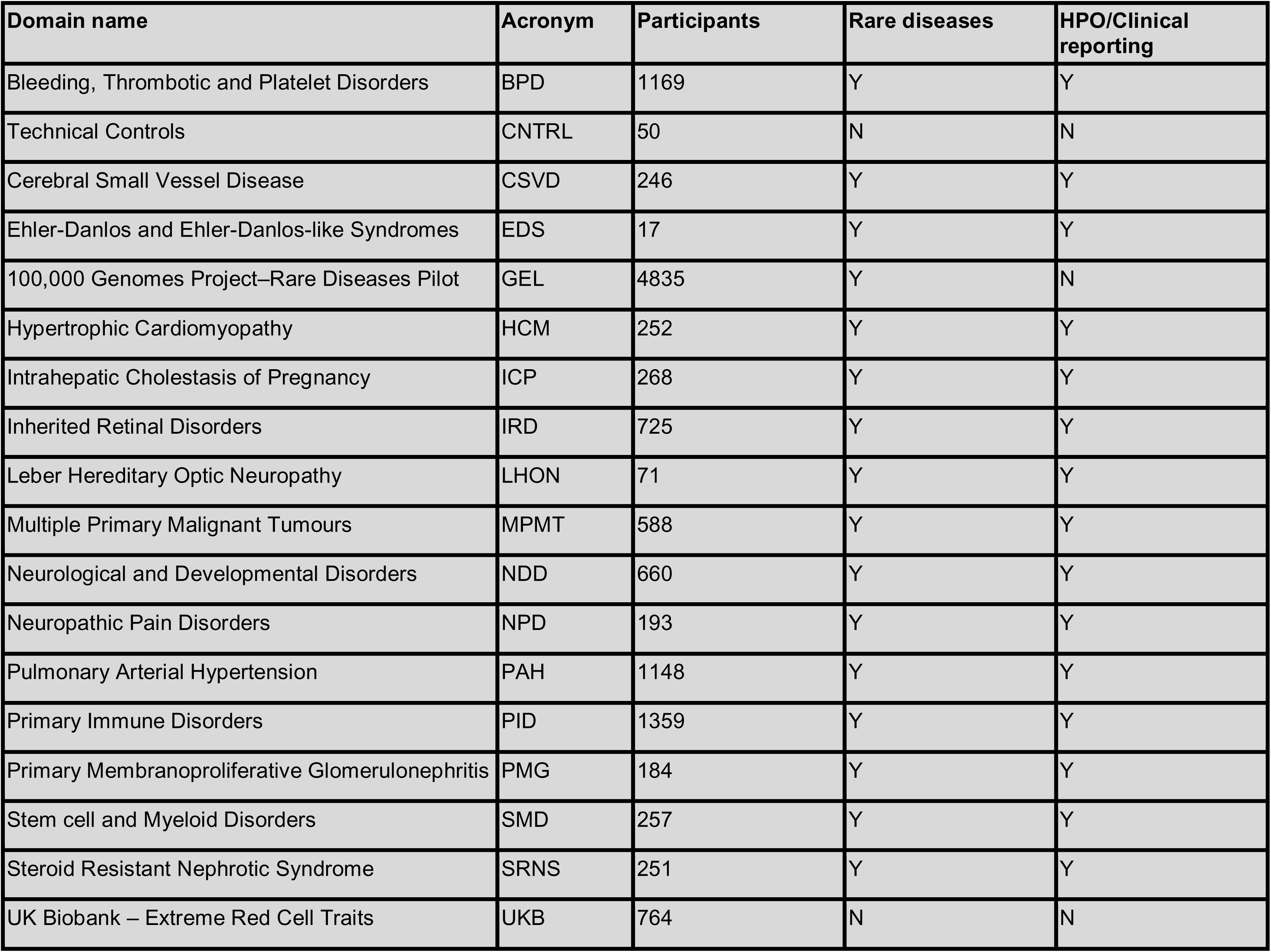
Study domain names, acronyms, numbers of participants, whether the domain included cases with a rare disease and whether domain participants were assigned HPO terms for diagnostic reporting.

Following bioinformatic quality control (QC) and data analysis (Extended Data Fig. 2–4), we identified 172,005,610 short variants, of which 157,411,228 (91.5%) were single nucleotide variants (SNVs) and 14,594,382 (8.5%) were indels up to 50bp long (Extended Data Fig. 5). 48.6% and 40.8% of the SNVs and indels, respectively, were absent from all major variant databases (Fig. 1e). 54.8% of the variants were observed in only one member of the maximal set of 10,259 unrelated participants, of which 82.6% were novel. Only 9.08% of novel variants were observed in more than one member of the unrelated set, typically in sets of individuals with recent common ancestry (Fig. 1f). SNVs and indels common in our dataset were well represented in genetic databases but, in accordance with theory, the vast majority of the variants we observed were very rare and most were uncatalogued. We called 24,436 distinct large deletions (>50bp) by synthesising inferences from two algorithms across individuals. We also called more complicated types of structural variant, such as inversions, but they were called unreliably and could not be confidently aggregated across individuals (Supplementary Information). We used the WGS data to determine that only 13 (0.1%) individuals had non-standard sex chromosomal karyotypes (Extended Data Fig. 3e–g). Using the high quality variant calls, we inferred a wide range of bioinformatically estimated family sizes, in keeping with differences in enrolment strategies (Supplementary Information), of which most comprised singletons (Fig. 1d).

We issued clinical reports for 1,103 distinct causal variants (731 SNVs, 264 indels, 102 large deletions, 6 other structural variants) affecting 303 genes (Extended Data Fig. 5). 266 of the 995 SNVs and indels (26.7%) were absent from the Human Gene Mutation Database (HGMD) and from the set of variants in ClinVar having a pathogenic or likely pathogenic interpretation and no benign interpretations. We identified strong evidence (posterior probability (PP) > 0.75) for 99 genetic associations between rare variants and groupings of patients with similar phenotypes using the Bayesian genetic association method, BeviMed^5^. Of these 99 associations, 62 are consistent with firmly established evidence and a further 11 have been reported in the literature since 2015, either by us or by other researchers. We showed that genetic associations with the extremes of a quantitative trait can identify genes in which mutations cause Mendelian pathologies. Finally, we used a novel method, RedPop, to call cell-type specific regulatory elements (REs) from open chromatin and histone modification data. We combined these calls with cell-type specific transcription factor binding information to identify four pathogenic rare non-coding variants that cause disease by disrupting the proper regulation of gene expression.

## Summary of clinical findings

For each of the 15 rare disease domains, we established a list of diagnostic-grade genes (DGGs) and lists of their corresponding transcripts on the basis of the scientific literature (Supplementary Information). The number of DGGs for each domain ranged from two for Intrahepatic Cholestasis of Pregnancy (ICP) to 1,423 for Neurological and Developmental Disorders (NDD). The DGG lists were not mutually exclusive because some genes harbour mutations that cause pathologies compatible with the enrolment criteria of multiple domains (Fig. 2c). Twelve multidisciplinary teams (MDTs) with domain-specific expertise examined the rare variants observed in DGGs in the context of the HPO phenotypes of the carriers. They categorised a subset of the variants as *pathogenic* or *likely pathogenic* following standard guidelines^6^ and assessed their allelic contribution to disease as *full* or *partial*. A variant’s contribution was considered to be at least partial if, given all other known variants in the case, it was considered to have a disease determining consequence. A conclusive molecular diagnosis was returned for 1,138 of the 7,065 (16.1%) patient records reviewed and those diagnoses featured 1,103 distinct causal variants (Supplementary Table 2). One quarter of the reports featured variants in *BMPR2*, *ABCA4* and *USH2A* and a further quarter featured variants in a group of 18 DGGs. The remaining half of the clinical reports concerned variants spread across 306 DGGs, which often featured in a single report (Fig. 2d, Extended Data Fig. 6). The diagnostic yield by domain ranged from no patients out of 184 (0%) for Primary Membranoproliferative Glomerulonephritis (PMG) to 391 patients out of 725 (53.9%) for Inherited Retinal Disease (IRD). The variability of diagnostic yield can be attributed to heterogeneity in: phenotypic and genetic pre-screening before enrolment, the genetic architecture of diseases and prior knowledge of genetic aetiologies.

Clinical reporting was enhanced by the use of PCR-free WGS with a mean autosomal depth >35X instead of whole-exome sequencing (WES). For example, a causal SNV encoding a start loss of *HPS6* in a case with Hermansky-Pudlak syndrome was identified by WGS but not identified by WES prior to the study. We compared the coverage obtained from the WGS samples to coverage obtained from research WES of UK Biobank samples (https://www.biorxiv.org/content/10.1101/572347v1), INTERVAL samples^7^ and samples from the Columbia University exome sequencing study for chronic kidney disease^8^ (Supplementary Information). Although less costly to generate per sample, all WES datasets exhibited much greater variation in coverage within and between genomic sites harbouring known pathogenic SNVs or indels than WGS (Extended Data Fig. 7). Of the 938 distinct autosomal aetiological SNVs and indels reported in this study, 25–99 (2.67%–10.5%) had insufficient coverage in WES for reliable genotyping, depending on dataset (Extended Data Fig. 7). Moreover, deletions spanning only a few short exons or part of a single exon are not reliably called by WES^9,10^. Of the 102 distinct large deletions that we reported (length range 203bp–16.80Mb, mean 786.33Kb, median 15.91Kb), 22 (21.6%) overlapped only one exon.

Our recent genetic discoveries have informed treatment decisions: 27 patients with early-onset dystonia due to variants in *KMT2B* can be treated by deep brain stimulation^11^; cases with *DIAPH1*-related macrothrombocytopenia and deafness^12^ can have their platelet count restored to a safe level in a preoperative setting with Eltrombopag^13^; and a case of severe thrombocytopenia accompanied by myelofibrosis and bleeding caused by a gain-of-function variant in *SRC*^14^ was cured by an allogeneic haematopoietic stem cell transplant. In addition, our diagnoses have helped stratify patient care: patients with Primary Immune Disorders (PID) due to variants in *NFKB1*, which we have shown are the commonest monogenic cause of combined variable immunodeficiency (CVID)^15^, have unexplained splenomegaly and an increased risk of cancer; 27 cases from the Bleeding, Thrombotic and Platelet Disorders (BPD) domain with isolated thrombocytopenia caused by variants in *ANKRD26*, *ETV6* or *RUNX1* have an increased risk of malignancy^16,17,18^ compared to 19 cases with benign thrombocytopenia due to variants in *ACTN1*, *CYCS* or *TUBB1*^19^. Furthermore, our discoveries have improved the accuracy of prognosis, which is worse for patients with Pulmonary Arterial Hypertension (PAH) if the cause is mutations in *BMPR2*^20^ or *EIF2AK4*^21^, while the impact of mutations in *ATP13A3*, *AQP1*, *GDF2* and *SOX17*, genes which we have recently reported as aetiological^22^, remains to be determined.

Quantitative intermediate phenotypes can contain information that is useful for understanding genetic aetiology in difficult to diagnose patients. We examined WGS read alignments for patients with complete absence of a protein encoded by a DGG but carrying an explanatory variant call on only one haplotype. Two patients with a severe unexplained bleeding disorder due to the absence of αIIbβ 3 integrin on their platelet membranes carried two different complex variants in intron 9 of *ITGB3*: a tandem repeat and an SVA retrotransposon which was not called by either of the two structural variant callers we employed, but was discernible due to an excess of improperly mapped reads and confirmed by long-read nanopore sequencing (Extended Data Fig. 8a–e). The third patient had an absence of RhD and RhCE proteins on the membrane of her red cells leading to severe haemolytic anemia. This was due to a large tandem repeat in *RHAG*, which encodes the Rh-associated glycoprotein (Extended Data Fig. 8f).

### Discovery of rare variants associated with rare diseases

Several cases with similar aetiologies are typically needed to make a novel discovery in rare disease genetics. Cases can be aggregated across siloed studies, using services such as Matchmaker Exchange (MME)^23^. We used MME to identify novel aetiologies for *SLC18A2*^24^ and *WASF1*^25^ (Supplementary Information). However, in the context of a study of a unified healthcare system, it is possible to make discoveries by statistical analyses of large patient collections.

We applied the statistical method BeviMed^5^ to identify genetic associations between gene loci and rare diseases under various modes of Mendelian inheritance (Supplementary Information). We defined a set of phenotypic tags for each domain to determine a set of case/control groupings for BeviMed. Groups of cases were assigned the same tag if their phenotypes were *a priori* judged compatible with a shared genetic aetiology of disease (Supplementary Table 3). The number of unrelated cases in each tag group ranged from three for Roifman syndrome to 1,101 for PAH. For each gene-tag pair, we compared the genotypes at rare variant sites between unrelated individuals with the tag (cases) and unrelated individuals without the tag (controls). We considered a PP of association > 0.75 to be strong evidence supporting a genetic aetiology. Additionally, for each analysis BeviMed inferred a conditional PP over the mode of inheritance, a conditional PP over the molecular consequence class of variants mediating disease risk (e.g. 5’ UTR variants or predicted loss-of-function variants) and conditional PPs of pathogenicity for each specific variant. These quantities were used to compare established to inferred modes of inheritance and to estimate the number of cases attributable to variants in each gene^5^.

We inferred strong evidence for an association between each of 95 genes, spanning nine domains, and one of 29 phenotypic tags. These genes included 68 established DGGs, 11 DGGs discovered since 2015^15,26,27,22,28,29,30,31,22,32,33^ and 16 candidates requiring further investigation (Fig. 3; Supplementary Table 3). Thus, 79 of 95 genetic associations are confirmed, which sets a lower bound on the observed positive predictive value (PPV) of 83%, which is broadly in line with an ancestry-controlled statistical estimate of the study-wide PPV of 79% (Supplementary Information). We estimated that 611.3 cases can be explained by rare variants in the 79 genes with a confirmed association, 115.6 of which can be explained by the association between variants in *BMPR2* and PAH. 51 of the 95 genetic associations relied only on evidence from alleles carried by single cases, showing the power of joint statistical modelling of rare variants. Only three of the unconfirmed associations relied on evidence from alleles carried by more than one case, demonstrating the robustness of the results to cryptic relatedness. For one gene (*GP1BB*), the mode of inheritance inferred by BeviMed differed from that established in the literature, challenging long-held assumptions^34^. These results and other findings from this project^22,35,36,37,25,38,15,39,40,10,19,41,42,25,43,11,44,12,45^ show that a unified analysis of standardised homogeneously collected genetic and phenotypic data from large cohorts of different rare disease domains is a powerful approach for genetic discovery.

### Polygenic and rare variant associations with the extremes of a quantitative trait in UK Biobank

Several rare diseases (e.g. familial hypercholesterolaemia, CVID, thrombocytopenia, von Willebrand disease) are diagnosed and clinically characterised by reference to a quantitative trait that acts as a causal intermediate (or close proxy) for pathology and symptoms. Mutation-selection equilibrium ensures strong negative selection in the extreme tails of heritable quantitative traits, so individuals in the tails should have lower fecundity, perhaps due to greater risk of disease. We sought to identify genes likely to carry mutations causing RBC pathologies by computing a univariate quantitative summary of baseline RBC full blood count (FBC) traits in the UK Biobank participants of European ancestry. We aimed to develop a red cell phenotype capturing as much rare-variant heritability as possible. To achieve this, we used the joint distribution of estimated effect sizes from published GWAS associations between variants with MAF < 1% and four mature RBC FBC traits as a model for the effect of causal rare alleles we hoped to identify by WGS^46^ (Fig 4a). We successfully sequenced 764 participants, 383 of which were extreme for the left tail of the phenotype, corresponding to a low RBC count (RBC#) and a high mean cell volume (MCV), and 381 of which were extreme for the right tail of the phenotype, corresponding to a high RBC# and a low MCV (Fig. 4b,c).

The distribution of a polygenic predictor of the quantitative phenotype, derived from genetic variants known to be associated with RBC# and MCV exhibits left and right shifts from the population distribution in the respectively named tails (Fig. 4d). However, these shifts are not as strong as those predicted by Gaussian variance components modelling, a discrepancy which could be explained partly by rare alleles generating excess density in the tails (kurtosis 6.9). A WGS GWAS of an ordinal outcome (left tail, unselected, right tail) did not yield novel associations. Consequently, we treated each of the tail groups as a set of cases in a BeviMed analysis, identifying 12 genes with PP evidence for an association stronger than 0.4, a liberal threshold (Fig. 4e). *HBB* and *TFRC* can be considered positive controls, as they are known to carry mutations causing Mendelian microcytic anaemias. Other genes, including *CUX1* and *ALG1* are biologically plausible candidates. These results (Supplementary Table 3) indicate that the analysis of quantitative extremes in apparently healthy population samples may identify medically relevant loci unidentified by GWAS for quantitative traits^46, 47^.

### Aetiological variants in regulatory elements

Recent statistical modelling suggests that only a small proportion of the burden of heritable neurodevelopmental disorders can be attributed to *de novo* pathogenic SNVs in non-coding elements^48^. Nevertheless, rare variants in REs are known to cause disease by disrupting transcription or translation^49,50,51^. We searched for aetiological variants in the REs of 246 DGGs implicated in recessive haematopoiesis-related disorders. Firstly, we defined a set of active REs we named a ‘regulome’ for each of six blood progenitor and mature blood cell types. We achieved this by merging transcription factor binding sites identified by ChIP-seq with genomic regions called by RedPop, a new detection method exploiting the anti-covariance of ATAC-seq and H3K27ac ChIP-seq coverage in REs (Supplementary Information). We linked the REs to genes on the basis of genomic proximity and promoter capture Hi-C data^52^. Secondly, we assigned each regulome to one or more of the BPD, PID and Stem Cell and Myeloid Disorders (SMD) domains, depending on the relevance of the corresponding cell types to these domains (Supplementary Table 3). Finally, we searched for cases carrying a rare homozygous or hemizygous deletion of an RE active in a cell type assigned to the domain of the case and which was linked to a DGG of that domain. We also searched for heterozygous deletions meeting these criteria that were in compound heterozygosity with a rare coding variant in a DGG linked to the deleted element (Fig. 5a). These approaches explained three cases: a PID patient carrying a deletion overlapping the 5’ UTR region of *ARPC1B* in compound heterozygosity with a frameshift variant in the same gene (Thaventhiran *et al*, under review), a nine year old boy with autism spectrum disorder and thrombocytopenia carrying a hemizygous deletion of a *GATA1* enhancer on the X chromosome, and a male with several autoimmune-mediated cytopenias carrying a homozygous deletion of intronic CTCF binding sites^53^ of *LRBA*.

The X-linked deletion in the boy with autism (Extended Data Fig. 9a–b) removed an element regulating *GATA1* as well as exons 1-4 of *HDAC6*. He had a persistently low platelet count (52×10^9^/l), a mean platelet volume in the 99.9^th^ percentile of the distribution for UK Biobank males (Fig. 5b)^54^ and normal RBC parameters except for mild dyserythropoiesis. Electron microscopic imaging of his platelets showed reduced α-granule content (Extended Data Fig. 9c–e). Culture of his stem cells recapitulated ineffective formation of platelets by megakaryocytes (Extended Data Fig. 9f–k). Macrothrombocytopenia, reduced α-granule content, ineffective platelet formation and dyserythropoiesis are all characteristic of patients with pathogenic coding mutations of *GATA1*^55, 56^. His platelets contained reduced GATA1 (Fig. 5g), consistent with reduced transcription due to deletion of the *GATA1* enhancer^57^. HDAC6 is the major deacetylase for removing the acetyl group from Lys40 of α-tubulin, which is located in polymerized microtubules^58^. The absence of HDAC6 in the child was accompanied by extremely high expression levels of acetylated α-tubulin in his platelets (Fig. 5e), concordant with observations of *Hdac6* knockout mice^59^. This aberrant acetylation is associated with bleeding^59^ and altered emotional behaviour^60^ in mice. Thus, the reduced expression of *GATA1* and the absence of HDAC6 jointly cause a new syndrome of macrothrombocytopenia accompanied by neurodevelopmental problems.

The patient with a homozygous deletion of a CTCF binding site in the first intron of *LRBA* presented with a pancytopenia, characterised mostly by neutropenia and anaemia, and complicated by periods of thrombocytopenia. These cytopenias were mediated by autoantibodies due to a loss of tolerance for multiple autoantigens, which is characteristic of patients with reduced *LRBA* function^61^.

We adapted our approach to solving cases caused by non-coding deletions to search for non-coding SNVs with a CADD^62^ score > 20, in the presence of a high-impact coding variant in compound heterozygosity in the assigned DGG. This approach identified two potentially aetiological SNVs in elements assigned to *AP3B1* and *MPL*, and we studied the 10 year old male patient carrying the latter mutation in more detail. *MPL* encodes the receptor for the megakaryocyte growth factor thrombopoietin^63^. Loss of *MPL* causes chronic amegakaryocytic thrombocytopenia in humans^64^ and *Mpl* knockout mice have severe thrombocytopenia^65, 66^. The SNV (chr1:43803414 G>A) was in an RE detected by RedPop, the activity of which is specific to megakaryocytes in blood cell physiology (Extended Data Fig. 10), had a CADD score of 21.8, was absent from gnomAD, and was in compound heterozygosity with a deletion of exon 10 of *MPL*, which was inherited from the patient’s mother (Extended data, Fig. 10a,b,c). A luciferase reporter assay showed approximately 50% reduced promoter activity for the A allele compared to the reference allele (Extended Data Fig. 10d). As a result, platelet MPL levels were significantly reduced in the patient compared to controls (Extended Data Fig. 10e). In contrast to *MPL*-null patients^67^, who are extremely thrombocytopenic because their bone marrow is almost devoid of megakaryocytes and eventually suffer haematopoietic stem cell exhaustion, this boy had platelet counts which stabilised around 45×10^9^/l and a marrow that was only moderately depleted of megakaryocytes. As the regulatory SNV does not abolish *MPL* transcription completely (Extended Fig. 10c), the boy has a milder clinical phenotype than *MPL*-null cases.

## Discussion

Before now there has been limited integration between clinical genetic testing services and aetiological studies of rare diseases on a national scale. We have shown that WGS in a universal national healthcare system can tackle these two objectives concurrently (Fig. 1a). This synergy can only be achieved if sequencing data from explained cases (Fig. 2), unexplained cases and unaffected individuals are analysed jointly and if consent to contact participants for follow-up studies has been obtained at enrolment. We have shown that long-read sequencing can aid the identification of complex structural variants, which can still be called unreliably by short-read WGS. We have demonstrated the utility of data aggregation and sharing through the number of genetic associations we have found across a diversity of rare diseases (Fig. 3). This study follows on from large-scale whole-exome and shallow genome sequencing studies in the UK^68, 69^ and has been the blueprint for the UK’s 100,000 Genomes Project, which recently completed sequencing. The NHS plans to increase provision of WGS-based diagnostics from 8,000 to 30,000 samples per month. To achieve this aim, it has reduced the number of clinical genomics laboratories to seven, each servicing approximately 8 million people. It has also introduced a unified and consistent WGS and informatics infrastructure for these seven hubs and is providing training in genomics to NHS staff. We have initiated WGS of UK Biobank participants to study individuals with extreme values for a quantitative phenotype. Extreme trait values may be the result of measurement error, extreme polygenic loads^47^ or rare genetic variation and such individuals are typically excluded from GWAS studies. We have shown that genetic associations with the tail of a quantitative distribution can identify genes mediating Mendelian pathologies in the same domain of human biology (Fig. 4). The forthcoming WGS of 0.5 million UK Biobank participants provides an opportunity to study other traits following similar approaches. Finally, we have provided examples of rare variants causing disease by disrupting non-coding REs of the genome. The reliability and affordability of WGS and the availability of cell-type specific epigenetic data make the exploration of the non-coding genome (Fig. 5, Extended Data Fig. 10) a promising focus for future research in unresolved rare disorders for which the aetiological cell types are known.

## Supporting information

Legends for Main Figures

Legends for Extended Data Figures

Legends for Supplementary Tables

## Methods

### Enrolment, research ethics and consent

Patients with rare diseases and their close relatives were enrolled to the NIHR BioResource (NBR) as part of a pilot study for the 100,000 Genomes Project. For this study, 15 rare disease domains were approved after review by the Sequencing and Informatics Committee of the NBR. Enrolment of participants for this pilot study was coordinated by the University of Cambridge, started in December 2012 and was completed in March 2017. In addition, samples from a second rare diseases pilot study, coordinated by GEL, are included together with a number of control samples and samples from the UK Biobank cohort^70^. The NBR–Rare Diseases study was coordinated by the University of Cambridge. Participants were recruited mainly at NHS Hospitals in the UK, but also at hospitals overseas (Supplementary Table 1, Extended Data Fig. 1a). All 13,187 participants provided written informed consent, either under the East of England Cambridge South national research ethics committee (REC) reference 13/EE/0325 or under alternative REC-approved studies. Obtaining consent for overseas samples was the responsibility of the respective principal investigators at the hospitals where enrollment took place. The NBR retained blank versions of the consent forms from overseas participants and a material transfer agreement was applied to regulate the exchange of samples and data between the donor institutions and the University of Cambridge.

### Clinical and laboratory phenotype data

Staff at hospitals responsible for enrolment were provided with the eligibility criteria for their respective domains as described above in the domain descriptions. The clinical and laboratory phenotype data were captured through case report forms (CRF) by paper questionnaires or by online CRF data capture applications and deposited in the NBR study database. Online data capture allowed for the free entry of HPO terms^71^ by staff at the enrolment centre and data from paper questionnaires were transformed into HPO terms by the study coordination office. Free text entries were transformed into HPO terms where feasible. An overview of the HPO data obtained for the 15 NBR rare disease domains is depicted in Extended Data Fig. 1c,d.

### DNA sequencing

Samples were received as either DNA extracted from whole blood or as whole blood EDTA samples, which were used for extraction at the NBR laboratory in Cambridge. Samples were tested for adequate concentration (Picogreen), quality controlled (QC) for DNA degradation (gel electrophoresis) and purity (OD 260/280; Trinean) before selection for WGS. DNA samples were prepared at a minimum concentration of 30 ng/μl in 110 μl, visually inspected for degradation and had to have an OD 260/280 between 1.75 and 2.04. They were then prepared in batches of 96 and shipped on dry ice to the sequencing provider (Illumina Inc, Great Chesterford, UK). Further sample QC was performed by Illumina to ensure that the concentration of the DNA was > 30 ng/μl and that every sample generated high quality genotyping results (Illumina Infinium Human Core Exome microarray). Samples with a repeated array genotyping call rate < 0.99, high levels of cross-contamination, mismatches with the declared gender that could not be resolved by further investigation, or for which consent had been withdrawn, were excluded from WGS (n=59). The genotyping data were also used for positive sample identification and sample identity was verified before data delivery. In short 0.5 μg of the DNA sample was fragmented using Covaris LE220 (Covaris Inc., Woburn, MA, USA) to obtain an average size of 450bp DNA fragments. DNA samples were processed using the Illumina TruSeq DNA PCR-Free Sample Preparation kit (Illumina Inc., San Diego, CA, USA) on the Hamilton Microlab Star (Hamilton Robotics, Inc, Reno, NV, USA). The final libraries were checked using the Roche LightCycler 480 II (Roche Diagnostics Corporation, Indianapolis, IN, USA) with KAPA Library Quantification Kit (Kapa Biosystems Inc., Wilmington, MA, USA) for concentration. From February 2014 to June 2017 three read lengths were used: 100bp, 125bp and 150bp (377, 3,154 and 9,656 samples, respectively). Samples sequenced with 100bp and 125bp reads utilised three and two lanes of an Illumina HiSeq 2500 instrument, respectively, while samples sequenced with 150bp reads utilised a single lane of a HiSeq X instrument. At least 95% of the autosomal genome had to be covered at 15X and a maximum of 5% of insert sizes had to be less than twice the read length. Following sample and data QC at Illumina, 13,187 sets of WGS data files were received at the University of Cambridge High Performance Computing Service (HPC) for further QC.

### WGS data processing pipeline

The WGS data for the 13,187 samples returned by the sequencing provider underwent a series of processing steps (Extended Data Fig. 2), described in detail in the Supplementary Information. Briefly, the samples were sex karyotyped and pairwise kinship coefficients were computed. This information was used to check for repeat sample submissions and sample swaps. Additionally, four further QC checks were applied to ensure the SNVs and indels were of a high standard. Overall, 150 samples (1.1%) were removed, leaving a dataset of 13,037 samples for downstream analysis. The 13,037 individuals were assigned one of the following ethnicities: European, African, South Asian, East Asian or Other. Pairwise relatedness adjusted for population stratification was then computed and used to generate networks of closely related individuals and to define a maximal set of 10,259 unrelated individuals. The variants in the 13,037 individuals were left-aligned and normalised with bcftools, loaded into our HBase database and filtered on their overall pass rate (OPR), defined in the Supplementary Information. The sex karyotypes, the ethnicities and the relatedness estimates were used, along with enrolment information, to annotate the samples and variants. Samples were annotated with: affected/unaffected status, membership of the set of probands, membership of the maximal unrelated set, ethnicity and sex karyotype. Variants were annotated with CellBase consequence predictions, HGMD information where available and population-specific allele frequencies.

### Pertinent findings

For each of the 15 rare disease domains (i.e. all domains except UKB, CNTRL and GEL) a list of DGGs was generated by domain-specific experts. Genes were included in the lists if there was a high enough level of evidence in the literature for gene-disease association. The 2,497 gene/domain pairs, encompassing 2,073 unique DGGs across all domains, were manually curated and annotated with the relevant RefSeq and/or Ensembl transcript identifiers to support variant reporting. Transcripts were selected based on, by order of priority, community input, presence in the Locus Reference Genomic (LRG) resource^72^ or designation as canonical in Ensembl. Variants (SNVs, indels) were shortlisted if (i) their MAF in control populations^73^ was < 1/1,000 for putative novel causal variants and < 25/1,000 for variants listed as disease-causing in HGMD, (ii) their predicted impact according to the Variant Effect Predictor^74^ was “HIGH” or “MODERATE” or if the consequences with respect to the designated transcript included one of “splice_region_variant” or “non_coding_transcript_exon_variant” if the variant was in a non-coding gene, (iii) the variant affected a DGG relevant to the patient’s disease. Variants with more than 3 alleles or a MAF >= 10% in the diseases cohort were discarded to, respectively, guard against errors in repetitive regions and remove potential systematic artefacts. The above filtering criteria were applied universally to all domains, except for ICP which adopted a higher MAF threshold of 3% for both novel and previously reported variants. The higher threshold accounted for causal variants being present in the male and non-child bearing female population. This strategy reduced the number of variants for review by the MDT from about 4 million per person to fewer than 10, while confidently retaining known regulatory or moderately common pathogenic variants. For each affected participant with prioritised variants, the variant calls, HPO-coded phenotype and the relevant metadata (unique study numbers; referring clinician and hospital; self-declared gender and genetically inferred sex, ancestry, relatedness, and consanguinity level) were transferred to Congenica Inc (Cambridge, United Kingdom) for visualisation in the Sapientia^TM^ web application during MDT meetings. MDTs brought together experts from different hospitals across the UK and abroad, and typically consisted of an experienced clinician with domain-specific knowledge, a scientist with experience in clinical genomics, a clinical bioinformatician and a member of the reporting team. Assignment of the level of pathogenicity followed the American College of Medical Genetics guidelines^6^ and variants (V) were marked in Sapientia^TM^ as pathogenic, likely pathogenic or of uncertain significance (VUS). Only pathogenic and likely pathogenic variants were systematically reported and VUSs were reported at the MDT’s discretion. As per REC-approved study protocol, secondary findings (e.g. breast cancer pathogenic variants in *BRCA1* in patients not presenting with this phenotype) were not reported.

### Genetic association testing in genes

We used the BeviMed statistical method^5^ to identify genetic associations with rare diseases in our dataset. Each run of BeviMed requires the definition of a set of cases and controls, all of which should be unrelated with each other, and a set of rare variants to include in the inference. To achieve adequate power, the cases should be chosen such that they potentially share a common genetic aetiology (e.g. because the phenotypes are similar) and the rare variants should be chosen such that they potentially share a mechanism of action on phenotype (e.g. because they are predicted to have a similar effect on a particular gene product). BeviMed computes PP values of no association, dominant association and recessive association and, conditional on dominant or recessive association, it computes the PP that each variant is pathogenic. We can impose a prior correlation structure on the pathogenicity of the variants that reflects competing hypotheses as to which class of variant is responsible for disease. These classifications typically group variants by their predicted consequences. The class of variant responsible can then be inferred by BeviMed, thereby suggesting a particular mechanism of disease. The methodology is described in further detail in the Supplementary Information and in reference^5^.

### Regulome analysis

We applied the BLUEPRINT protocol for ChIP-seq data analysis (http://dcc.blueprint-epigenome.eu/#/md/chip_seq_grch37). We defined regulomes for activated CD4+ T cells (aCD4), B cells (B), erythroblasts (EB), megakaryocytes (MK), monocytes (MON) and resting CD4+ T cells (rCD4). For each cell type, we used open chromatin data (ATAC-seq or DNAse-seq) and histone modification data (H3K27ac) to identify REs using the RedPop method (see below). Additionally, for MK and EB, we had access to the following transcription factor (TF) ChIP-seq data, which were used to call peaks (see below) and supplement the regulomes: FLI1, GATA1, GATA2, MEIS1, RUNX1, TAL1 and CTCF for MK; GATA1, KLF1, NFE2 and TAL1 for EB; and CTCF for MON and B. For each cell type, the regulome build process proceeded as follows: 1. Call RedPop regions using ATAC-seq/DNAse-seq and H3K27ac-seq data; 2. Call TF/CTCF binding peaks using ChIP-seq data if available and obtain enrichment scores; 3. Discard TF regions with an enrichment score < 10 unless they overlap between at least two different TFs; 4. Collapse overlapping features to obtain a single genomic track; 5. Merge features within 100bp of each other. Each regulome feature was assigned a gene label using either gene annotations from Ensembl (v75) or a compendium of previously published promoter capture Hi-C data (pcHi-C)^52^ as follows: 1. Assign to a gene if the feature overlaps the gene or the region up to 10Kb either side of the gene body; 2. Assign to a gene if the feature overlaps the gene’s pcHi-C ‘blind’ spot. This region is defined by three *Hind*III restriction fragments, incorporating the capture fragment overlapping target gene TSS, and 5’ and 3’ adjacent fragments; 3. Assign to a gene if the feature overlaps a linked promoter interacting region identified using pcHi-C in the same cell type.

### Functional analysis of the *GATA1* enhancer/*HDAC6* deletion

The *GATA1* enhancer/*HDAC6* deletion was confirmed by PCR using primers HDAC6-F: 5’-catcttcaagaggatcagagg and HDAC6-R: 5’-catagctagacactggtt. Electron microscopy for platelets was performed as described^55^. Immunostaining of resting and fibrinogen spread platelets was performed as described^44^ and analyzed by Structured Illumination Microscopy (SIM, Elyra S.1, Zeiss, Heidelberg, D.E). Total protein lysates were obtained from platelets for immunoblot analysis as described^75^. The following antibodies were used for SIM and immunoblot analysis: rabbit anti-HDAC6 (clone D2E5, Cell Signaling technology, Danvers, MA, USA), mouse anti-acetylated tubulin antibody (clone 6-11B-1, Sigma, St Louis, MO, USA), mouse anti-alpha-tubulin (A11126, Thermo Fisher Scientific, Waltham, MA, USA), rabbit anti-VWF (Dako, Aligent Technologies, Leuven, BE), mouse anti-CD63 and rat anti-GATA1 N6 (Santa Cruz Biotechnology, Dallas, TX, USA), rabbit anti-GATA1 (NF that was produced against recombinant N-terminal zinc finger^76^, rabbit anti-GAPDH (14C10, Cell Signaling) and anti-β3 integrin (sc-14009; Santa Cruz Biotechnology).

### MPL expression on platelets

The level of MPL protein on the platelet membrane was measured by flow cytometry (Beckman Coulter FC500) using the monoclonal antibodies: APC-labelled IgG1 against CD42b (clone HIP1, BD Pharmingen, number: 551061), PE-labelled IgG1 against CD110 (clone REA250, Miltenyi Biotec) and a PE-labelled isotype control (clone MOPC-21, BD Pharmingen, number: 555749). In short, a sample of EDTA anticoagulated blood was incubated with anti-CD110 (or control) and anti-CD42b for 30 minutes. Mean fluorescence intensity (MFI) produced by the anti-CD110 was measured by flow cytometry on cells gated on the CD42b APC signal, side and forward scatter.

### Nanopore sequencing

Oxford Nanopore-based sequencing of long-range PCR-amplified target DNA was performed as previously described^77^ with the aim to resolve the genetic architecture of intron 9 of *ITGB3* in a case with Glanzmann’s thrombasthenia. The flow cell ran for 3 hours, and the mean coverage was 863,986X.

### Code availability

Code to run HBASE is available from https://github.com/mh11/VILMAA. The RedPop software package is available from https://gitlab.haem.cam.ac.uk/et341/redpop/.

### Data availability

Genotype and phenotype data from the 4,835 participants enrolled in the NIHR BioResource for the 100,000 Genomes Project–Rare Diseases Pilot can be accessed by seeking access via Genomics England Limited following the procedure outlined at: https://www.genomicsengland.co.uk/about-gecip/joining-research-community/. The genotype data for the 764 UK Biobank samples will be made available through a data release process overseen by UK Biobank (https://www.ukbiobank.ac.uk/). The phenotype data from UK Biobank participants are available from UK Biobank using their normal access procedures.

The genotype data from the vast majority of the remaining 7,438 NBR participants have been deposited in the European Genome-phenome Archive (EGA) at the EMBL European Bioinformatics Institute. Deposition of genotype at EGA is grouped by rare disease domain: EGA accession codes: BPD: EGAD00001004519, CSVD: EGAD00001004513, EDS: EGAD00001005123, HCM: EGAD00001004514, ICP: EGAD00001004515, IRD: EGAD00001004520, LHON: EGAD00001005122, MPMT: EGAD00001004521, NDD: EGAD00001004522, NPD: EGAD00001004516, PAH: EGAD00001004525, PID: EGAD00001004523, PMG: EGAD00001004517, SMD: EGAD00001004524, SRNS: EGAD00001004518. Access to genotype data provided by the EGA is overseen by a Data Access Committee (DAC) (https://www.ebi.ac.uk/ega/). Access to all NBR WGS data and detailed phenotype data on the 7,438 NBR participants can be requested by completing the NBR Data Access Agreement application.

The ATAC-seq and H3K27ac ChIP-seq data to support the generation of the regulomes are available from GEO or EGA, or referenced to their publication as follows. H3K27ac ChIP-seq: aCD4^78^, B (ERR1043004, ERR1043129, ERR928206, ERR769436), EB (EGAD00001002377), MK (EGAD00001002362), MON (ERR829362 (ERS257420), ERR829412 (ERS222466), ERR493634 (ERS214696)), rCD4^78^. ATAC-seq: aCD4 (GSE124867), B (SRR2126769 (GSE71338)), EB (SRR5489430 (GSM2594182)), MK (EGAD00001001871), MON (accession number requested), rCD4 (GEO accession will be available before publication).

MDT-reported alleles and their clinical interpretation have been deposited in ClinVar (under the name “NIHR Bioresource Rare Diseases”) and DECIPHER.

## Author Contributions

**Corresponding author:** Willem H Ouwehand^1,2,3,4,5^

**Writing Group:** William J Astle^4,6^, Kathleen Freson^7^, Karyn Megy^1,2^, F Lucy Raymond^2,8^, Willem H Ouwehand^1,2,3,4,5^, Kathleen E Stirrups^1,2^, Ernest Turro^1,2,6^

**NIHR BioResource Principal Investigators:** Timothy J Aitman^9,10^, David L Bennett^11^, Mark J Caulfield^12,13^, Patrick F Chinnery^2,14,15^, Peter H Dixon^16^, Kathleen Freson^7^, Daniel P Gale^17^, Ania Koziell^18,19^, Taco W Kuijpers^20,21^, Michael A Laffan^22,23^, Eamonn R Maher^8,24^, Hugh S Markus^25^, Nicholas W Morrell^2,26^, Irene Roberts^27,28,29^, Kenneth G C Smith^26^, Adrian J Thrasher^30^, Hugh Watkins^31,32,33^, Catherine Williamson^16,34^, Christopher Geoffrey Woods^8,35^, F Lucy Raymond^2,8^, Willem H Ouwehand^1,2,3,4,5^

**Ethics, Governance, Recruitment Coordination and Clinical Bioinformatics:** Matthew Brown^1,2^, Naomi Clements Brod^1,2^, John Davis^1,2^, Eleanor F Dewhurst^1,2^, Marie Erwood^1,2^, Amy J Frary^1,2^, Rachel Linger^2,36^, Jennifer Martin^2,26,36^, Sofia Papadia^2,36^, Crina Samarghitean^1,2^, Emily Staples^26^, Catherine Titterton^1,2^, Julie von Ziegenweidt^1,2^, Neil Walker^1,2^, Katherine Yates^1,2,26^, Ping Yu^1,2^, Hannah Stark^2,36^, Roger James^1,2^, Sofie Ashford^2,36^

**Sample and Data processing: Congenica** Eugene Bragin^37^, Calvin Cheah^37^, Radhika Prathalingam^37^, Anthony Rogers^37^, Charles Steward^37^, Katie Tate^37^, Nick Lench^37^; **EMBL-European Bioinformatics Institute** Jeff Almeida-King^38^, Fiona Cunningham^38^, Aoife McMahon^38^, Glen Threadgold^38^, Joannella Morales^38^; **GENALICE** Jack Findhammer^39^, Tim Karten^39^, Bas Tolhuis^39^, Maarten Vandekuilen^39^, Johannes Karten^39^; **High Performance**

**Computing Facility, University of Cambridge** Robert Klima^40^, Ignacio Medina Castello^40^, Stuart Rankin^40^, Wojciech Turek^40^, Paul Calleja^40^; **Illumina** Christian J Bourne^41^, Camilla Colombo^41^, Claire Geoghegan^41^, Terence S A Gerighty^41^, Russell J Grocock^41^, Joseph Hughes^41^, Sarah Hunter^2,41^, John Peden^41^, Christine Rees^41^, Sean Humphray^41^, David R Bentley^41^; **University of Cambridge** Anthony Attwood^1,2^, Abigail Crisp-Hihn^1,2^, Sri V V Deevi^1,2^, Karen Edwards^1,2^, James Fox^1,2^, Fengyuan Hu^1,2^, Jennifer Jolley^1,2^, Rutendo Mapeta^1,2^, Stuart Meacham^1,2^, Paula J Rayner-Matthews^1,2^, Olga Shamardina^1,2^, Ilenia Simeoni^1,2^, Simon Staines^1,2^, Jonathan Stephens^1,2^, Paul Treadaway^1,2^, Salih Tuna^1,2^, Christopher Watt^1,2^, Deborah Whitehorn^1,2^, Yvette Wood^1,2^, Christopher J Penkett^1,2^, Kathleen E Stirrups^1,2^

**Software Development: High Performance Computing Facility, University of Cambridge** Stuart Rankin^40^; **Medical Research Council (MRC) Biostatistics Unit** Sylvia Richardson^6^; **University of Cambridge** Keren Carss^1,2^, Daniel Greene^1,2,6^, Matthias Haimel^1,2,26^, Christopher J Penkett^1,2^, Alba Sanchis-Juan^1,2^, Olga Shamardina^1,2^, Tobias Tilly^1,2^, Salih Tuna^1,2^, Eliska Zlamalova^1^, Ernest Turro^1,2,6^, Stefan Gräf^1,2,26^

**Data Analysis:** William J Astle^4,6^, Christian Babbs^27,29^, Agnieszka Bierzynska^42^, Marta Bleda^26^, Oliver S Burren^26^, Peter H Dixon^16^, Courtney E French^43^, Daniel Greene^1,2,6^, Charaka Hadinnapola^26^, Matthias Haimel^1,2,26^, Adam P Levine^17^, Eleni Louka^27,29^, Adam J Mead^27^, Karyn Megy^1,2^, Monika Mozere^17^, Jennifer O’Sullivan^44^, Steven Okoli^27,29^, David Parry^10^, Beth Psaila^27,29,45^, Anupama Rao^46^, Omid Sadeghi-Alavijeh^17^, Alba Sanchis-Juan^1,2^, Katherine R Smith^12^, Emilia M Swietlik^26^, Rhea Y Y Tan^25^, Natalie van Zuydam^11^, Wei Wei^14,15^, James Whitworth^8,24,47^, Eliska Zlamalova^1^, Augusto Rendon^1,12^, Keren Carss^1,2^, Stefan Gräf^1,2,26^, Hana Lango Allen^1,2^, Ernest Turro^1,2,6^

**Clinical Interpretation and Multi-Disciplinary Teams:** Stephen Abbs^48^, Timothy J Aitman^9,10^, Philip Ancliff^46^, Gavin Arno^49,50^, Chiara Bacchelli^30^, David L Bennett^11^, Agnieszka Bierzynska^42^, Iulia Blesneac^11^, Siobhan O Burns^51,52^, Keren Carss^1,2^, Louise C Daugherty^1,2,12^, Sri V V Deevi^1,2^, Peter H Dixon^16^, Kate Downes^1,2^, Anna M Drazyk^25^, Daniel Duarte^1,2^, Courtney E French^43^, Kathleen Freson^7^, Daniel P Gale^17^, Kimberly C Gilmour^30,46^, Keith Gomez^53,54^, Detelina Grozeva^8^, Charaka Hadinnapola^26^, Simon Holden^55^, Ania Koziell^18,19^, Taco W Kuijpers^20,21^, Dinakantha Kumararatne^56^, Michael A Laffan^22,23^, Hana Lango Allen^1,2^, D Mark Layton^22,23^, Adam P Levine^17^, Eleni Louka^27,29^, Eamonn R Maher^8,24^, Jesmeen Maimaris^30^, Rutendo Mapeta^1,2^, Hugh S Markus^25^, Jennifer Martin^2,26,36^, Sarju Mehta^55^, Nicholas W Morrell^2,26^, Andrew D Mumford^57,58^, David Parry^10^, Irene Roberts^27,28,29^, Noemi B Roy^27,28,29^, Moin A Saleem^42,59^, Alba Sanchis-Juan^1,2^, Sinisa Savic^60,61^, Ilenia Simeoni^1,2^, Emily Staples^26^, Emilia M Swietlik^26^, Rhea Y Y Tan^25^, James E Thaventhiran^62^, Andreas C Themistocleous^11^, David Thomas^26^, Marc Tischkowitz^48,63^, Matthew Traylor^25^, Ernest Turro^1,2,6^, Natalie van Zuydam^11^, Anthony M Vandersteen^64^, Andrew R Webster^49,50^, James Whitworth^8,24,47^, Catherine Williamson^16,34^, Christopher Geoffrey Woods^8,35^, Willem H Ouwehand^1,2,3,4,5^, F Lucy Raymond^2,8^, Stefan Gräf^1,2,26^, Karyn Megy^1,2^

**Non-coding Space Analysis Group: University of Cambridge** Oliver S Burren^26^, Stefan Gräf^1,2,26^, Luigi Grassi^1,2^, Daniel Greene^1,2,6^, Myrto Kostadima^1^, Roman Kreuzhuber^1,2^, Hana Lango Allen^1,2^, Romina Petersen^1,2^, Denis Seyres^1,2^, James E Thaventhiran^62^, Mattia Frontini^1,2,5^, Ernest Turro^1,2,6^; **University of Oxford** Anthony J Cutler^65^, John A Todd^65^; **Wellcome Sanger Institute** Patrick J Short^3^, Matthew Hurles^3^

**Functional Analysis Group:** Nichola Cooper^66^, Nicholas S Gleadall^1,2^, Andrew D Mumford^57,58^, Helen Oram^67^, Alba Sanchis-Juan^1,2^, Olga Shamardina^1,2^, Jonathan Stephens^1,2^, Patrick Thomas^1,2^, Chantal Thys^7^, Sarah K Westbury^57,58^, Suthesh Sivapalaratnam^4,68,69,70^, Kate Downes^1,2^, Kathleen Freson^7^

**Data Visualisation:** Salih Tuna^1,2^, William J Astle^4,6^, Sri V V Deevi^1,2^, Stefan Gräf^1,2,26^, Daniel Greene^1,2,6^, Matthias Haimel^1,2,26^, Hana Lango Allen^1,2^, Karyn Megy^1,2^, Christopher J Penkett^1,2^, Alba Sanchis-Juan^1,2^, Olga Shamardina^1,2^, Kathleen E Stirrups^1,2^, Ernest Turro^1,2,6^

**Steering groups: NIHR BioResource Sequencing and Informatics Committee (SIC)** Gerome Breen^71,72^, John Chambers^73,74,75,76,77^, Matthew Hurles^3^, Nathalie Kingston^2^, Mark I McCarthy^29,32,78^, Nilesh Samani^79^, Michael Simpson^80^, Nicholas Wood^81,82^, Willem H Ouwehand^1,2,3,4,5^, F Lucy Raymond^2,8^; **NIHR BioResource – Rare Diseases Senior Management Team (SMT)** Sofie Ashford^2,36^, Debra Fletcher^1,2^, Mary A Kasanicki^35^, Christopher J Penkett^1,2^, Hannah Stark^2,36^, Kathleen E Stirrups^1,2^, Neil Walker^1,2^, Timothy Young^1,2^, Roger James^1,2^, Nathalie Kingston^2^, F Lucy Raymond^2,8^, John R Bradley^2,24,26,35,83^, Willem H _Ouwehand_1,2,3,4,5

**NIHR BioResource-Rare Diseases Study teams: Bleeding, thrombotic and Platelet Disorders (BPD)** Tadbir Bariana^53,54^, Claire Lentaigne^22,23^, Suthesh Sivapalaratnam^4,68,69,70^, Sarah K Westbury^57,58^, David J Allsup^84,85^, Tamam Bakchoul^86^, Tina Biss^87^, Sara Boyce^88^, Janine Collins^1,68^, Peter W Collins^89^, Nicola S Curry^90^, Kate Downes^1,2^, Tina Dutt^91^, Wendy N Erber^92^, Gillian Evans^93^, Tamara Everington^94,95^, Remi Favier^96,97^, Keith Gomez^53,54^, Daniel Greene^1,2,6^, Andreas Greinacher^98^, Paolo Gresele^99^, Daniel Hart^68^, Rashid Kazmi^88^, Anne M Kelly^35^, Michele Lambert^100,101^, Bella Madan^44^, Sarah Mangles^95^, Mary Mathias^102^, Carolyn Millar^22,23^, Paquita Nurden^103^, Samya Obaji^104^, Kathelijne Peerlinck^7^, Catherine Roughley^93^, Sol Schulman^105^, Marie Scully^106^, Susan E Shapiro^90^, Keith Sibson^102^, Ilenia Simeoni^1,2^, Matthew C Sims^1,107^, R Campbell Tait^108^, Kate Talks^87^, Chantal Thys^7^, Cheng-Hock Toh^91^, Chris Van Geet^7^, John-Paul Westwood^106^, Sofia Papadia^2,36^, Ernest Turro^1,2,6^, Andrew D Mumford^57,58^, Willem H Ouwehand^1,2,3,4,5^, Kathleen Freson^7^, Michael A Laffan^22,23^; **Cerebral Small Vessel Disease (CSVD)** Rhea Y Y Tan^25^, Julian Barwell^109,110^, Kate Downes^1,2^, Kirsty Harkness^111^, Sarju Mehta^55^, Keith W Muir^112^, Ahamad Hassan^113^, Matthew Traylor^25^, Anna M Drazyk^25^, Stefan Gräf^1,2,26^, Hugh S Markus^25^; **Ehlers Danlos Syndrome (EDS)** David Parry^10^, Munaza Ahmed^114^, Alex Henderson^115^, Hanadi Kazkaz^106^, Anthony M Vandersteen^64^, Timothy J Aitman^9,10^; **Hypertrophic Cardiomyopathy (HCM)** Elizabeth Ormondroyd^31,33^, Kate Thomson^31,33^, Timothy Dent^33^, Paul Brennan^115,116,117^, Rachel J Buchan^118,119^, Teofila Bueser^18,120,121^, Gerald Carr-White^122^, Stuart Cook^118,123,124,125^, Matthew J Daniels^31,33,126^, Andrew R Harper^31,32^, Alex Henderson^115^, James S Ware^118,119,123^, Hugh Watkins^31,32,33^; **Intrahepatic Cholestasis of Pregnancy (ICP)** Peter H Dixon^16^, Jenny Chambers^16,127^, Floria Cheng^127^, Maria C Estiu^128^, William M Hague^129^, Hanns-Ulrich Marschall^130^, Marta Vazquez-Lopez^127^, Catherine Williamson^16,34^; **Inherited Retinal Disorders (IRD)** Gavin Arno^49,50^, Eleanor F Dewhurst^1,2^, Marie Erwood^1,2^, Courtney E French^43^, Michel Michaelides^49,50^, Anthony T Moore^49,50,131^, Alba Sanchis-Juan^1,2^, Keren Carss^1,2^, Andrew R Webster^49,50^, F Lucy Raymond^2,8^; **Leber Hereditary Optic Neuropathy (LHON)** Patrick F Chinnery^2,14,15^, Philip Griffiths^132,133^, Rita Horvath^134,135^, Gavin Hudson^134^, Neringa Jurkute^49,54^, Angela Pyle^134^, Wei Wei^14,15^, Patrick Yu-Wai-Man^14,15,136^; **Multiple Primary Malignant Tumours (MPMT)** James Whitworth^8,24,47^, Julian Adlard^137^, Munaza Ahmed^114^, Ruth Armstrong^8,24,47^, Julian Barwell^109,110^, Carole Brewer^138^, Ruth Casey^8,24,47^, Trevor R P Cole^139^, Dafydd Gareth Evans^140^, Lynn Greenhalgh^141^, Helen L Hanson^142^, Alex Henderson^115^, Jonathan Hoffman^139^, Louise Izatt^143^, Ajith Kumar^114^, Fiona Lalloo^144^, Kai Ren Ong^139^, Soo-Mi Park^24,47,48^, Joan Paterson^8,24,47^, Claire Searle^145^, Lucy Side^146^, Katie Snape^142^, Emma Woodward^144^, Marc Tischkowitz^48,63^, Eamonn R Maher^8,24^; **Neurological and Developmental Disorders (NDD)** Keren Carss^1,2^, Eleanor F Dewhurst^1,2^, Marie Erwood^1,2^, Courtney E French^43^, Detelina Grozeva^8^, Alba Sanchis-Juan^1,2^, Manju A Kurian^147,148^, F Lucy Raymond^2,8^; **Neuropathic Pain Disorders (NPD)** Andreas C Themistocleous^11^, Iulia Blesneac^11^, David Gosal^149^, Rita Horvath^134,135^, Andrew Marshall^150,151,152^, Emma Matthews^153,154^, Mark I McCarthy^29,32,78^, Tara Renton^121^, Andrew S C Rice^155,156^, Tom Vale^11^, Natalie van Zuydam^11^, Suellen M Walker^30,46^, Christopher Geoffrey Woods^8,35^, David L Bennett^11^; **Primary Immune Disorders (PID)** James E Thaventhiran^62^, Hana Lango Allen^1,2^, Siobhan O Burns^51,52^, Sinisa Savic^60,61^, Oliver S Burren^26^, Hana Alachkar^149^, Richard Antrobus^157^, Helen E Baxendale^26,43,158,159^, Michael J Browning^160^, Matthew S Buckland^161^, Nichola Cooper^66^, Elizabeth Drewe^162^, J David M Edgar^163,164^, William Egner^165^, Kimberly C Gilmour^30,46^, Sarah Goddard^166^, Pavels Gordins^167^, Sofia Grigoriadou^168^, Scott Hackett^169^, Rosie Hague^170^, Grant Hayman^171^, Archana Herwadkar^149^, Aarnoud P Huissoon^169^, Stephen Jolles^172^, Peter Kelleher^173,174^, Dinakantha Kumararatne^56^, Hilary Longhurst^168^, Lorena E Lorenzo^168^, Paul A Lyons^26^, Jesmeen Maimaris^30^, Sadia Noorani^175^, Alex Richter^157^, Crina Samarghitean^1,2^, Ravishankar B Sargur^165^, W A Carrock Sewell^176^, Ilenia Simeoni^1,2^, Emily Staples^26^, David Thomas^26^, Moira J Thomas^177,178^, Steven B Welch^179^, Austen Worth^46^, Patrick F K Yong^180^, Taco W Kuijpers^20,21^, Adrian J Thrasher^30^, Kenneth G C Smith^26^; **Primary Membranoproliferative Glomerulonephritis (PMG)** Adam P Levine^17^, Melanie M Y Chan^17^, Omid Sadeghi-Alavijeh^17^, Edwin K S Wong^117,181^, H Terence Cook^182^, Martin T Christian^183^, Matthew Hall^162^, Claire Harris^181^, Paul McAlinden^181^, Kevin J Marchbank^181,184^, Stephen Marks^46^, Heather Maxwell^170^, Monika Mozere^17^, Julie Wessels^166^, MPGN/C3 Glomerulopathy Rare Renal Disease group^185^, Sally A Johnson^181,186^, Daniel P Gale^17^; **Pulmonary Arterial Hypertension (PAH)** Marta Bleda^26^, Charaka Hadinnapola^26^, Matthias Haimel^1,2,26^, Emilia M Swietlik^26^, Harm Bogaard^187^, Colin Church^188^, Gerry Coghlan^161^, Robin Condliffe^189^, Paul Corris^181,190^, Cesare Danesino^191^, Mélanie Eyries^192^, Henning Gall^193^, Stefano Ghio^194^, Hossein-Ardeschir Ghofrani^66,193^, J Simon R Gibbs^118^, Barbara Girerd^195,196,197^, Simon Holden^55^, Arjan Houweling^187^, Luke S Howard^118,198^, Marc Humbert^195,196,197^, David G Kiely^189^, Gabor Kovacs^199,200^, Allan Lawrie^201^, Robert V MacKenzie Ross^202^, Jennifer Martin^2,26,36^, Shahin Moledina^46^, David Montani^195,196,197^, Michael Newnham^26,159^, Andrea Olschewski^199^, Horst Olschewski^199,200^, Andrew Peacock^188^, Joanna Pepke-Zaba^159^, Christopher J Rhodes^66^, Laura Scelsi^194^, Werner Seeger^193^, Nicole Soranzo^1,3^, Florent Soubrier^192^, Jay Suntharalingam^202^, Mark Toshner^26,159^, Carmen Treacy^26,159^, Richard Trembath^18^, Anton Vonk Noordegraaf^187^, Quinten Waisfisz^203^, John Wharton^66^, Martin R Wilkins^66^, Stephen J Wort^119,204^, Katherine Yates^1,2,26^, Stefan Gräf^1,2,26^, Nicholas W Morrell^2,26^; **Stem cell and Myeloid Disorders (SMD)** Eleni Louka^27,29^, Noemi B Roy^27,28,29^, Anupama Rao^46^, Philip Ancliff^46^, Christian Babbs^27,29^, D Mark Layton^22,23^, Adam J Mead^27^, Jennifer O’Sullivan^44^, Steven Okoli^27,29^, Irene Roberts^27,28,29^; **Steroid Resistant Nephrotic Syndrome (SRNS)** Moin A Saleem^42,59^, Agnieszka Bierzynska^42^, Carmen Bugarin Diz^18^, Elizabeth Colby^42^, Melanie N Ekani^122^, Simon Satchell^42,205^, Ania Koziell^18,19^; **UK Biobank Extreme Red Blood Cell Traits (UKB)** William J Astle^4,6^, Suthesh Sivapalaratnam^4,68,69,70^, Noemi B Roy^27,28,29^

**Provision of WES data for coverage analysis: The INTERVAL Study** Klaudia Walter^3^, Nicole Soranzo^1,3^; **The Columbia University exome sequencing study for chronic kidney disease** Ali G Gharavi^206^, David B Goldstein^207^

**Genomics England Rare Diseases Pilot Study (GEL RD Pilot): Genomics England Core Teams** Tom Fowler^12^, Chris Odhams^12^, Richard Scott^12,46^, Damian Smedley^12,13^, Katherine R Smith^12^, Alex Stuckey^12^, Ellen Thomas^12,122^, Augusto Rendon^1,12^, Mark J Caulfield^12,13^; **Cambridge University Hospitals NHS Foundation Trust** Stephen Abbs^48^, Nigel Burrows^35^, Manali Chitre^43^, Eleanor F Dewhurst^1,2^, R Andres Floto^26,35,159^, Michael Gattens^35^, Mark Gurnell^26,35^, Simon Holden^55^, Wilf Kelsall^35^, Sarju Mehta^55^, Ken E S Poole^26,35^, Robert Ross-Russell^35^, Olivera Spasic-Boskovic^48^, Philip Twiss^48^, Annette Wagner^35^, F Lucy Raymond^2,8^; **Central Manchester University Hospitals NHS Trust and Manchester University** Siddharth Banka^144,208^, Graeme C Black^144,208^, Jill Clayton-Smith^144,208^, Sofia Douzgou^144,208^, William G Newman^144,208^; **Great Ormond Street Hospital for Children NHS Foundation Trust and University College London** Lara Abulhoul^46^, Paul Aurora^46^, Detlef Bockenhauer^46^, Maureen Cleary^46^, Mehul Dattani^209,210^, Vijeya Ganesan^46^, Clarissa Pilkington^46^, Shamima Rahman^46,209^, Neil Shah^30,46^, Lucy Wedderburn^30,211,212^, Maria A K Bitner-Glindzicz^46,209^; **Guy’s and St Thomas’ Hospital NHS Foundation Trust and King’s College London** Teofila Bueser^120,213^, Cecilia J Compton^120^, Charu Deshpande^120^, Hiva Fassihi^214^, Eshika Haque^120^, Louise Izatt^120^, Dragana Josifova^120^, Shehla N Mohammed^120^, Leema Robert^120^, Sarah J Rose^120^, Deborah M Ruddy^120^, Robert N Sarkany^214^, Genevieve Sayer^120^, Adam C Shaw^120^, Melita Irving^120^, Frances A Flinter^120^; **Moorfields Eye Hospital NHS Trust and University College London** Gavin Arno^49,50^, Samantha Malka^49,50^, Michel Michaelides^49,50^, Anthony T Moore^49,50,131^, Andrew R Webster^49,50^; **Oxford University Hospitals NHS Trust and the University of Oxford** Carolyn Campbell^215^, Kate Gibson^215^, Nils Koelling^216^, Tracy Lester^215^, Andrea H Nemeth^11,217^, Claire Palles^218^, Smita Patel^219^, Noemi B Roy^216,220^, Arjune Sen^29,221,222^, John M Taylor^215^, Ian P Tomlinson^218^, Jenny C Taylor^29,32^, Andrew O Wilkie^216^; **Newcastle upon Tyne Hospitals NHS Foundation Trust and Newcastle University** Paul Brennan^115,116,117^, Andrew C Browning^223^, John Burn^115^, Patrick F Chinnery^2,14,15^, Anthony De Soyza^117,181,224^, Jodie Graham^225^, Rita Horvath^132^, Simon Pearce^117,225^, Richard Quinton^117,134^, Andrew M Schaefer^117,132^, Brian T Wilson^114,117,134^, Michael Wright^115^, Patrick Yu-Wai-Man^14,15,136^, John A Sayer^117,134^; **University College London Hospitals NHS Trust and University College London** Michael Simpson^80^, Petros Syrris^226^, Perry Elliott^226,227^, Henry Houlden^81^, Phil L Beales^46,209^

## Acknowledgements

This research was made possible through access to the data and findings generated by two pilot studies for the 100,000 Genomes Project. The enrolment was coordinated for one by the NIHR BioResource and for the other by Genomics England Limited (GEL), a wholly owned company of the Department of Health in the UK. These pilot studies were mainly funded by grants from the National Institute for Health Research (NIHR) in England to the University of Cambridge and GEL, respectively. Additional funding was provided by the BHF, MRC, NHS England, the Wellcome Trust and many other fund providers (also see Funding acknowledgment for individual researchers). The pilot studies use data provided by patients and their close relatives and collected by the NHS and other healthcare providers as part of their care and support. The vast majority of participants in the two pilot studies have been enrolled in the NIHR BioResource. We thank all volunteers for their participation, and also gratefully acknowledge NIHR Biomedical Research Centres, NIHR BioResource Centres, NHS Trust Hospitals, NHS Blood and Transplant and staff for their contribution. This research has been conducted using the UK Biobank Resource under Application Number 9616, granting access to DNA samples and accompanying participant data. UK Biobank has received funding from the MRC, Wellcome Trust, Department of Health, British Heart Foundation (BHF), Diabetes UK, Northwest Regional Development Agency, Scottish Government, and Welsh Assembly Government. The MRC and Wellcome Trust played a key role in the decision to establish UK Biobank.

**Funding acknowledgment for individual researchers** AMM and JMo are funded by The Wellcome Trust (WT200990/Z/16/Z) and the European Molecular Biology Laboratory; KGCS holds a Wellcome Investigator Award, MRC Programme Grant (number MR/L019027/1); MIM is a Wellcome Senior Investigator and receives support from the Wellcome Trust (090532, 0938381) and is a member of the DOLORisk consortium funded by the European Commission Horizon 2020 (ID633491); RHo is a Wellcome Trust Investigator (109915/Z/15/Z), who receives support from the Wellcome Centre for Mitochondrial Research (203105/Z/16/Z), MRC (MR/N025431/1), the European Research Council (309548), the Wellcome Trust Pathfinder Scheme (201064/Z/16/Z), the Newton Fund (UK/Turkey, MR/N027302/1) and the European Union H2020 – Research and Innovation Actions (SC1-PM-03-2017, Solve-RD); DLB is a Wellcome clinical scientist (202747/Z/16/Z) and is a member of the DOLORisk consortium funded by the European Commission Horizon 2020 (ID633491); JSW is funded by Wellcome Trust [107469/Z/15/Z], NIHR Cardiovascular Biomedical Research Unit at Royal Brompton & Harefield NHS Foundation Trust and Imperial College London; AJT is supported by the Wellcome Trust (104807/Z/14/Z) and the NIHR Biomedical Research Centre at Great Ormond Street Hospital for Children NHS Foundation Trust and University College London; LSo is supported by the Wellcome Trust Institutional Strategic Support Fund (204809/Z/16/Z) awarded to St. George’s, University of London; MJD receives funding from Wellcome Trust (WT098519MA); MCS holds an MRC Clinical Research Training Fellowship (MR/R002363/1); JAS is funded by MRC UK grant MR/M012212/1; AJM received funding from an MRC Senior Clinical Fellowship (MR/L006340/1); CLe received funding from an MRC Clinical Research Training Fellowship (MR/J011711/1); MRW holds a NIHR award to the NIHR Imperial Clinical Research Facility at Imperial College Healthcare NHS Trust; DJW receives part of his salary from the NIHR University College London Hospitals Biomedical Research Centre; MAKu holds a NIHR Research Professorship (NIHR-RP-2016-07-019) and Wellcome Intermediate Fellowship (098524/Z/12/A); MJC is an NIHR Senior Investigator and is funded by the NIHR Barts Biomedical Research Centre; NCo is partially funded by NIHR Imperial College Biomedical Research Centre; CHad was funded through a PhD Fellowship by the NIHR Translational Research Collaboration - Rare Diseases; ADM and SKW were funded by the NIHR Bristol Biomedical Research Centre; ELM received funding from the NIHR Biomedical Research Centre at University College London Hospitals; KCG received funding from the NIHR Great Ormond Street Biomedical Research Centre; IR and ELo are supported by the NIHR Translational Research Collaboration - Rare Diseases; JCT, JMT and SPat are funded by the NIHR Oxford Biomedical Research Centre; GArn is funded by the NIHR Moorfields Biomedical Research Centre and UCL Institute of Ophthalmology, Fight for Sight (UK) Early Career Investigator Award, Moorfields Eye Hospital Special Trustees, Moorfields Eye Charity, Foundation Fighting Blindness (USA) and Retinitis Pigmentosa Fighting Blindness; ATM is funded by Retinitis Pigmentosa Fighting Blindness, PY-W-M is supported by grants from MRC UK (G1002570), Fight for Sight (1570/1571), Fight for Sight (24TP171), NIHR (IS-BRC-1215-20002); SOB is supported by NIHR Translational Research Collaboration - Rare Diseases (01/04/15-30/04/2017); ARW works for the NIHR Moorfields Biomedical Research Centre and the UCL Institute of Ophthalmology and Moorfields Eye Hospital; the following NIHR Biomedical Research Centres contributed to the enrolment for the ICP domain: Imperial College Healthcare NHS Trust, Guy’s and St Thomas’ NHS Foundation Trust and King’s College London. All authors affiliated with Moorfields Eye hospital and Institute of Ophthalmology are funded by the NIHR Biomedical Resource Centre at UCL Institute of Ophthalmology and Moorfields; ACT is a member of the International Diabetic Neuropathy Consortium, the Novo Nordisk Foundation (Ref. NNF14SA0006) and is a member of the DOLORisk consortium funded by the European Commission Horizon 2020 (ID633491); JWhi is a recipient of a Cancer Research UK Cambridge Cancer Centre Clinical Research Training Fellowship; PSh holds a Henry Smith Charity and Department of Health (UK) Senior Fellowship; SAJ is funded by Kids Kidney Research; DPG is funded by the MRC, Kidney Research UK and St Peters Trust for Kidney, Bladder and Prostate Research; KJM is supported by the Northern Counties Kidney Research Fund; PHD receives funding from ICP Support; TKB received a PhD fellowship from the NHSBT and British Society of Haematology; HSM receives support from BHF Programme Grant no. RG/16/4/32218; AL is a BHF Senior Basic Science Research Fellow - FS/13/48/30453; KF and CVG are supported by the Research Council of the University of Leuven (BOF KU Leuven, Belgium; OT/14/098); HJB works for the Netherlands CardioVascular Research Initiative (CVON); GBa holds a WA Department of Health, Raine Clinician Research Fellowship 2015GB.

## Disclaimer

The views expressed are those of the author(s) and not necessarily those of the NHS, the NIHR, the Department of Health and Social Care or any of the other funding agencies.

## Additional information

**Extended data** is available for this paper at TBC

**Supplementary information** is available for this paper at TBC

**Reprints and permissions information** is available at http://www.nature.com/reprints. **Correspondence and requests for materials** should be addressed to. **Publisher’s note** Springer Nature remains neutral with regard to jurisdictional claims in published maps and institutional affiliations.

## APPENDIX – Affiliations

^1^Department of Haematology, University of Cambridge, Cambridge Biomedical Campus, Cambridge, UK. ^2^NIHR BioResource, Cambridge University Hospitals NHS Foundation, Cambridge Biomedical Campus, Cambridge, UK. ^3^Wellcome Sanger Institute, Wellcome Genome Campus, Hinxton, Cambridge, UK. ^4^NHS Blood and Transplant, Cambridge Biomedical Campus, Cambridge, UK. ^5^British Heart Foundation Cambridge Centre of Excellence, University of Cambridge, Cambridge, UK. ^6^MRC Biostatistics Unit, Cambridge Institute of Public Health, University of Cambridge, Cambridge, UK. ^7^Department of Cardiovascular Sciences, Center for Molecular and Vascular Biology, KULeuven, Leuven, Belgium. ^8^Department of Medical Genetics, Cambridge Institute for Medical Research, University of Cambridge, Cambridge Biomedical Campus, Cambridge, UK. ^9^MRC Clinical Sciences Centre, Faculty of Medicine, Imperial College London, London, UK. ^10^Institute of Genetics and Molecular Medicine, University of Edinburgh, Edinburgh, UK. ^11^The Nuffield Department of Clinical Neurosciences, University of Oxford, John Radcliffe Hospital, Oxford, UK. ^12^Genomics England, Charterhouse Square, London, UK. ^13^William Harvey Research Institute, NIHR Biomedical Research Centre at Barts, Queen Mary University of London, London, UK. ^14^Department of Clinical Neurosciences, School of Clinical Medicine, University of Cambridge, Cambridge Biomedical Campus, Cambridge, UK. ^15^Medical Research Council Mitochondrial Biology Unit, Cambridge Biomedical Campus, Cambridge, UK. ^16^Women and Children’s Health, School of Life Course Sciences, King’s College London, London, UK. ^17^UCL Centre for Nephrology, University College London, London, UK. ^18^King’s College London, London, UK. ^19^Department of Paediatric Nephrology, Evelina London Children’s Hospital, Guy’s & St Thomas’ NHS Foundation Trust, London, UK. ^20^Department of Pediatric Hematology, Immunology, Rheumatology and Infectious Diseases, Emma Children’s Hospital, Academic Medical Center (AMC), University of Amsterdam, Amsterdam, The Netherlands. ^21^Department of Blood Cell Research, Sanquin, Amsterdam, The Netherlands. ^22^Department of Haematology, Hammersmith Hospital, Imperial College Healthcare NHS Trust, London, UK. ^23^Centre for Haematology, Imperial College London, London, UK. ^24^NIHR Cambridge Biomedical Research Centre, Cambridge Biomedical Campus, Cambridge, UK. ^25^Stroke Research Group, Department of Clinical Neurosciences, University of Cambridge, Cambridge Biomedical Campus, Cambridge, UK. ^26^Department of Medicine, School of Clinical Medicine, University of Cambridge, Cambridge Biomedical Campus, Cambridge, UK. ^27^MRC Molecular Haematology Unit, MRC Weatherall Institute of Molecular Medicine, University of Oxford, Oxford, UK. ^28^Department of Paediatrics, Weatherall Institute of Molecular Medicine, University of Oxford, Oxford, UK. ^29^NIHR Oxford Biomedical Research Centre, Oxford University Hospitals Trust, Oxford, UK. ^30^UCL Great Ormond Street Institute of Child Health, London, UK. ^31^Department of Cardiovascular Medicine, Radcliffe Department of Medicine, University of Oxford, Oxford, UK. ^32^Wellcome Centre for Human Genetics, University of Oxford, Oxford, UK. ^33^Oxford University Hospitals NHS Foundation Trust, Oxford, UK. ^34^Institute of Reproductive and Developmental Biology, Surgery and Cancer, Hammersmith Hospital, Imperial College Healthcare NHS Trust, London, UK. ^35^Addenbrookes Hospital, Cambridge University Hospitals NHS Foundation Trust, Cambridge, UK. ^36^Department of Public Health and Primary Care, University of Cambridge, Cambridge, UK. ^37^Congenica, Biodata Innovation Centre, Wellcome Genome Campus, Hinxton, Cambridge, UK. ^38^European Molecular Biology Laboratory, European Bioinformatics Institute (EMBL-EBI), Wellcome Genome Campus, Hinxton, Cambridge, UK. ^39^GENALICE BV, Harderwijk, The Netherlands. ^40^High Performance Computing Service, University of Cambridge, Cambridge, UK. ^41^Illumina Cambridge Limited, Chesterford Research Park, Little Chesterford, Saffron Walden, Essex, UK. ^42^Bristol Renal and Children’s Renal Unit, Bristol Medical School, University of Bristol, Bristol, UK. ^43^Department of Paediatrics, School of Clinical Medicine, University of Cambridge, Cambridge Biomedical Campus, Cambridge, UK. ^44^Department of Haematology, Guy’s and St Thomas’ NHS Foundation Trust, London, UK. ^45^Centre for Haematology, Department of Medicine, Hammersmith Hospital, Imperial College Healthcare NHS Trust, London, UK. ^46^Great Ormond Street Hospital for Children NHS Foundation Trust, London, UK. ^47^Cancer Research UK Cambridge Centre, Cambridge Biomedical Campus, Cambridge, UK. ^48^East Anglian Medical Genetics Service, Cambridge University Hospitals NHS Foundation Trust, Cambridge, UK. ^49^Moorfields Eye Hospital NHS Foundation Trust, London, UK. ^50^UCL Institute of Ophthalmology, University College London, London, UK. ^51^Institute of Immunity and Transplantation, University College London, London, UK. ^52^Department of Immunology, Royal Free London NHS Foundation Trust, London, UK. ^53^The Katharine Dormandy Haemophilia Centre and Thrombosis Unit, Royal Free London NHS Foundation Trust, London, UK. ^54^University College London, London, UK. ^55^Department of Clinical Genetics, Addenbrookes Hospital, Cambridge University Hospitals NHS Foundation Trust, Cambridge, UK. ^56^Department of Clinical Immunology, Addenbrookes Hospital, Cambridge University Hospitals NHS Foundation Trust, Cambridge, UK. ^57^School of Cellular and Molecular Medicine, University of Bristol, Bristol, UK. ^58^University Hospitals Bristol NHS Foundation Trust, Bristol, UK. ^59^Bristol Royal Hospital for Children, University Hospitals Bristol NHS Foundation Trust, Bristol, UK. ^60^The Department of Clinical Immunology and Allergy and The NIHR Leeds Biomedical Research Centre, Leeds, UK. ^61^Leeds Institute of Rheumatic and Musculoskeletal Medicine, St James’s University Hospital, Leeds, UK. ^62^MRC Toxicology Unit, School of Biological Sciences, University of Cambridge, Cambridge, UK. ^63^Department of Medical Genetics and NIHR Cambridge Biomedical Research Centre, University of Cambridge, Cambridge, UK. ^64^Division of Medical Genetics, IWK Health Centre, Dalhousie University, Halifax, Canada. ^65^JDRF/Wellcome Diabetes and Inflammation Laboratory, Wellcome Centre for Human Genetics, Nuffield Department of Medicine, NIHR Oxford Biomedical Research Centre, University of Oxford, Oxford, UK. ^66^Department of Medicine, Imperial College London, London, UK. ^67^Department of Paediatric Haematology, University Hospital Southampton NHS Foundation Trust, Southampton, UK. ^68^The Royal London Hospital, Barts Health NHS Foundation Trust, London, UK. ^69^Department of Haematology, Cambridge University Hospitals NHS Foundation Trust, Cambridge, UK. ^70^Queen Mary University of London, London, UK. ^71^MRC Social, Genetic & Developmental Psychiatry Centre, Institute of Psychiatry, Psychology & Neuroscience, King’s College London, London, UK. ^72^NIHR Biomedical Research Centre for Mental Health, Maudsley Hospital, London, UK. ^73^Lee Kong Chian School of Medicine, Nanyang Technological University, Singapore, Singapore. ^74^Department of Epidemiology and Biostatistics, Imperial College London, London, UK. ^75^Department of Cardiology, Ealing Hospital, Middlesex, UK. ^76^Imperial College Healthcare NHS Trust, London, UK. ^77^MRC-PHE Centre for Environment and Health, Imperial College London, London, UK. ^78^Oxford Centre for Diabetes, Endocrinology and Metabolism, University of Oxford, Churchill Hospital, Oxford, UK. ^79^Department of Cardiovascular Sciences and NIHR Leicester Biomedical Research Research Centre, University of Leicester, Leicester, UK. ^80^Genetics and Molecular Medicine, King’s College London, London, UK. ^81^Department of Molecular Neuroscience, UCL Institute of Neurology, London, UK. ^82^UCL Genetics Institute, London, UK. ^83^Department of Renal Medicine, Addenbrookes Hospital, Cambridge University Hospitals NHS Foundation Trust, Cambridge, UK. ^84^Queens Centre for Haematology and Oncology, Castle Hill Hospital, Hull and East Yorkshire NHS Trust, Cottingham, UK. ^85^Hull York Medical School, University of Hull, Hull, UK. ^86^Center for Clinical Transfusion Medicine, University Hospital of Tübingen, Tübingen, Germany. ^87^Haematology Department, Royal Victoria Infirmary, The Newcastle upon Tyne Hospitals NHS Foundation Trust, Newcastle upon Tyne, UK. ^88^Southampton General Hospital, University Hospital Southampton NHS Foundation Trust, Southampton, UK. ^89^Institute of Infection and Immunity, School of Medicine Cardiff University, Cardiff, UK. ^90^Oxford Haemophilia and Thrombosis Centre, Oxford University Hospitals NHS Trust, Oxford Comprehensive Biomedical Research Centre, Oxford, UK. ^91^The Roald Dahl Haemostasis and Thrombosis Centre, The Royal Liverpool University Hospital, Liverpool, UK. ^92^Medical School and School of Biomedical Sciences, Faculty of Health and Medical Sciences, The University of Western Australia, Crawley, Australia. ^93^Haemophilia Centre, Kent & Canterbury Hospital, East Kent Hospitals University Foundation Trust, Canterbury, UK. ^94^Salisbury District Hospital, Salisbury NHS Foundation Trust, Salisbury, UK. ^95^Haemophilia, Haemostasis and Thrombosis Centre, Hampshire Hospitals NHS Foundation Trust, Basingstoke, UK. ^96^INSERM UMR 1170, Gustave Roussy Cancer Campus, Universite Paris-Saclay, Villejuif, France. ^97^Service d’Hematologie biologique, Centre de Reference des Pathologies Plaquettaires, Hopital Armand Trousseau, Assistance Publique-Hopitaux de Paris, Paris, France. ^98^Institute for Immunology and Transfusion Medicine, University Medicine Greifswald, Greifswald, Germany. ^99^Section of Internal and Cardiovascular Medicine, University of Perugia, Perugia, Italy. ^100^Division of Hematology, The Children’s Hospital of Philadelphia, Philadelphia, USA. ^101^Department of Pediatrics, Perelman School of Medicine at the University of Pennsylvania, Philadelphia, USA. ^102^Department of Haematology, Great Ormond Street Hospital for Children NHS Foundation Trust, London, UK. ^103^Institut Hospitalo-Universitaire de Rythmologie et de Modelisation Cardiaque, Plateforme Technologique d’Innovation Biomedicale, Hopital Xavier Arnozan, Pessac, France. ^104^The Arthur Bloom Haemophilia Centre, University Hospital of Wales, Cardiff, UK. ^105^Beth Israel Deaconess Medical Centre and Harvard Medical School, Boston, USA. ^106^University College London Hospitals NHS Foundation Trust, London, UK. ^107^Oxford Haemophilia and Thrombosis Centre, The Churchill Hospital, Oxford University Hospitals NHS Trust, Oxford, UK. ^108^Glasgow Royal Infirmary, NHS Greater Glasgow and Clyde, Glasgow, UK. ^109^Department of Clinical Genetics, Leicester Royal Infirmary, University Hospitals of Leicester, Leicester, UK. ^110^University of Leicester, Leicester, UK. ^111^Department of Neurology, Sheffield Teaching Hospitals NHS Foundation Trust, Sheffield, UK. ^112^Institute of Neuroscience and Psychology, University of Glasgow, Glasgow, UK. ^113^Department of Neurology, Leeds Teaching Hospital NHS Trust, Leeds, UK. ^114^North East Thames Regional Genetics Service, Great Ormond Street Hospital for Children NHS Foundation Trust, London, UK. ^115^Northern Genetics Service, Newcastle upon Tyne Hospitals NHS Foundation Trust, Newcastle upon Tyne, UK. ^116^Newcastle University, Newcastle upon Tyne, UK. ^117^Newcastle upon Tyne Hospitals NHS Foundation Trust, Newcastle upon Tyne, UK. ^118^National Heart and Lung Institute, Imperial College London, London, UK. ^119^Royal Brompton Hospital, Royal Brompton and Harefield NHS Foundation Trust, London, UK. ^120^Clinical Genetics Department, Guy’s and St Thomas NHS Foundation Trust, London, UK. ^121^King’s College Hospital NHS Foundation Trust, London, UK. ^122^Guy’s and St Thomas’ Hospital, Guy’s and St Thomas’ NHS Foundation Trust, London, UK. ^123^MRC London Institute of Medical Sciences, Imperial College London, London, UK. ^124^National Heart Research Institute Singapore, National Heart Centre Singapore, Singapore, Singapore. ^125^Division of Cardiovascular & Metabolic Disorders, Duke-National University of Singapore, Singapore, Singapore. ^126^Department of Biotechnology, Graduate School of Engineering, Osaka University, Suita, Osaka, Japan. ^127^Women’s Health Research Centre, Surgery and Cancer, Faculty of Medicine, Hammersmith Hospital, Imperial College Healthcare NHS Trust, London, UK. ^128^Ramón Sardá Mother’s and Children’s Hospital, Buenos Aires, Argentina. ^129^Robinson Research Institute, Discipline of Obstetrics and Gynaecology, The University of Adelaide, Women’s and Children’s Hospital, Adelaide, Australia. ^130^Department of Molecular and Clinical Medicine, Sahlgrenska Academy, University of Gothenburg, Gothenburg, Sweden. ^131^Ophthalmology Department, UCSF School of Medicine, San Francisco, USA. ^132^Wellcome Centre for Mitochondrial Research, Institute of Genetic Medicine, Newcastle University, Newcastle upon Tyne, UK. ^133^Institute of Genetic Medicine, Newcastle University, Newcastle upon Tyne, UK. ^134^Institute of Genetic Medicine, Newcastle University, Newcastle upon Tyne, UK. ^135^John Walton Muscular Dystrophy Research Centre, Institute of Genetic Medicine, Newcastle University, Newcastle upon Tyne, UK. ^136^NIHR Biomedical Research Centre at Moorfields Eye Hospital and UCL Institute of Ophthalmology, London, UK. ^137^Yorkshire Regional Genetics Service, Chapel Allerton Hospital, Leeds Teaching Hospitals NHS Trust, Leeds, UK. ^138^Department of Clinical Genetics, Royal Devon & Exeter Hospital, Royal Devon and Exeter NHS Foundation Trust, Exeter, UK. ^139^West Midlands Regional Genetics Service, Birmingham Women’s and Children’s NHS Foundation Trust, Birmingham, UK. ^140^Manchester University NHS Foundation Trust, Manchester, UK. ^141^Department of Clinical Genetics, Liverpool Women’s NHS Foundation, Liverpool, UK. ^142^Department of Clinical Genetics, St George’s University Hospitals NHS Foundation Trust, London, UK. ^143^Department of Clinical Genetics, Guy’s and St Thomas’ NHS Foundation Trust, London, UK. ^144^Manchester Centre for Genomic Medicine, St Mary’s Hospital, Manchester Universities Foundation NHS Trust, Manchester, UK. ^145^Department of Clinical Genetics, Nottingham University Hospitals NHS Trust, Nottingham, UK. ^146^Wessex Clinical Genetics Service, University Hospital Southampton NHS Foundation Trust, Southampton, UK. ^147^Developmental Neurosciences, UCL Great Ormond Street Institute of Child Health, London, UK. ^148^Department of Neurology, Great Ormond Street Hospital for Children NHS Foundation Trust, London, UK. ^149^Salford Royal NHS Foundation Trust, Salford, UK. ^150^Faculty of Biology, Medicine and Health, School of Biological Sciences, Division of Neuroscience and Experimental Psychology, University of Manchester, Manchester, UK. ^151^Department of Clinical Neurophysiology, Manchester University NHS Foundation Trust, Manchester, Manchester Academic Health Science Centre,, Manchester, UK. ^152^National Institute for Health Research/Wellcome Trust Clinical Research Facility, Manchester, UK. ^153^The National Hospital for Neurology and Neurosurgery, University College London Hospitals NHS Foundation Trust, London, UK. ^154^MRC Centre for Neuromuscular Diseases, Department of Molecular Neuroscience, UCL Institute of Neurology, London, UK. ^155^Pain Research, Department of Surgery and Cancer, Faculty of Medicine, Imperial College London, London, UK. ^156^Pain Medicine, Chelsea and Westminster Hospital NHS Foundation Trust, London, UK. ^157^University Hospitals Birmingham NHS Foundation Trust, Birmingham, UK. ^158^Division of Clinical Biochemistry and Immunology, Cambridge University Hospitals NHS Foundation Trust, Cambridge, UK. ^159^Royal Papworth Hospital NHS Foundation Trust, Cambridge, UK. ^160^Department of Immunology, Leicester Royal Infirmary, Leicester, UK. ^161^Royal Free London NHS Foundation Trust, London, UK. ^162^Nottingham University Hospitals NHS Trust, Nottingham, UK. ^163^Regional Immunology Service, The Royal Hospitals, Belfast, UK. ^164^Queen’s University Belfast, Belfast, UK. ^165^Sheffield Teaching Hospitals NHS Foundation Trust, Sheffield, UK. ^166^University Hospitals of North Midlands NHS Trust, Stoke-on-Trent, UK. ^167^East Yorkshire Regional Adult Immunology and Allergy Unit, Hull Royal Infirmary, Hull and East Yorkshire Hospitals NHS Trust, Hull, UK. ^168^Barts Health NHS Foundation Trust, London, UK. ^169^Birmingham Heartlands Hospital, University Hospitals Birmingham NHS Foundation Trust, Birmingham, UK. ^170^Royal Hospital for Children, NHS Greater Glasgow and Clyde, Glasgow, UK. ^171^Epsom & St Helier University Hospitals NHS Trust, London, UK. ^172^Immunodeficiency Centre for Wales, University Hospital of Wales, Cardiff, IUK. ^173^Centre for Immunology & Vaccinology, Chelsea & Westminster Hospital, Department of Medicine, Imperial College London, London, UK. ^174^Department of Respiratory Medicine Royal Brompton & Harefield NHS Foundation Trust, London, UK. ^175^Sandwell and West Birmingham Hospitals NHS Trust, Birmingham, UK. ^176^Scunthorpe General Hospital, Northern Lincolnshire and Goole NHS Foundation Trust, Scunthorpe, UK. ^177^Gartnavel General Hospital, NHS Greater Glasgow and Clyde, Glasgow, UK. ^178^Queen Elizabeth University Hospital, Glasgow, UK. ^179^Birmingham Chest Clinic and Heartlands Hospital, University Hospitals Birmingham NHS Foundation Trust, Birmingham, UK. ^180^Frimley Park Hospital, NHS Frimley Health Foundation Trust, Camberley, UK. ^181^Institute of Cellular Medicine, Faculty of Medical Sciences, Newcastle University, Newcastle upon Tyne, UK. ^182^Imperial College Renal and Transplant Centre, Hammersmith Hospital, Imperial College Healthcare NHS Trust, London, UK. ^183^Children’s Renal and Urology Unit, Nottingham Children’s Hospital, QMC, Nottingham University Hospitals NHS Trust, Nottingham, UK. ^184^The National Renal Complement Therapeutics Centre, Royal Victoria Infirmary, Newcastle upon Tyne, UK. ^185^MPGN/C3 Glomerulopathy Rare Renal Disease group, UK. ^186^Department of Paediatric Nephrology, Great North Children’s Hospital, Newcastle upon Tyne Hospitals NHS Foundation Trust, Newcastle upon Tyne, UK. ^187^Department of Pulmonary Medicine, VU University Medical Centre, Amsterdam, The Netherlands. ^188^Golden Jubilee National Hospital, Glasgow, UK. ^189^Sheffield Pulmonary Vascular Disease Unit, Royal Hallamshire Hospital NHS Foundation Trust, Sheffield, UK. ^190^National Pulmonary Hypertension Service (Newcastle), The Newcastle upon Tyne Hospitals NHS Foundation Trust, Newcastle upon Tyne, UK. ^191^Department of Molecular Medicine, General Biology, and Medical Genetics Unit, University of Pavia, Pavia, Italy. ^192^Departement de Genetique & ICAN, Hopital Pitie-Salpetriere, Assistance Publique Hopitaux de Paris, Paris, France. ^193^University of Giessen and Marburg Lung Center (UGMLC), Giessen, Germany. ^194^Division of Cardiology, Fondazione IRCCS Policlinico S. Matteo, Pavia, Italy. ^195^Univ. Paris-Sud, Faculty of Medicine, University Paris-Saclay, Le Kremlin Bicetre, France. ^196^Service de Pneumologie, Centre de Reference de l’Hypertension Pulmonaire, Hopital Bicetre (Assistance Publique Hopitaux de Paris), Le Kremlin Bicetre, France. ^197^INSERM U999, Hospital Marie Lannelongue, Le Plessis Robinson, France. ^198^National Pulmonary Hypertension Service, Imperial College Healthcare NHS Trust, London, UK. ^199^Ludwig Boltzmann Institute for Lung Vascular Research, Graz, Austria. ^200^Dept of Internal Medicine, Division of Pulmonology, Medical University of Graz, Graz, Austria. ^201^Department of Infection, Immunity & Cardiovascular Disease, University of Sheffield, Sheffield, UK. ^202^Royal United Hospitals Bath NHS Foundation Trust, Bath, UK. ^203^Department of Clinical Genetics, VU University Medical Centre, Amsterdam, The Netherlands. ^204^Imperial College London, London, UK. ^205^North Bristol NHS Trust, Bristol, UK. ^206^Division of Nephrology and Center for Precision Medicine and Genomics, Department of Medicine Columbia University Vagelos College of Physicians and Surgeons, New York, USA. ^207^Institute of Genomic Medicine and the Department of Genetics and Development, Columbia University Vagelos College of Physicians and Surgeons, New York, NY 10032, New York, USA. ^208^Evolution and Genomic Sciences, Faculty of Biology, Medicine and Health, University of Manchester, Manchester, UK. ^209^Genetics and Genomic Medicine Programme, UCL Great Ormond Street Institute of Child Health, London, UK. ^210^London Centre for Paediatric Endocrinology and Diabetes, Great Ormond Street Hospital for Children, London, UK. ^211^NIHR Great Ormond Street Biomedical Research Centre, London, UK. ^212^Arthritis Research UK Centre for Adolescent Rheumatology at UCL UCLH and GOSH, London, UK. ^213^Florence Nightingale Faculty of Nursing, Midwifery & Palliative Care, King’s College London, London, UK. ^214^St Johns Institute of Dermatology, St Thomas’ Hospital, Guy’s and St Thomas’ NHS Foundation Trust, London, UK. ^215^Oxford Medical Genetics Laboratories, Oxford University Hospitals NHS Foundation Trust, Oxford, UK. ^216^MRC Weatherall Institute of Molecular Medicine, University of Oxford, John Radcliffe Hospital, Oxford, UK. ^217^Department of Clinical Genetics, Churchill Hospital, Oxford University Hospitals NHS Trust, Oxford, UK. ^218^Institute of Cancer and Genomic Sciences, Institute of Biomedical Research, University of Birmingham, Birmingham, UK. ^219^Department of Clinical Immunology, John Radcliffe Hospital, Oxford University Hospitals NHS Foundation Trust, Oxford, UK. ^220^Department of Haematology, Oxford University Hospital Foundation Trust, Oxford, UK. ^221^Oxford Epilepsy Research Group, Nuffield Department of Clinical Neurosciences, University of Oxford, Oxford, UK. ^222^Nuffield Department of Surgery, University of Oxford, Oxford, UK. ^223^Newcastle Eye Centre, Royal Victoria Infirmary, The Newcastle upon Tyne Hospitals NHS Foundation Trust, Newcastle upon Tyne, UK. ^224^NIHR Centre for Aging, Newcastle University, Newcastle upon Tyne, UK. ^225^Newcastle BRC, Newcastle University, Newcastle upon Tyne, UK. ^226^UCL Institute of Cardiovascular Science, University College London, London, UK. ^227^Barts Heart Centre, St Bartholomew’s Hospital, Barts Health NHS Trust, London, UK.

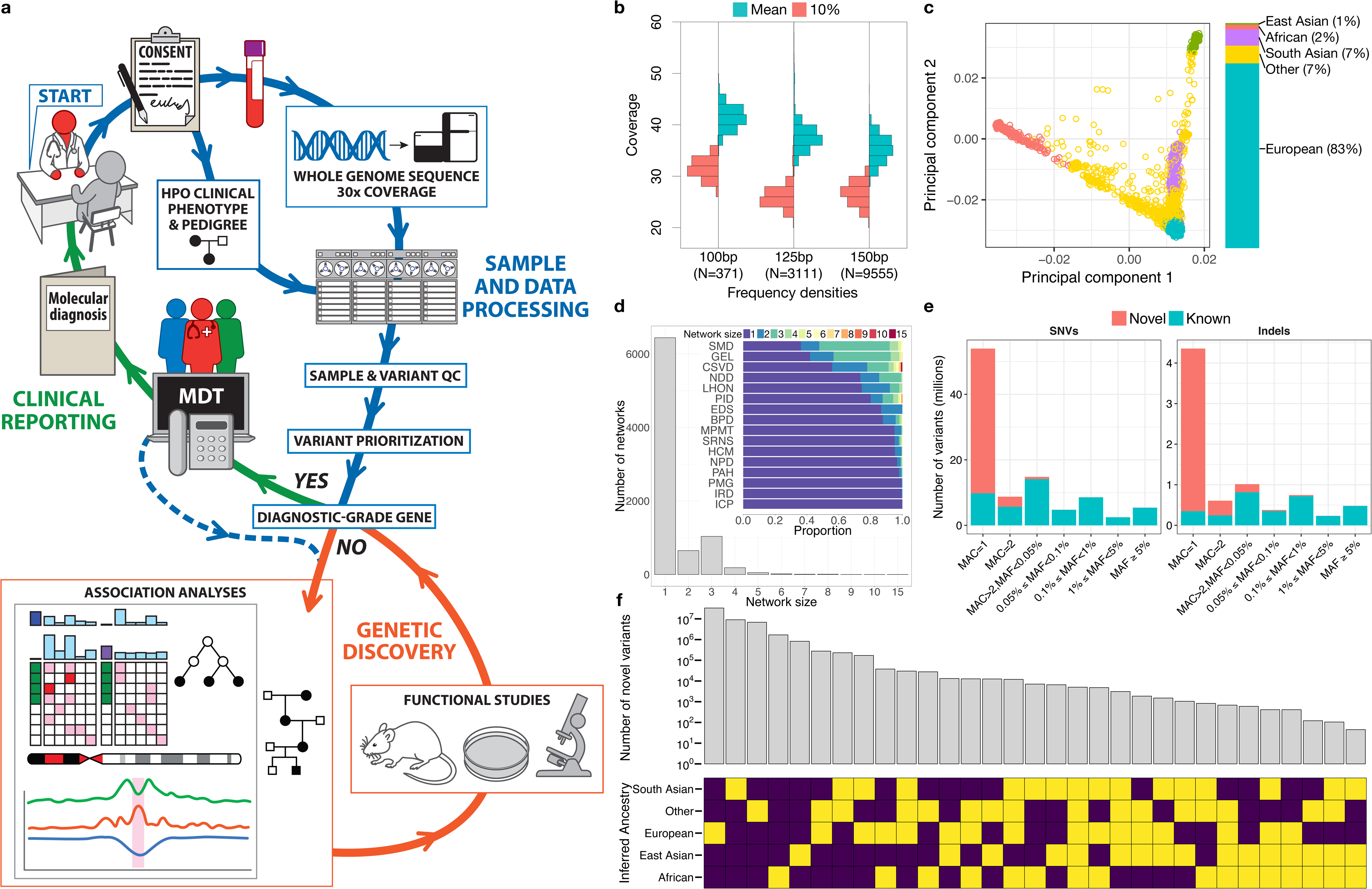

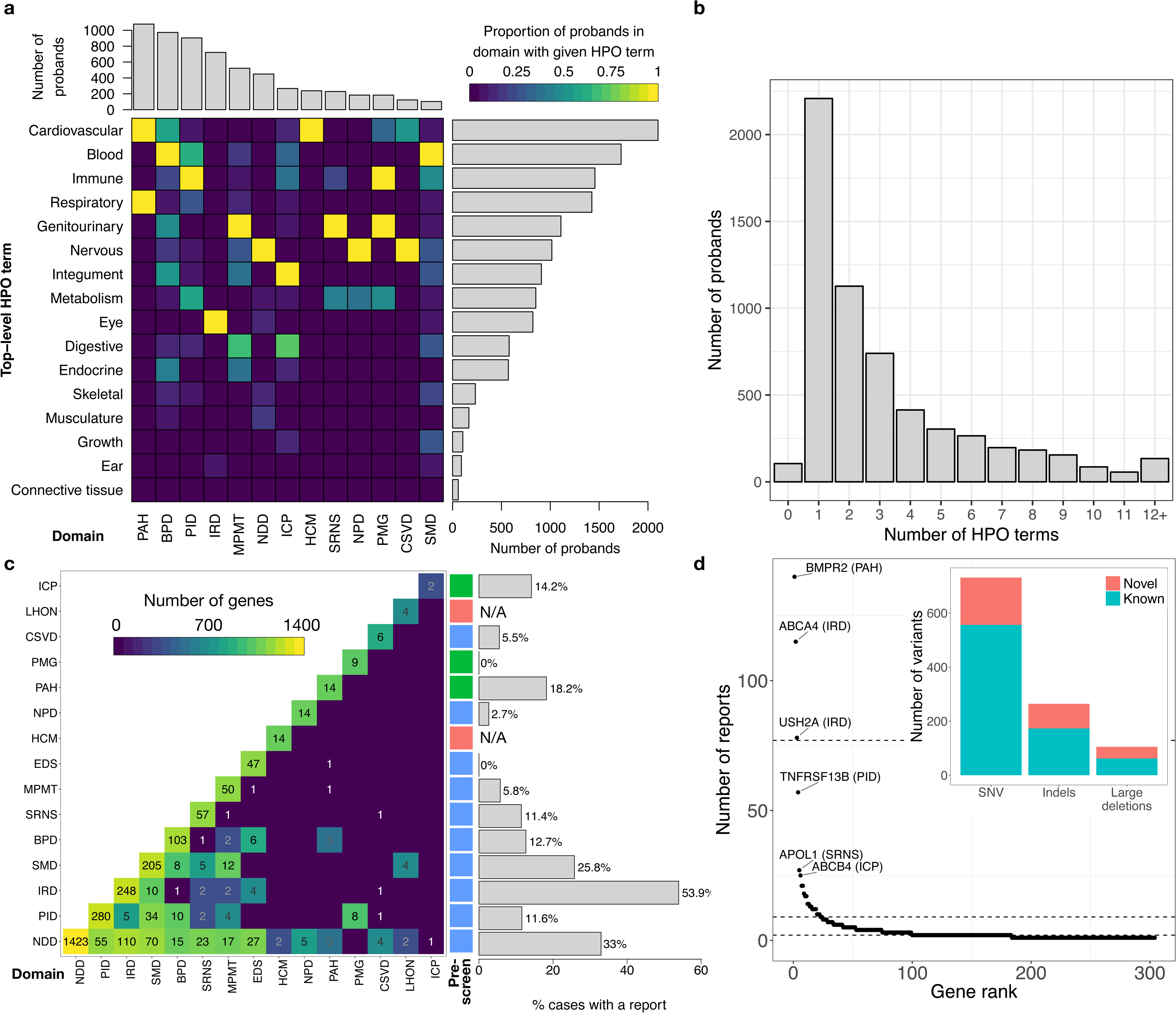

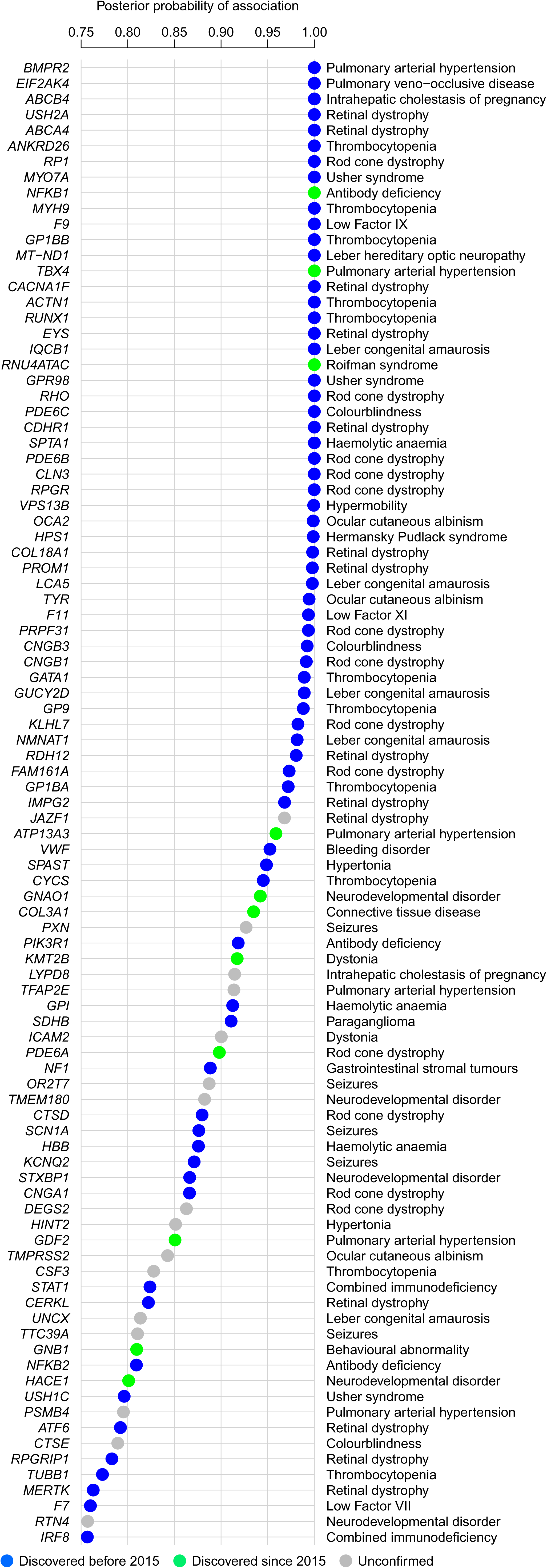

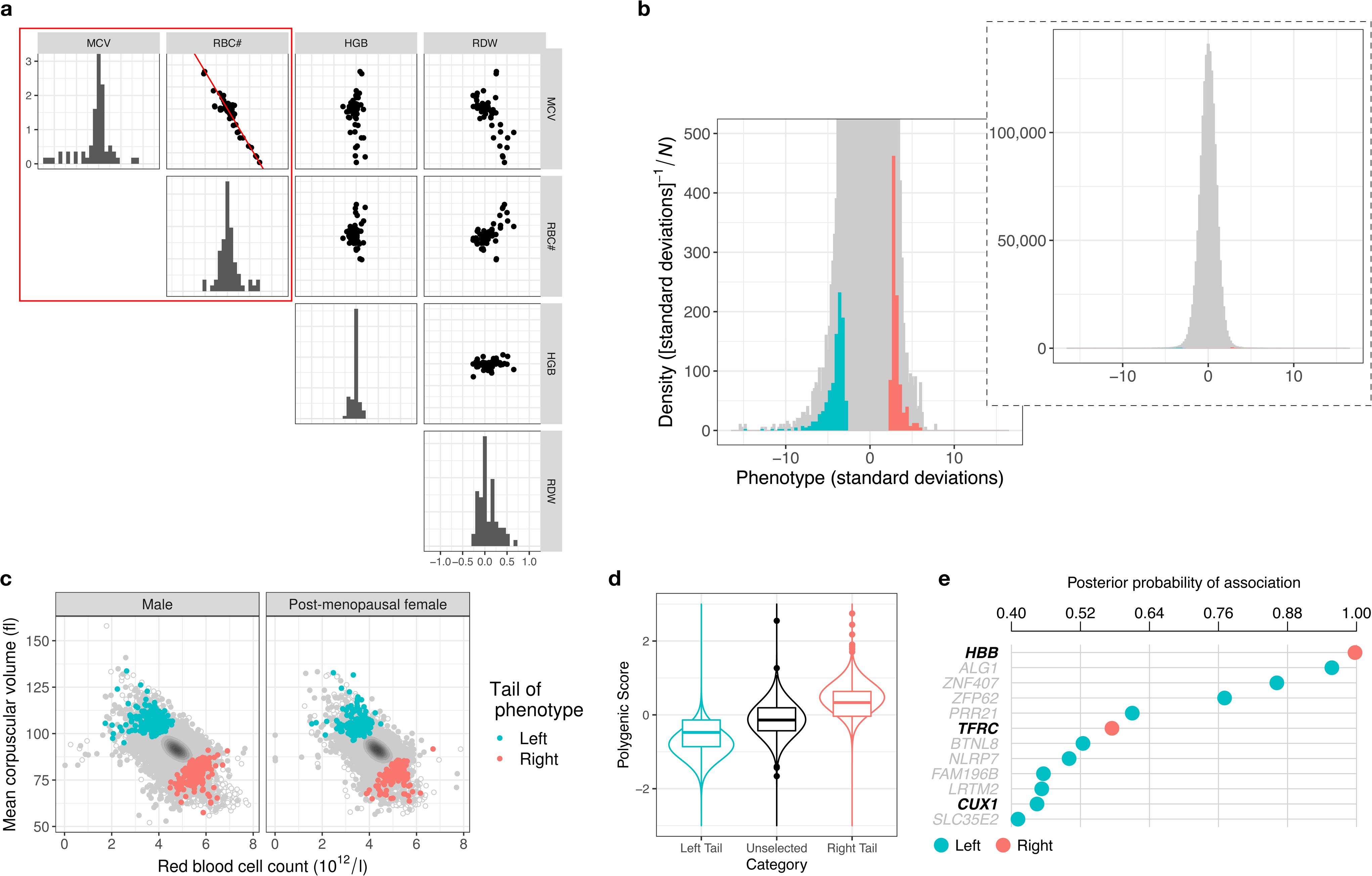

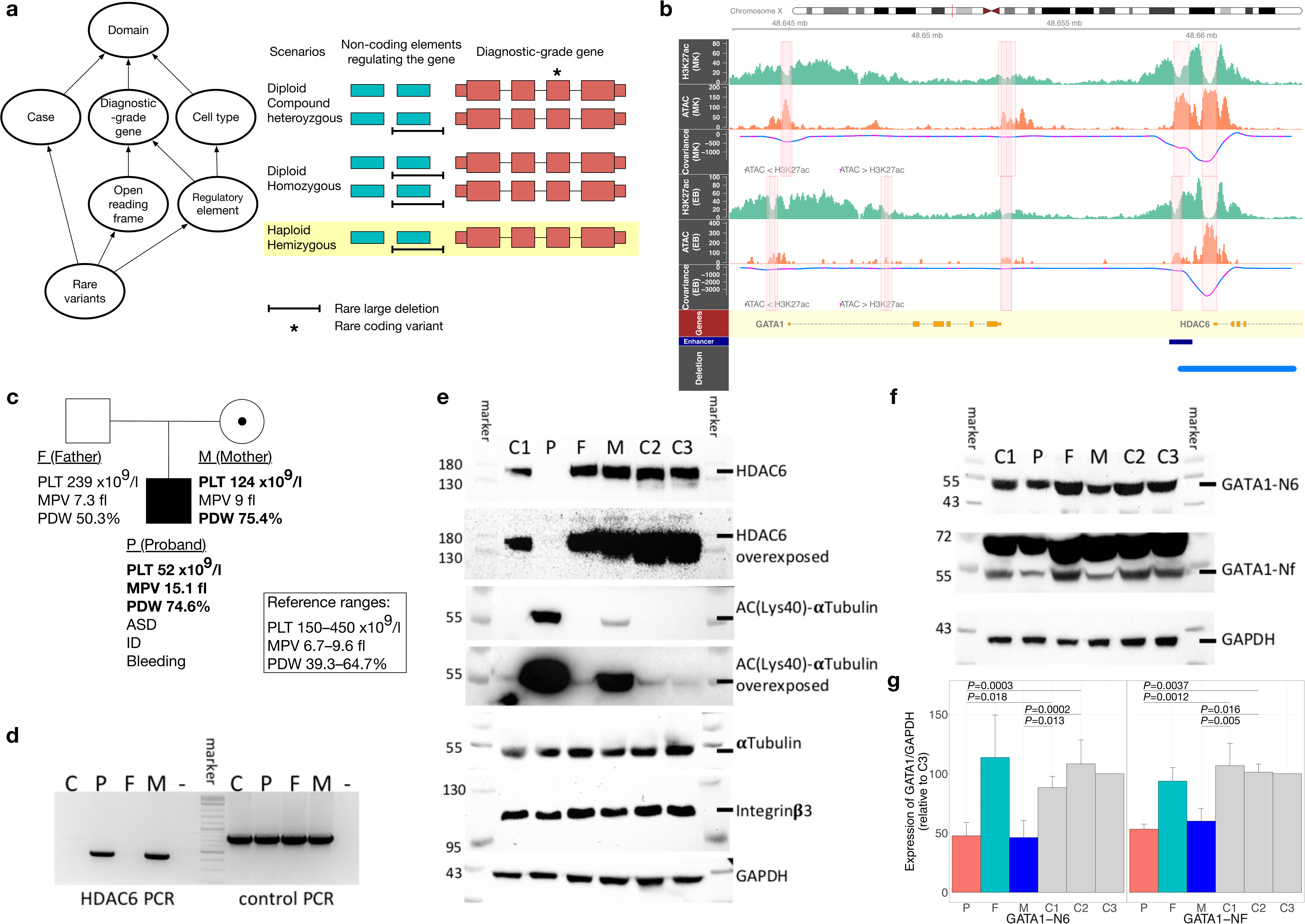

