## Supplementary material for "Whole-genome sequencing of rare disease patients in a national healthcare system": Legends for Main Figures

Main Text Figure Legends

**Fig. 1. Overview of the study and genetic data analysis. a**, Schematic depicting the flow of information through the study and the synergy between diagnosis and discovery. In blue: an undiagnosed patient is recruited into the study by his/her clinician, informed consent is obtained and the clinician enters HPO terms and pedigree information into the study database, biological samples are taken and DNA is sent to a single Illumina laboratory for WGS, sequencing data are transferred to a high performance computing cluster for bioinformatic QC and the prioritisation of variants in DGGs. In green: selected variants meeting predefined characteristics are presented to the MDTs using the Sapientia^TM^ web application, variants are categorised as pathogenic or likely pathogenic, a molecular diagnosis may be returned to the referring clinician. In orange: statistical and bioinformatic analyses are applied to the genetic and phenotypic data to identify aetiological variants, disease-mediating genes and regulatory regions. Participants and close relatives are invited to participate in co-segregation and functional studies, and model systems are used to study disease mechanisms. **b**, Histograms showing the distribution of read coverage across 13,037 samples, stratified by sequencing read lengths of 100bp,125bp and 150bp. **c**, Projection of participants onto the first two principal components of genetic variation in the 1000 Genomes Project (left sub-panel), bar plot showing the percentage of participants whose ancestry was assigned to different 1000 Genomes populations (right sub-panel). **d**, Bar plot showing the size distribution of genetically determined networks of closely related individuals across all 13,037 samples. Inset: Distributions of network sizes for each rare disease domain. **e**, Histograms illustrating the observed allele frequency distribution of variants measured in 10,259 unrelated samples, stratified by variant type (SNV or indel). Variants were labelled novel if they were uncatalogued in the following databases: 1000 Genomes, UK10K, TOPMed, gnomAD, HGMD Pro. MAC: minor allele count; MAF: minor allele frequency. **f**, Histogram counting (log_10_ scale) the novel variants according to the ancestry groups in which they were observed (yellow: present, navy: absent).

**Fig. 2. Phenotyping data, diagnostic-grade genes and MDT-reported results. a**, Bar plot showing the distribution of probands by domain (top); bar plot counting the number of probands with each top-level HPO term (right). The heat map shows the proportion of probands in each domain who have been assigned a particular top-level HPO term. Top-level HPO terms have been abbreviated. The full term names read ‘Abnormality of,’ followed by, from top to bottom: the cardiovascular system; blood and blood-forming tissues; the respiratory system; the immune system; the genitourinary system; the nervous system; integument; metabolism/homeostasis; the eye; the endocrine system; the digestive system; the skeletal system; the musculature; of the ear; growth; connective tissue. **b**, Bar plot showing the count distribution of the number of HPO terms assigned to affected probands in 13 rare disease domains. **c**, Heat map showing the number of DGGs shared by pairs of domains (left). Pre-screening level for each domain indicated in red (full), blue (partial) or green (none). Bar plot of the proportion of cases in each domain for which a clinical report was issued (right). **d**, Number of reports issued per DGG ordered inversely by count. Dashed lines indicate quartiles of the count distribution. Inset: bar plot showing the number of distinct clinically reported variants stratified by variant type (SNVs, indels and large deletions). The proportion of each bar coloured iris blue/salmon indicates the proportion of variants which are known/novel (absent from HGMD Pro and from the set of variants in ClinVar having a likely pathogenic or pathogenic interpretation and without any benign interpretations) .

**Fig. 3. BeviMed genetic association results for rare diseases.** BeviMed was applied gene-wise to infer associations between the genotypes of filtered rare variants and various case/control groupings (tags). For a given gene, only the maximum PP over tags was retained, to account for correlation between tags. The PPs for genetic association inferred by BeviMed exceeding 0.75 are shown. Gene names are given on the left and the corresponding tag names of the case groupings are given on the right. The green and blue colouring denotes genetic associations supported by original scientific publications since or before 2015, respectively, while grey denote associations that are currently unconfirmed in the literature. The mean of the PPs above the threshold is equal to 0.93, implying a posterior expectation that 93% of the reported associations are true.

**Fig 4. Polygenic and rare variant associations with extreme RBC traits in UK Biobank. a**, Graphical summaries of the joint distribution of the estimated per allele additive effect of 65 variants with MAF < 1% on the mean of four rank inverse normalised quantitative properties of RBCs: mean corpuscular volume (MCV), RBC count (RBC#), haemoglobin concentration (HGB) and RBC distribution width (RDW). The 65 variants were chosen for being significantly associated with at least one of the 12 red cell traits in Astle *et al*. The on-diagonal panels depict the univariate distribution of the estimated effect sizes of each trait (measured in standard deviations per allele) and the off-diagonal panels depict the bivariate relationship between the estimated effect sizes. The red square highlights the bivariate marginal distribution used to develop the quantitative selection phenotype. The red line in the (RBC#, MCV) panel was estimated by a Deming regression of the MCV effect sizes on the RBC# effect sizes. **b**, Both sub-panels show the distribution of the (centred and standardised) quantitative phenotype in the European ancestry male and post-menopausal female UK Biobank participants without a baseline self report or medical history of an illness or treatment known to perturb RBC indices (grey density histograms) and the distribution of the phenotype in individuals selected from the left (iris blue density histogram) and right (salmon density histogram) tails. The density scale has been chosen so that the area under each histogram represents the respective number of contributing individuals, *N*=316,739. The lower left sub-panel is a vertical stretch of the bottom part of the overlying upper right sub-panel. Many participants in the extreme tails were not selected because of poor quality DNA in the UK Biobank archive or because of recalibration of the phenotype to ensure a representative distribution of age and sex amongst those selected. **c**, Bivariate scatter showing the distribution of RBC# and MCV (both adjusted for technical but not biological noise) in UK Biobank males (left sub-panel) and post-menopausal females (right sub-panel). The overlaid ellipsoids are contours from kernel density estimates of the central parts of the distributions. The closed grey circles represent participants contributing to the grey density histograms in sub-figure 4b. The (underlying) open circles represent participants excluded from selection for sequencing on the grounds of ancestry or medical history. The excess of open circles in the bottom right of each sub-panel is probably explained by the high prevalence of thalassemia in participants with African or Mediterranean ancestry. The coloured circles indicate the participants selected for sequencing from the two tails of the phenotype. **d**, Each box plot summarises the distribution of a polygenic score developed to predict the quantitative phenotype using variants associated with RBC# or MCV by Astle *et al*. in the indicated category of study participants. The ‘Unselected’ category is composed of unrelated European participants whose disease phenotype is explained fully by rare variants. The underlying violin plots show the expected distribution of the polygenic score under a Gaussian variance components model, conditional on the proportion of phenotypic variance explained by the score and the tail selection thresholds. **e**, BeviMed computed PPs for genetic association with each of the tails of the phenotype (distinguished by colour), for genes with PPs > 0.4. The strength of concordant biological evidence for the genes in bold is such that they can be considered positive controls.

**Fig. 5. Causal variants in regulatory elements. a**, Left: schematic of the procedure for identifying causal deletions in REs of DGGs. Right: schematic showing models of expanded DGGs (in salmon) including regulatory elements (in iris blue) and three possible genetic architectures underlying a rare disease. **b**, From top to bottom: X chromosome, with the position on the chromosome shown with a red bar, genomic coverage of H3K27ac ChIP-seq (green) in MKs, genomic coverage of ATAC-seq (orange) in MKs, the smoothed covariance between MK H3K27ac ChIP-seq and MK ATAC-seq coverage, which was used to call the REs indicated by the shaded pink panels in overlay, the corresponding three epigenetic tracks and overlays for erythroblasts, gene exons in orange, the *GATA1* enhancer as a dark blue horizontal bar and the large deletion in the proband as a light blue horizontal bar. An REs overlapping the enhancer was identified by RedPop in MKs and EBs but not in the other four cell types (tracks for these other cell types not shown). The deleted element binds the transcription factors characteristic of the MK lineage: FLI1, GATA1/2, MEIS1, RUNX1 and TAL1 (not shown). **c**, Pedigree of proband (P) with thrombocytopenia and autism and his parents (F and M). PLT: platelet count, MPV: mean platelet volume, PDW: platelet distribution width, ASD: autism spectrum disorder, ID: Intellectual disability. **d**, Left: Gel electrophoresis showing presence and absence of short PCR amplicons using primers flanking the deletion. Right: control PCR (C); no DNA added indicated by ‘-‘. **e**, Representative immunoblots performed in duplicate for total platelet lysates for the indicated proteins and individuals. C1, C2 and C3 are controls. **f**, Representative immunoblots of total platelet lysates using two different GATA1 antibodies. **g**, Quantification of GATA1 protein levels obtained from three independent immunoblots (as per **f**) showing the mean and SEM and two-way ANOVA analysis *P*-values (multiple comparisons).
