## Supplementary figures and images for "Whole-genome sequencing of rare disease patients in a national healthcare system"

### Extended Data Figures

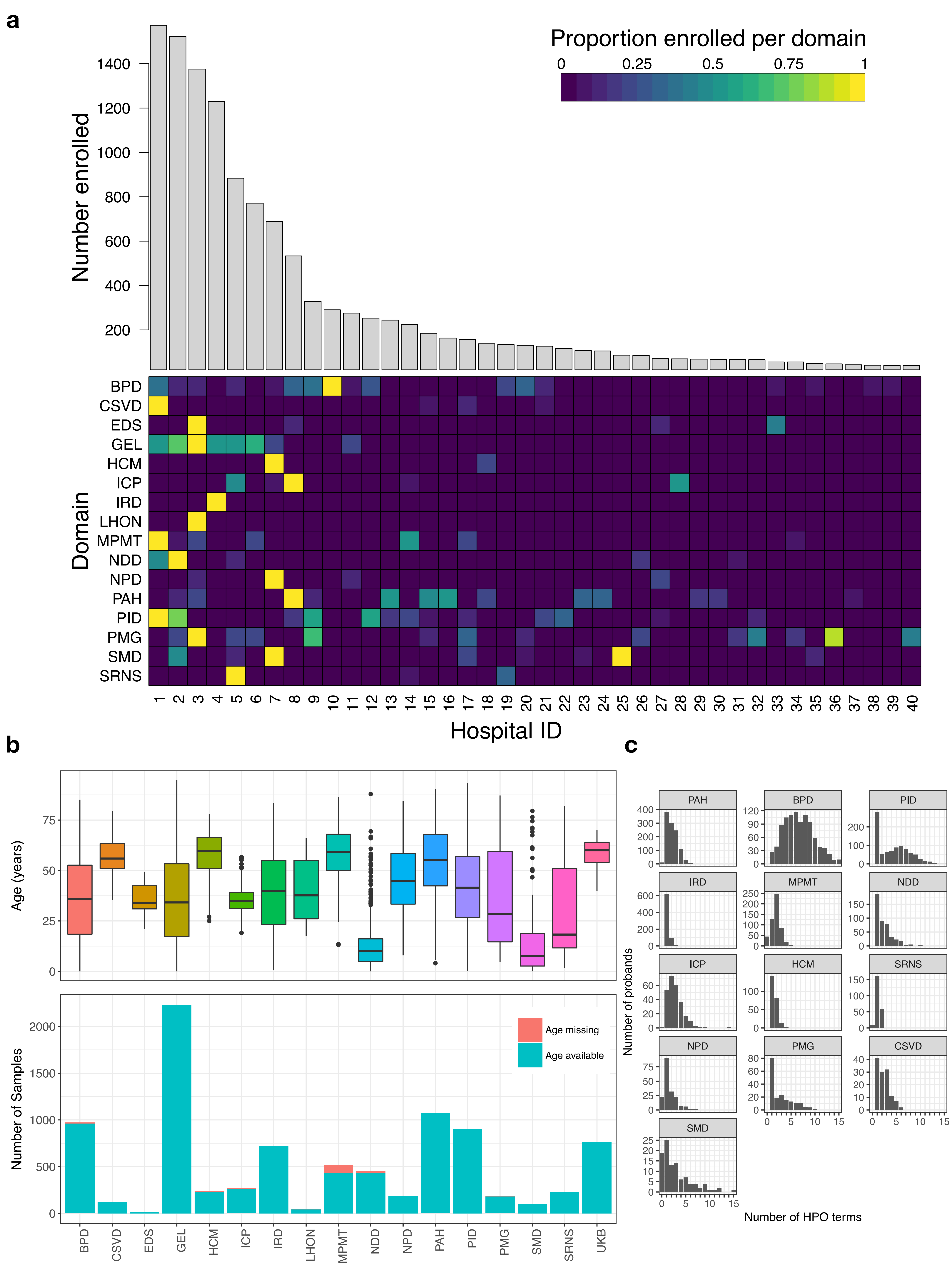

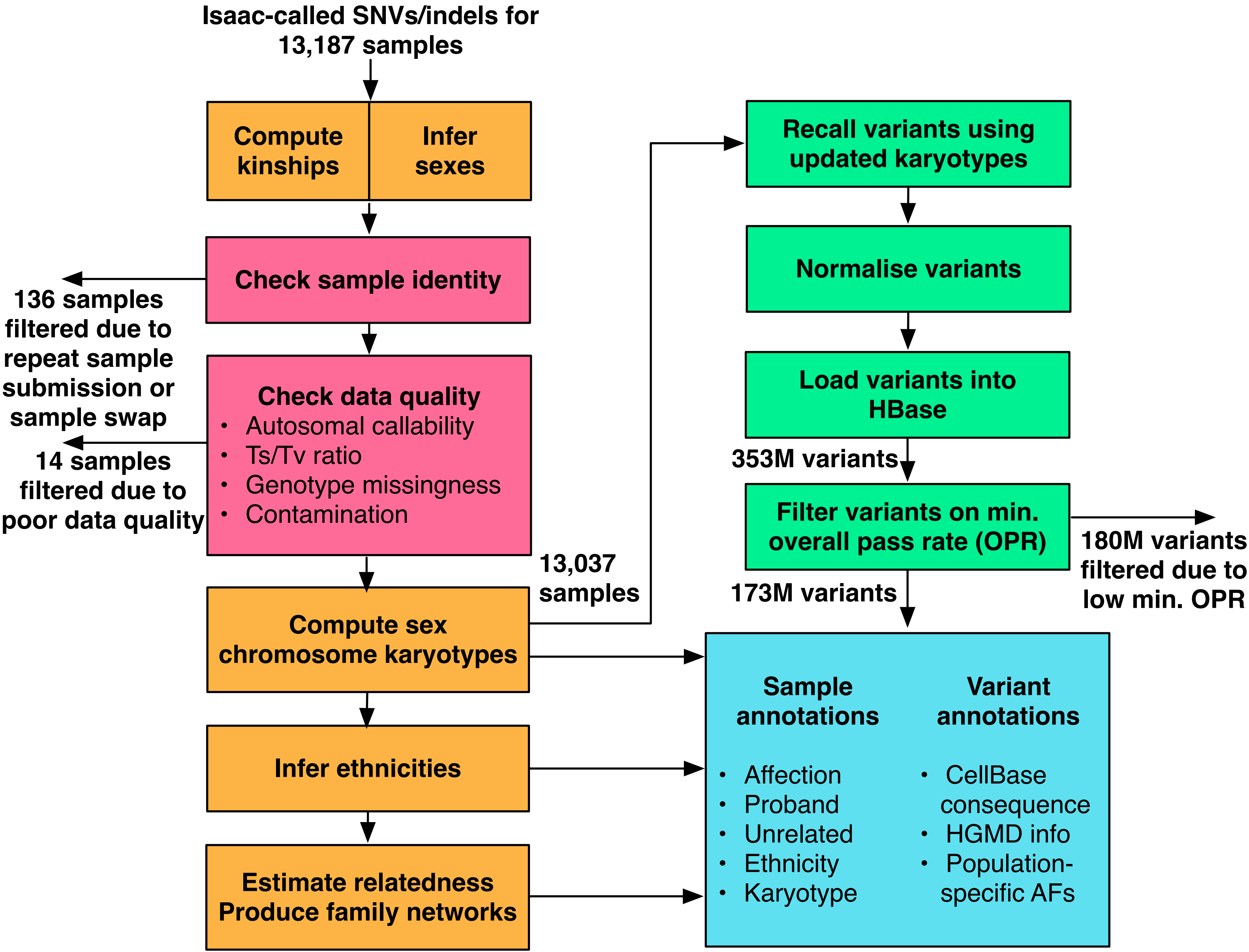

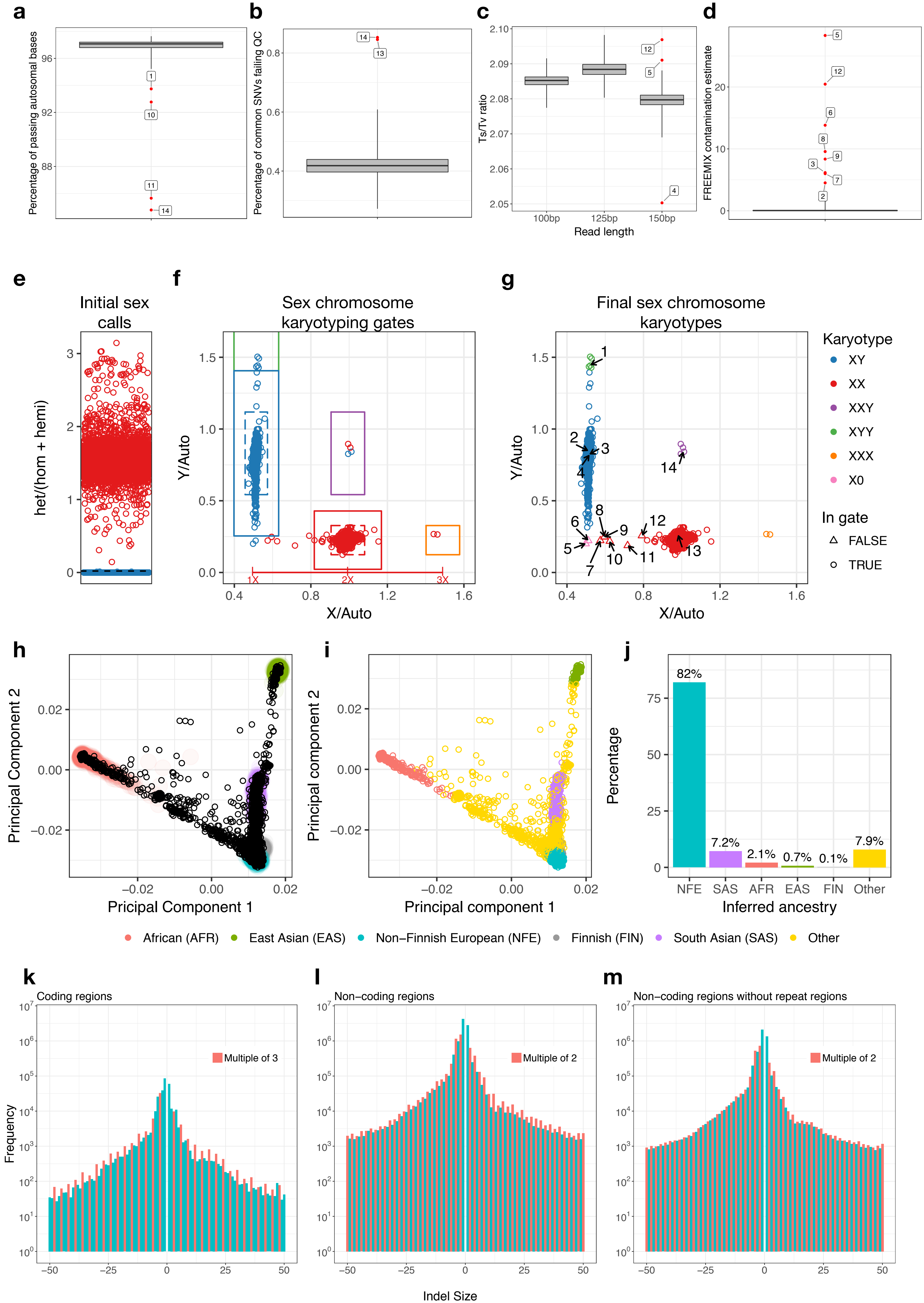

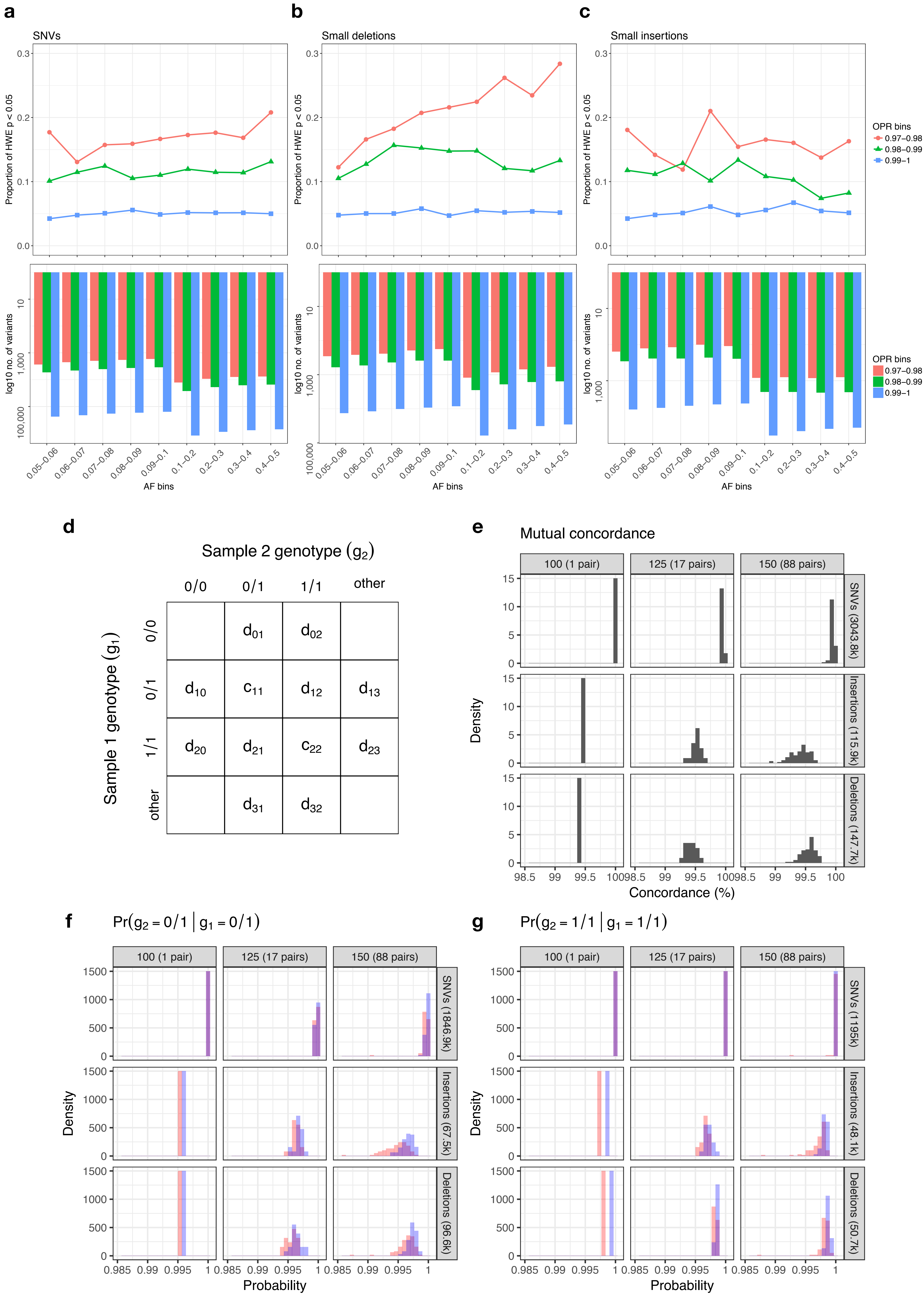

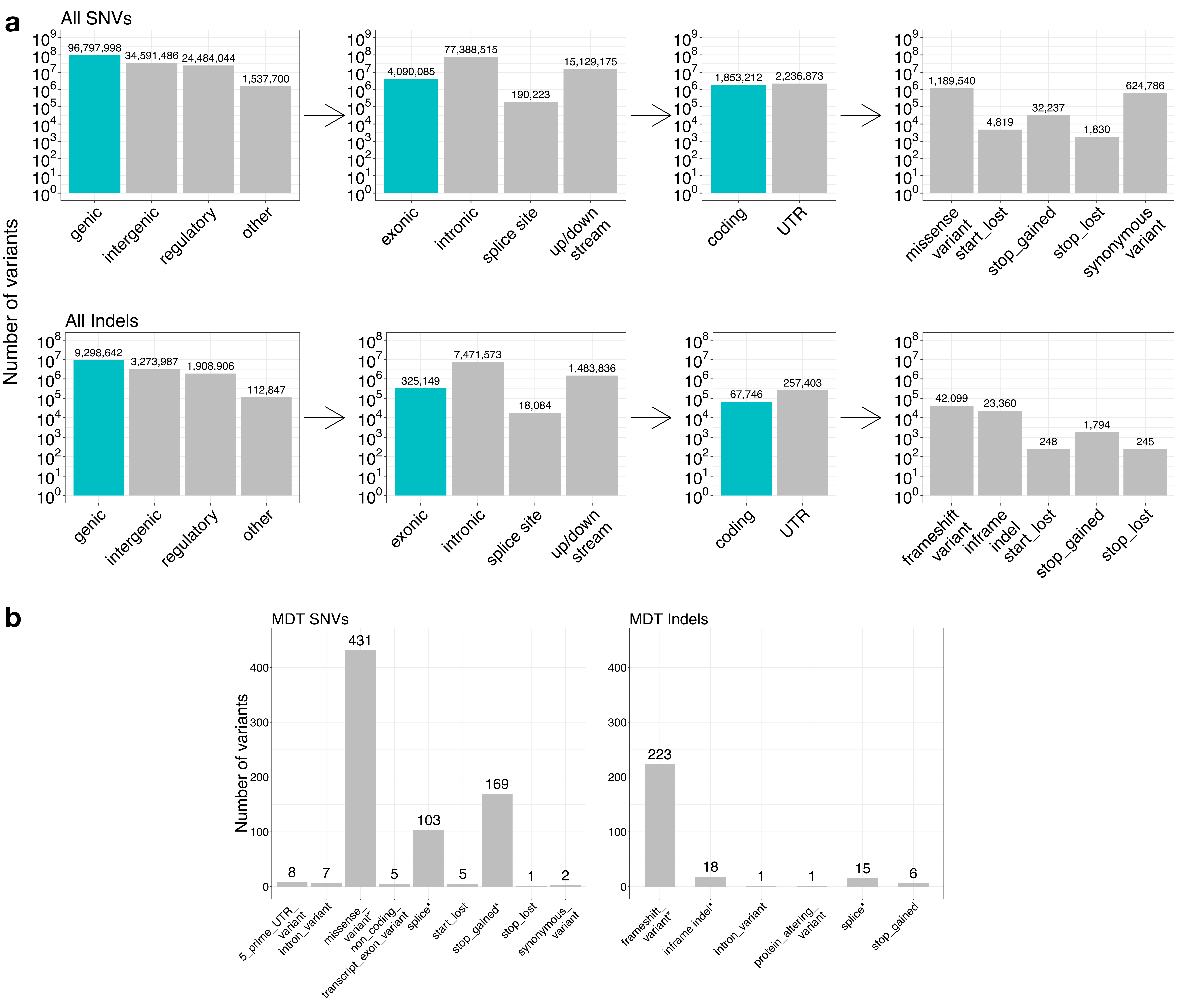

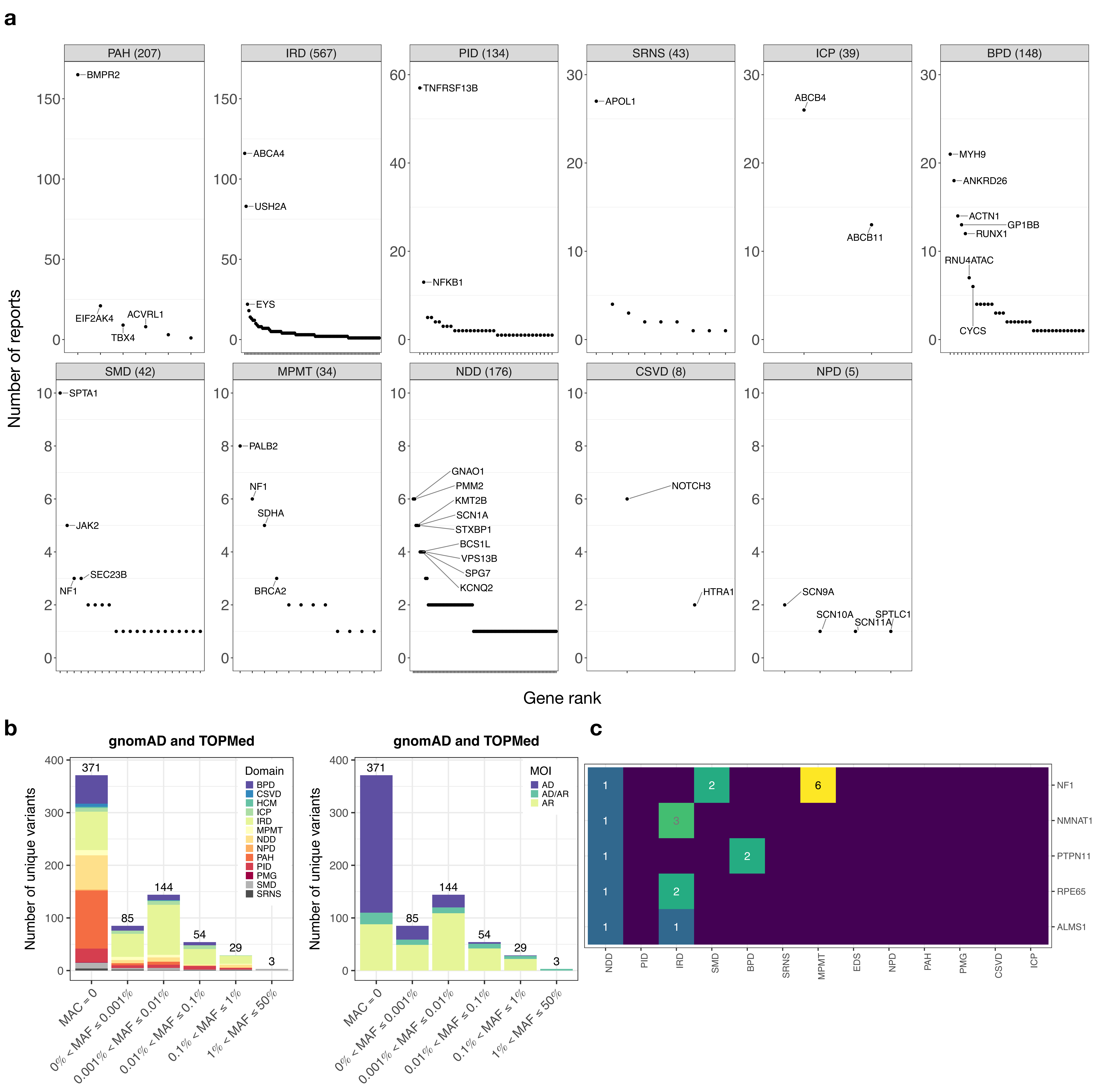

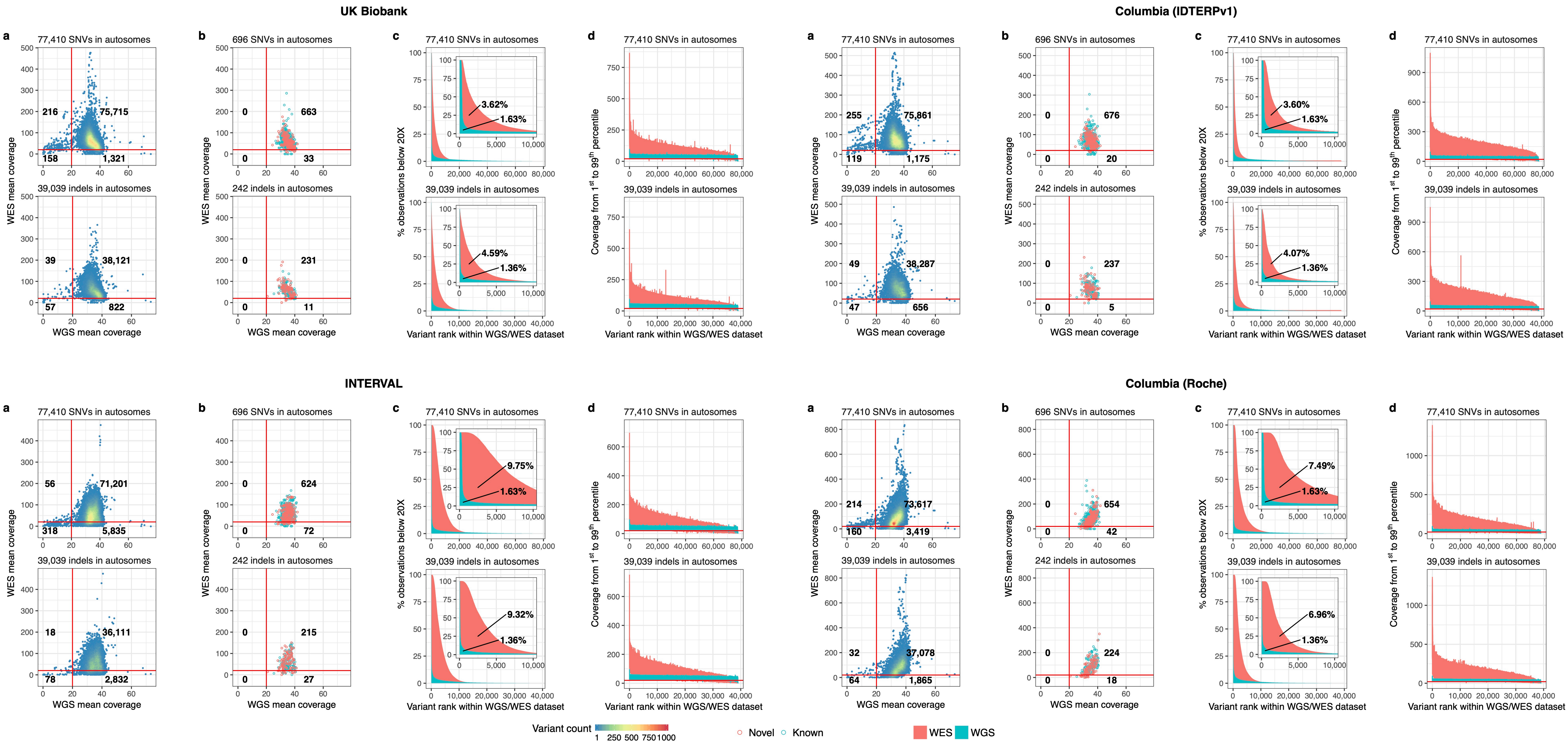

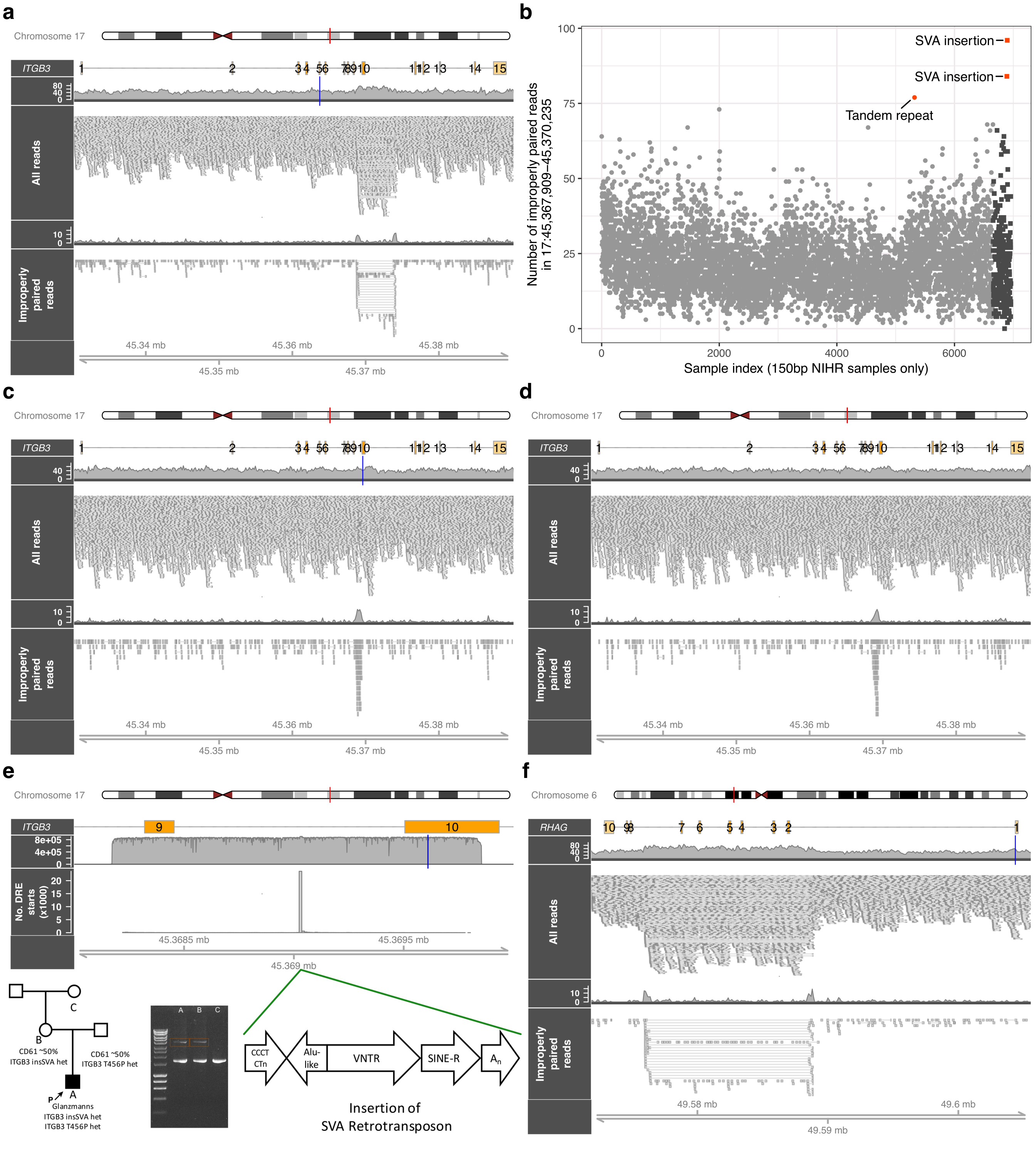

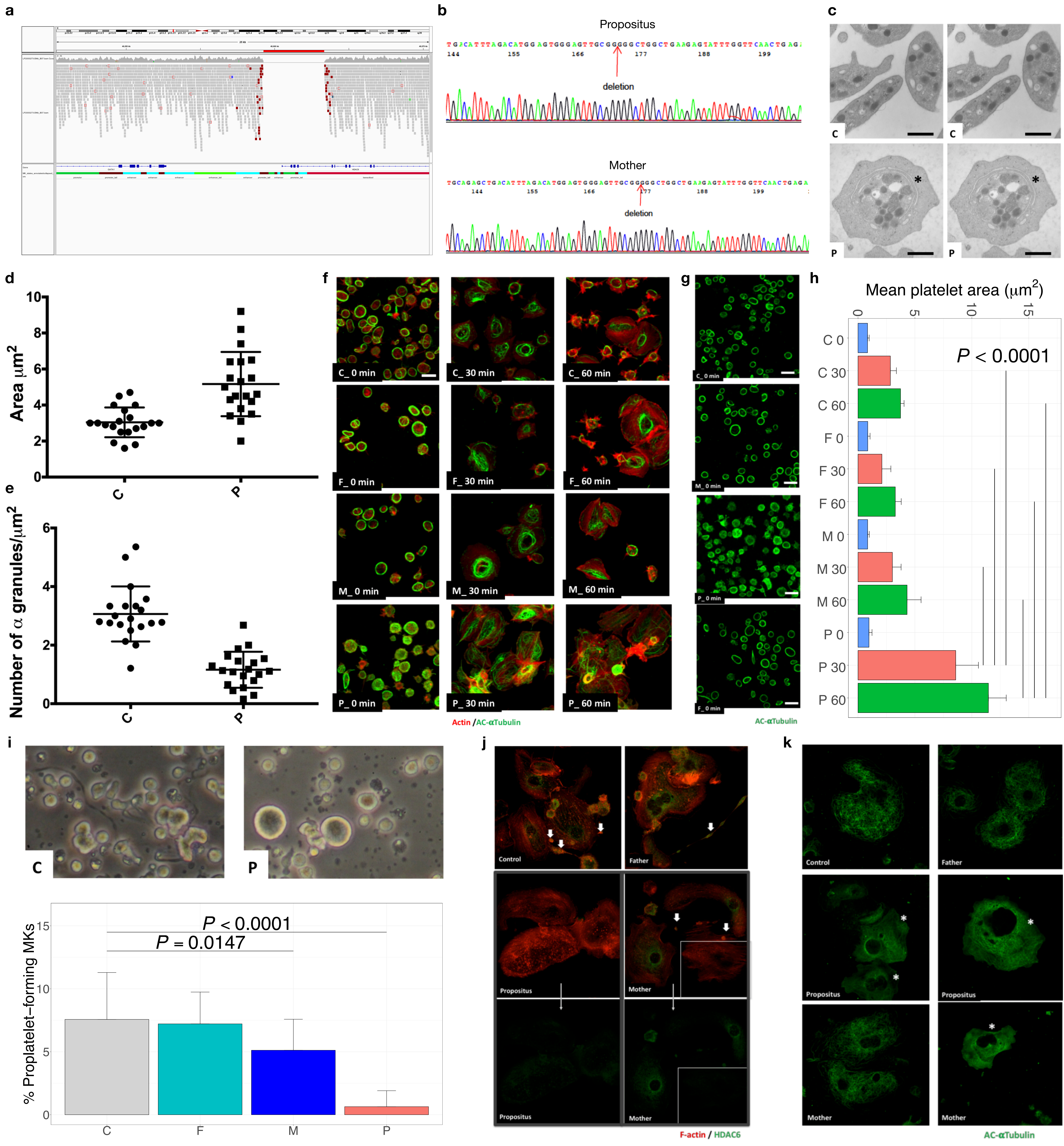

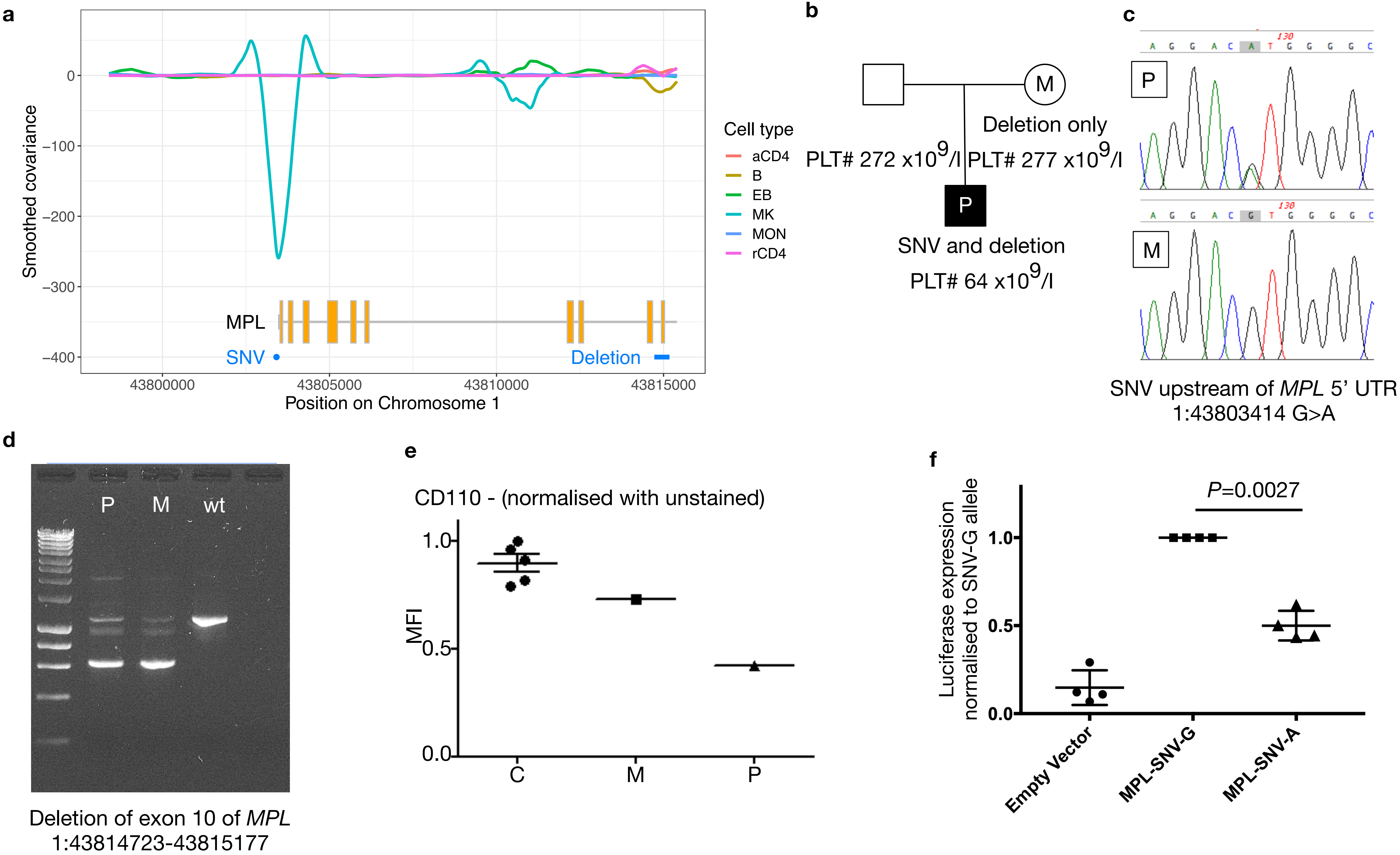
