## Supplementary material for "Whole-genome sequencing of rare disease patients in a national healthcare system": Legends for Extended Data Figures

Extended Data Figure Legends

**Extended Data Fig. 1: Demographic and phenotypic characteristics.**

**a**, Barplot of the number of enrolments for the 40 hospitals with at least 20 enrolled participants. The heat map shows the proportion of enrolments per domain in each of the 40 hospitals. Hospital IDs are detailed in Supplementary Table 1. **b**, Box plot of age at sampling for all probands for the 15 rare disease domains, GEL and UK Biobank above the corresponding stacked bar plots of number of probands for each domain with and without an available age at sampling. **c**, Histograms of the number of HPO terms appended to affected probands for 13 rare disease domains.

**Extended Data Fig. 2: Flow chart of the bioinformatic data processing.**

Flow chart describing the processing of samples and variants. Beginning at the top left, all samples were checked for data quality (see **Extended Data Fig. 3**) and sample identity upon receipt of data using regular quick kinship and sex checks. Samples failing sample QC, those samples with clearly incorrect sexes or lower quality versions of identical samples were removed before further analysis (pink boxes). Sex chromosome karyotypes, ethnicities, and relatedness/family trees were computed on these filtered samples (orange boxes) and variants were recalled for those samples with X/Y-chromosome ploidies different to that automatically predicted in the original pipeline. After variant normalisation, variant calls were loaded into HBase and merged, and summary statistics were calculated for both technical (100, 125, and 150bp) and population cohorts (e.g., unrelated African) (green boxes). A minimum OPR was calculated for each variant and filtering carried out on those below a cut-off of 0.99 (see **Extended Data Fig. 4**). Finally, variants were annotated in HBase with CellBase consequence information and annotation/AFs from external databases (e.g., gnomAD) (blue box).

**Extended Data Fig. 3: Sample QC, sex chromosome karyotyping and ancestry inference.**

**a**, Boxplot of the percentage of passing autosomal bases (4 exclusions). **b**, Boxplot of the percentage of common SNVs that failed quality control (2 exclusions). **c**, Batch-specific boxplots of Ts/Tv ratios (3 exclusions). **d**, Boxplot of FREEMIX values representing sample contamination (8 exclusions). Excluded samples are marked in red and labelled with an integer. Three samples were excluded due to failing more than one of the four QC checks (samples 5,12,14). **e**, H-ratios for 13,037 samples and predicted initial sexes. **f**, Scatterplot of ratios of X/Auto and Y/Auto coloured by the initial sex calls and showing the five sex karyotyping gates. **g**, Scatterplot of ratios of X/Auto and Y/Auto coloured by the final sex karyotype. Circles indicate samples falling within a sex karyotyping gate and triangles indicate samples falling outside all sex karyotyping gates. 1: confirmed XYY case; 2–4: confirmed XY female cases; 5, 6: confirmed XO cases; 7: confirmed XO case, this sample has some part of the second X chromosome present; 8–10: samples with large part of the X chromosome missing; 11–12: samples with multiple deletions on the X chromosome; 13: sample with two almost identical X chromosomes; 14: confirmed XXY case. **h**, Projection of the 13,037 samples, shown as round circles, onto the 1000 Genomes derived PCAs. The 1000 Genomes samples are shown as diffuse points underneath in colour. **i**, Projection of the 13,037 samples, shown as round circles, coloured by assigned population. **j**, Bar plot showing the number of individuals assigned to each population. The percentages are shown above each bar. NFE: Non-Finnish European; SAS: South Asian; AFR: African; EAS: East Asian; FIN: Finnish. **k**, **l**, **m**, Distribution of the sizes of small insertions (indel size > 0) and small deletions (indel size < 0) in coding regions, non-coding regions and non-coding regions excluding those in repetitive regions, specifically, the RepeatMasker track from the UCSC table browser and the Tandem Repeats Finder locations from the UCSC hg19 full data set download. In coding regions, natural selection against frameshift variants results in a systematic depletion of indel sizes that are not a multiple of 3bp. In non-coding regions, there is a slight excess of indel sizes that are a multiple of 2bp, but this pattern is almost indiscernible if repetitive regions are excluded.

**Extended Data Fig. 4: Variant QC.**

**a**, **b**, **c**, The proportion of HWE *P*-values < 0.05 amongst 8,510 unrelated Europeans across different AF bins is shown for SNVs, small deletions and small insertions. Boxplots of the number of variants in each OPR and AF bin are shown in the bottom panels. **d**, Combination of genotypes in a pair of samples, values in cells are the numbers of variants. **e, f, g**, Concordance of sequencing results for pairs of duplicates and twins with results from 100, 125 and 150bp reads from left to right. **e**, Distribution of mutual non-reference concordance in pairs of duplicates and twins. **f**, Probability of having a heterozygous variant in a sample, given its duplicate/twin has this heterozygous variant. **g**, Probability of having a homozygous variant in a sample, given its duplicate/twin has this homozygous variant. In panels **e**, **f** and **g**, the mean number of variants is shown in brackets after the variant type. In panels **f** and **g**, red and blue colours represent distribution of the lowest and highest of the two probabilities (sample 1 compared to sample 2 and sample 2 compared to sample 1) in a pair of duplicates/twins.

**Extended Data Fig. 5: Breakdown of genetic variants by their predicted primary consequence.**

a, Counts of SNVs and indels in various consequence classes shown on logarithmic scales with exact numbers above each bar. Variants in the green bars are subdivided into more granular regions of genome space in the following panel in a recursive manner from left to right. Categories have been chosen to represent the likely highest impact consequences at each stage: i.e., from left, overall genome space, within genes, exonic parts of genes, and protein coding regions. b, Count of MDT SNVs and indels in various consequence classes with exact numbers above each bar. A star denotes a super-category with *m*issense_variant including missense_variant or missense_variant&splice_region_variant; *s*plice including splice_acceptor_variant, splice_donor_variant, splice_donor_variant&coding_sequence_variant or splice_region_variant or splice_region_variant&intron_variant; *s*top_gained including stop_gained, stop_gained&splice_region_variant or stop_gained&splice; *f*rameshift variant including frameshift_variant, frameshift_variant&splice_region_variant or retained_intron; *i*nframe indel including inframe_deletion or inframe_insertion.

**Extended Data Fig. 6: Breakdown of diagnostic reports by domain.**

**a**, Number of reports issued for diagnostic-grade genes for 11 rare disease domains. Each panel shows a domain that issued reports, the title denotes the domain acronym and number of reports issued. Plots are presented in decreasing order by maximal y-axis scale. PMG and EDS are not included as no reports were issued for cases in these domains. Each point is a unique gene that was reported for that domain. The genes with the most reports issued per domain are labelled. Full details of all reports issued are in Supplementary Table 2. **b**, **c**, gnomAD and TOPMed frequency or allele count for the reported autosomal SNVs and indels in samples of European ancestry, per rare disease domains (**b**) and per mode of inheritance (**c**). MAC: Minor Allele Count. MAF: Minor Allele Frequency. Domain's acronyms as explained in Supp. Material. MOI: Mode Of Inheritance. AD: Autosomal Dominant. AR: Autosomal Recessive. For variants absent from gnomAD, both allele count (AC) and allele number (AN) were set to 0; for variants absent from TOPMed, AC was set to 0 and AN set to 125,568, in line with the AN value for all the present variants. For a given position and a given allele, the combined MAF was defined as the sum of ACs divided by the sum of ANs. The first bin in the plots (MAC=0) corresponds to variants not observed in either gnomAD or TOPMed. **c**, Number of report issued for diagnostic-grade genes shared between rare disease domains.

**Extended Data Fig. 7: Comparison of WGS and WES for genetic testing.**

For each of four WES datasets – UK Biobank, INTERVAL, Columbia (IDTERv1) and Columbia (Roche) – four sets of figures are shown for different sets of SNVs and indels, as follows. **a**, Scatterplot of WGS vs WES mean coverage. The red axes show the threshold for clinical reporting and the number of variants in each quadrant is indicated. **b**, Scatterplot of WGS vs WES coverage of the MDT-reported known (turquoise) and novel (salmon) SNVs and indels in autosomal diagnostic-grade genes. **c**, Superimposed scatterplots of the percentage of samples with WGS (turquoise) and WES (salmon) coverage below the threshold for clinical reporting, with variants ranked by their corresponding values on the y-axis within the WGS and WES datasets. The mean percentage of individuals covered below 20X for WGS and WES is shown in the zoomed-in view. **d**, Vertical bars indicate the 1%–99% coverage range in WGS (turquoise) and WES (salmon), with variants ranked by the 1% coverage values within the WGS and WES datasets.

**Extended Data Fig. 8: Cases with protein-null phenotypes.**

**a**, Alignments in the *ITGB3* locus for a Glanzmann’s thrombasthenia case with a premature stop (blue bar) and a tandem repeat revealed by improperly mapped read pairs. **b**, Number of improperly mapped read pairs in the 9th intron of *ITGB3* in 6,656 samples sequenced by 150bp reads before (light grey dots) or after (dark grey squares) the data freeze. The Glanzmann’s thrombasthenia cases with the tandem repeat and with the SVA insertion, and the carrier mother of the latter, are highlighted. **c, d,** Alignments for the *ITGB3* locus for the Glanzmann’s thrombasthenia proband (**c**) and his mother (**d**) with a p.T456P variant for the proband (blue bar) and an insertion revealed by an excess of mapped reads for the 9th intron for the proband and his mother. **e** - upper panel, Long-read alignments for the PCR-amplified *ITGB3* DNA from the Glanzmann’s thrombasthenia proband covering the element with excess reads. Downstream Read Elements (DRE) starts are represented in the histogram. **e** - lower panel (from left to right), Graphic depiction of the Glanzmann’s thrombasthenia pedigree (A, proband, B mother and C grandmother) with the flow cytometric measurements of platelet GPIIbIIIa expression indicated as percentage of normal levels and genotypes; Confirmation of the insertion by gel electrophoresis of PCR products covering the insertion; Diagram of the inserted SVA (Alu, SINE-VNTR-Alu) retrotransposon element (insSVA). **f**, Alignments in the *RHAG* locus of the Rhesus-null case with a splice donor variant (blue bar) and a tandem duplication revealed by improperly mapped read pairs.

**Extended Data Fig. 9: Deletion of a *GATA1* enhancer and part of the *HDAC6* open reading frame and its effects.**

**a**, WGS reads show a hemizygous 4108 bp deletion (X:48,659,245-48,663,353) in the propositus. **b**, Sanger sequencing of PCR fragment with primers flanking the 4801 bp deletion (fragments shown in Fig. 5d). The red arrow points to the position of the fusion between bp 48,659,245 and bp 48,663,353. **c**, Electron microscopy images show enlarged more rounded platelets with the presence of abnormal semi-circular empty vacuoles in some platelets (*) and platelets depleted of alpha granules for the propositus compared to platelets from an unrelated healthy control. Marker is 1.5 𝝻M. **d, e**, Electron microscopy image analysis (n=20 platelets/condition) for platelet size (µm2) and alpha granules (per platelet adjusted for the size) using ImageJ software. *P*-values for both conditions are *P*< 0.0001 using unpaired *t*-test. The values for the platelet area and number of alpha granules shown in the propositus (P) are very similar to the defects described for *GATA1* deficiency (PMID:28082341). **f**, Platelet spreading analysis using SIM (Z-stacks) and staining for F-actin (red) and acetylated 𝛂-tubulin (green). Washed platelets were spread on fibrinogen for 0 (basal condition), 30 and 60 minutes for control (C), father (F), mother (M) and propositus (P). Representative images are shown. Marker is 1.5 𝝻M. **g**, Platelet analysis using structured illumination microscopy (SIM) and staining for acetylated 𝛂-tubulin (green) before spreading (time point 0). The microtubule marginal bands are clearly disturbed and hyper-acetylated for non-activated platelets of the propositus while being normal for the father (f) and mother (m). Marker is 1.5 𝝻M. **h**, Quantification of mean area size for platelets stained with F-actin was performed for 5 images of the different conditions using ImageJ. Platelet spreading on fibrinogen was increased for the propositus when compared to the parents and unrelated control. The means and SDs are shown. Statistically significant differences in means, assessed using one-way ANOVA (multiple comparisons), are shown with connecting lines. **i**, The upper panels are representative images for the control (c) and the propositus (p) where large MKs are present but proplatelet formation was strongly reduced. The lower panel shows the quantification of proplatelet formation by MK at day 12 of differentiation from cultures performed in duplicate for each individual. All MK with proplatelets were counted as positive in 10 images per condition. Results were analyzed by one-way ANOVA. **j**, Day 12 differentiated MKs for the indicated individuals were stained for F-actin (red) and HDAC6 (green). Upper two panels: HDAC6 is expressed in the cytosol and is trafficked to proplatelets as shown in MKs for the control and father (bold arrows). Middle two panels: MKs for the propositus show no HDAC6 expression while cultures from the mother contain a mixture of MKs that are positive and negative (15 of the 45 MKs) for HDAC6 expression. Lower two panels show only the HDAC6 staining. **k**, Day 12 differentiated MKs for the indicated individuals were stained for Acetylated 𝛂-tubulin (green). Highly organised tubulin structures are present in all MKs from the control and father while the patient (47 of the 57 MKs) and mother (16 of the 46 MKs) contain MKs that show signs of tubulin depolymerisation (*).

**Extended Data Fig. 10: Thrombocytopenia due to compound regulatory and coding rare variants in *MPL*.**

**a,** Top: Smoothed covariance between H3K27ac ChIP-seq and ATAC-seq (as per b) and coverage tracks generated by RedPop for activated CD4+ T-cells (aCD4), B, EB, MK MON and resting CD4+ T-cells (rCD4); Middle: MPL gene with exons in yellow; Bottom: positions of the deletion (blue bar) and SNV (blue dot) in the proband. **b**, Pedigree for the proband (P) with thrombocytopenia due to a 454bp deletion encompassing exon 10 of *MPL*, which was inherited from the mother (M), and an SNV just upstream of the 5’ UTR of *MPL*. **c**, Sanger sequencing traces confirming the presence of the heterozygous SNV in P and its absence in M. **d**, Gel electrophoresis of PCR amplicons covering the deletion confirming presence of the deletion in P and M. **e**, Mean fluorescence intensities (MFI) on the y-axis obtained by the flow cytometric measurement of MPL abundance (CD110) on the membrane of platelets from unrelated healthy controls (C), M and P. The MFI was normalised with unstained. **f**, Results of luciferase reporter assays in K562 cells expressed with empty pGL3 vector or after cloning with an MPL promoter fragment containing the wild type G allele (MPL-SNV-G) or the variant A allele (MPL-SNV-A). The measurements were derived from four independent transfection experiments. Statistically significant differences in means, assessed using one-way ANOVA adjusted for multiple comparisons, are shown with connecting lines.
