## Supplementary material for "Whole-genome sequencing of rare disease patients in a national healthcare system": Legends for Supplementary Tables

**Supplementary Table 1 – Enrolment and Rare Disease Domains.**

Enrolment by Hospital: The total number of samples (following QC) enrolled for each rare disease domain, broken down by hospital and geographic location. Hospitals are listed in descending order of total number of participants recruited over all the domains.

Domain metrics: For each domain, the following is shown shown: the degree of pre-screening, whether phenotype data were available, the number of QC-passing samples which have undergone WGS, the number of individuals who belong to the maximally unrelated set, the number of probands, the percentage female individuals, the percentage of individuals affected by the phenotype for their domain, the percentage of individuals without close relatives in the study and the percentage of individuals of European ancestry. The metrics for the entire collection are shown in the last row.

NPD – Diagnostic Criteria: A summary of the neuropathic pain disorders and the key diagnostic criteria used to classify the study participants enrolled. Genes (including the OMIM reference) and pattern of inheritance for the Mendelian inherited pain disorders are listed. The variants of genes implicated in small fibre neuropathy are relatively common and represent a gene/environment interaction rather than a Mendelian pain disorder.

NPD – Outcome Measures: A summary of the outcome measures used to classify and grade neuropathic pain of the study participants.

**Supplementary Table 2 – Pertinent Findings.**

Diagnostic-grade genes: List of the diagnostic-grade genes per disease domain, and including gene name and symbol (HGNC), associated disorder and mode of inheritance (OMIM), RefSeq and Ensembl transcripts selected for reporting and LRG identifiers, as well as OMIM identifier for the gene.

SNV and indel list: List of the reported pathogenic and likely pathogenic SNV and indels, including position in GRCh37, reference and alternative alleles, consequence according to VEP, genotype, MDT outcome (pathogenicity), HGVS annotations for cDNA and protein, minor allele frequency (MAF) in gnomAD, whole NIHR BioResource dataset, and unrelated NIHR BioResource dataset, WES coverage (according to ExAC), and novel status (absent from HGMD Pro 2018.1, and not likely pathogenic/pathogenic in ClinVar unless also labelled as benign). Note that we have recently relabelled the variant encoding Arg698His in *ABCB11* as having uncertain significance following functional studies.

Large deletion list: List of the reported pathogenic and likely pathogenic large deletions, including position in GRCh37, genotype, length in base pairs and number of genes, affected diagnostic-grade gene, MDT outcome (pathogenicity), and confirmed status.

**Supplementary Table 3 – Genetic association and regulome analysis.**
Phenotypic Tags: List of the phenotypic tags used for the different rare disease domains in the BeviMed genetic association analysis.

BeviMed Associations UK Tags: Genes with a BeviMed posterior probability > 0.75 of being associated with one of 28 phenotypic tags. For each gene is included the phenotypic tag, the main domain for that tag, the posterior probability of association, the modal model, the genomic region, the total number of modelled variants, the posterior expected number of cases explained by the variants, the posterior expected number of explanatory variants, whether the genetic association is unconfirmed or established as prior to or since 2015 and, in the latter case, the Pubmed ID of the key publication leading to DGG status for the gene and phenotype.

BeviMed Associations UK Biobank: Genes with a BeviMed posterior probability > 0.4 of being associated with either the left, right or merged tails of the red blood cell score. Individuals in the left/right tail have a low/high mean corpuscular volume (MCV) and high/low red blood cell count (RBC#). For each gene is included its position, BeviMed posterior probability of association, function, associated rare disorder (if any), genome wide association signals for blood cell indices, description of the phenotype in a murine model (if any), and the potential plausibility. Platelet cell type: MPV: Mean platelet volume; this is the mean volume of platelets. PDW: Platelet distribution width; this is the spread of the platelet volume distribution. PCT: Plateletcrit; this is the volume fraction of blood occupied by platelets. Mature red cell type: MCH: Mean corpuscular hemoglobin; this is the average mass of haemoglobin per red cell. MSCV: mean sphered red cell volume; mean volume of sphered red cells. HGB: haemoglobin concentration; this is the concentration of haemoglobin with respect to unit of volume of blood. RDW: red cell distribution width; this is coefficient of variation of red cell volume distribution. Immature red cell type: RET#: Reticulocyte count; this is the count of reticulocytes per unit volume of blood. RET%: Reticulocyte fraction of red cells; this is the percentage of red blood cells that are reticulocytes. IRF: Immature fraction of reticulocytes; this is the fraction of reticulocytes with high RNA content, as measured by light scatter. MRV: Mean reticulocyte volume; this is the mean volume of reticulocytes. HLSR#: High light scatter reticulocyte count; this is the count of high RNA content (immature) reticulocytes per unit volume of blood. HLSR%: High light scatter reticulocyte percentage of red cells; this is the immature reticulocyte count as a percentage of red blood cell count. Leukocyte cell type: EO#: Eosinophil count; this is the count of eosinophils per unit volume of blood. EO%: Eosinophil percentage of white cells; this is the percentage of white cells that are eosinophils. BeviMed BeviMed Variants UK Biobank: the variants with a posterior probability of being pathogenic greater than 0.5 conditional on the genetic association models. CHROM: chromosome, POS: position on the GRCh37 reference genome, REF: reference allele, ALT: alternate allele, AC: allele count (note: all are carriers are heterozygous).

Regulome Cell Data: Details of the data sources used to construct the cell-type specific regulomes, the number of elements called in each regulome and the genomic sizes of the regulomes.
