## Supplementary Information for "Whole-genome sequencing of rare disease patients in a national healthcare system"

**Enrolment, research ethics and consent** 3

**Domain descriptions** 3

**Overview of the collection of participants** 8

**Clinical and laboratory phenotype data** 9

**DNA sequencing** 9

**Analysis of genotyping data** 10

Overview of data processing pipeline 10
Alignment and SNV/indel calling 11
Sample provenance checks 11
Variant quality checks at the sample level 12
Coverage assessment 12
Computation of sex chromosome karyotypes 12
Ancestry and relatedness estimation 13
Variant normalisation and loading of variants into HBase 15
SNV and indel annotations 15
SNV and indel quality control 16
Concordance in duplicates and twins 16
Overview of SNVs and indels in the dataset 17

**Integrative Variant Analysis (IVA) web application** 18

**Large deletion calling and quality control** 18

**Clinical reporting** 21

Gene lists and transcript selection 21
Variant filtering 22
Web application for variant assessment 22
Variant interpretation in MDT meetings 22
Sensitivity of WGS to detect known pathogenic variants 23
Overview of diagnostic results 24
Upload of reported variants to public repositories 26
Reported variants informing changes in clinical management 26

**Structural variants in patients with null protein phenotypes** 27

**Genetic association between rare variants and rare diseases** 28

Statistical approach 28
Case/control groupings using phenotypic tags 29

Selection of variants affecting transcript sequences 29
Post-processing of BeviMed results 30
BeviMed results for genes 31
Observed false positive rate 31
Positive predictive value 32
Estimated false positive rate 33

**Polygenic and rare variant associations with phenotypic extremes in the UK Biobank cohort 33**
Selection of participants 33
Polygenic contribution to tail status 34
Whole genome sequencing GWAS 35
BeviMed analysis 35

**Matchmaker Exchange 36**

**Regulome analysis 36**
Ethics 36
Definition of cell type specific regulomes 37

**Regulatory element detection using patterns of peaks (RedPop) 38**
Transcription factor binding peak calling 39
Identifying possibly causal deletions of elements regulating diagnostic-grade genes 39
Deletion of GATA1 enhancer and HDAC6 open reading frame 39
Details on the functional analysis of the GATA1 enhancer/HDAC6 deletion 41
Homozygous deletion of CTCF binding sites in the first intron of LRBA 42

**Identifying causal SNVs in elements regulating diagnostic-grade genes 42**
An SNV in the promoter of MPL, combined with deletion of exon 10 of MPL 42
Details of the luciferase reporter assay 43

**Alternative variant datasets for versions 37 and 38 of the human reference genome 44**

**Appendix 44**
Appendix 1: Neuropathic pain disorders 44
Appendix 2: Extreme red cell traits in UK Biobank 45

**References 49**

**Enrolment, research ethics and consent**

As part of a pilot study for the 100,000 Genomes Project, the NIHR BioResource (NBR) enrolled a group of 9,742 study participants comprising patients with rare diseases and some of their close relatives. Participants were enrolled into one of 15 rare disease domains approved by the NBR's Sequencing and Informatics Committee. Each domain corresponded to a group of clinically related rare diseases and their recruitment criteria covered pathologies of various body systems. The domains adopted differing approaches to recruitment in order to address particular research objectives (see below, Domain descriptions). Enrolment took place between December 2012 and March 2017. In addition to the 15 rare disease domains recruited by the NBR, the study included three additional domains: a second rare diseases pilot study coordinated by Genomics England Ltd (GEL), a domain comprising a set of control samples and a domain comprising samples from UK Biobank[1]. Consequently, there were 18 domains in total.

The enrolment of NBR participants was coordinated by the University of Cambridge. Although most participants were recruited at NHS Hospitals in the UK, some recruitment occurred at overseas hospitals (**Supplementary Table 1 (Enrolment by hospital)**, **Extended Data Figure 1a**). All 13,187 participants provided written informed consent, either under the East of England Cambridge South national research ethics committee (REC) reference 13/EE/0325 or under local institutional review board approval and governance. Obtaining consent for overseas samples was the responsibility of the respective principal investigators at the enrolling hospitals. The NBR retained de-identified versions of the consent forms from overseas participants and a material transfer agreement was applied to regulate the exchange of samples and data between the donor institutions and the University of Cambridge.

**Domain descriptions**

**Supplementary Table 1 (Domain metrics)** contains quantitative summaries of each domain, including the distributions of participants by sex, ethnicity and phenotype, while in this section we set out a brief description of each domain in prose.

<u>BPD</u> (Bleeding, Thrombotic and Platelet Disorders)**.** Participants in this domain were enrolled at 31 hospitals. The probands had one of: a platelet disorder (with or without pathological bleeding); a pathological bleeding disorder not explained by a platelet function defect; a coagulation factor defect; or multiple thrombotic events at a young age. To enrich for inherited causes, probands with a positive family history and/or early onset and/or presence of syndromic features were enrolled preferentially. Where participants had informative pedigrees, affected and unaffected individuals were recruited if possible. The clinical characteristics of a subset of the participants in this domain have been reported previously[2]. With some exceptions, patients with a BPD of known molecular aetiology (e.g. Haemophilia A and B) were excluded from enrolment.

<u>CNTRL</u> (Technical Controls). A set of DNA control samples from consented participants: 27 healthy individuals from the NBR were enrolled under the REC-approved study BLUEPRINT

(REC 12/EE/0040), 5 healthy individuals enrolled under the REC-approved study GENES & PLATELETS Healthy (REC 10/H0304/65) and 18 patients (9 individuals with BMI > 40 and 9 individuals with partial lipodystrophy) collected under the REC-approved study Inherited platelet conditions (BPD) (REC 10/H0304/66). These samples were recruited primarily in order to test our laboratory procedures and our data transfer and analysis pipelines.

CSVD (Cerebral Small Vessel Disease). The participants in this domain were enrolled at 10 hospitals. Patients were eligible if they were suspected to have familial CSVD, based on clinical features such as lacunar stroke or cognitive impairment, at an early age (typically < 60 years). Evidence consistent with SVD from magnetic resonance imaging (MRI) of the brain, such as lacunar infarcts and white matter hyperintensities, was also required. Data on features typical of monogenic SVD were collected, including clinical history of migraines, encephalopathy and psychiatric disturbance. Clinical information and MRI results for all recruits were assessed by consultant neurologists and databases of previous stroke admissions were screened. Individuals with causal variants in *NOTCH3* were excluded from enrolment.

EDS (Ehlers-Danlos and Ehlers-Danlos-like Syndromes). The participants in this domain were enrolled at five hospitals. Patients with EDS subtypes of known molecular aetiology were excluded from enrolment. Patients met the Villefranche criteria for hypermobility EDS[3]. Probands had generalised joint hypermobility, skin hyperextensibility, connective tissue fragility and chronic pain. At the time of recruitment, patients had previously undergone ophthalmological and cardiac assessments to exclude Marfan syndrome and other non-EDS hereditary disorders of connective tissue. The participants had a family history suggestive of autosomal dominant inheritance. Where possible, close relatives were also recruited.

GEL (100,000 Genomes Project–Rare Diseases Pilot).The main purpose of this domain was to pilot the recruitment processes for the 100,000 Genomes Project and to assess the ability of National Health Service (NHS) hospitals to enrol rare disease patients and their close relatives, under the governance framework of the NHS. Enrolment was coordinated by GEL in partnership with the NBR. The recruitment strategy differed from that of the NBR domains by enrolling trios and pedigrees when possible. Patients exhibiting clinical symptoms of one of 161 rare inherited disorders and their close relatives (preferably both parents) were enrolled at eight NHS hospitals in England. Patients with known causal mutations were not eligible. Detailed clinical characteristics were collected on the participants using the human phenotyping ontology (HPO). All participants will be followed over their life course using electronic health data from primary care (general practitioners), secondary care (hospitals) and relevant registries. A full analysis of this domain will be published in due course by GEL. For the present study, data from this domain were used principally to increase the number of WGS genomes from genetically independent individuals, thus increasing the power of genetic association analyses (see below). Otherwise, only the binary affection status of these participants were available to researchers working on this study.

HCM (Hypertrophic Cardiomyopathy). The participants in this domain had an unequivocal diagnosis of HCM made at one of four UK NHS centres providing specialist HCM care. HCM is characterised by primary left ventricular hypertrophy and has an estimated population prevalence of approximately 1 in 500. It is the leading cause of arrhythmia and sudden death in athletes and young adults aged under 35 years. All participants were diagnosed below the age of 70, or were affected relatives. To qualify for enrolment patients had to have undergone clinical genetic testing on an HCM gene panel, with no pathogenic mutation identified.

ICP (Intrahepatic Cholestasis of Pregnancy). Severe ICP is defined as gestational pruritus in association with maternal serum bile acids ≥ 40 µmol/L. It is associated with adverse pregnancy outcomes, including spontaneous preterm labour, fetal asphyxia and intrauterine death. ICP typically presents in the third trimester. Affected women and their relatives are at increased risk of biliary disease in later life, e.g. drug-induced cholestasis, gallstones, and hepatic fibrosis. The ICP domain comprises women with disease onset before 32 completed weeks of gestation, which affects approximately 1 in 3,000 pregnant women in the UK. Individual cases with severe, early onset ICP were recruited from 14 UK consultant-led antenatal NHS clinics and from three international units in Argentina, Australia and Sweden. Women were excluded from the study if they had other known causes of hepatic dysfunction such as haemolysis, elevated liver enzymes and low platelets (HELLP) syndrome, preeclampsia, acute fatty liver of pregnancy, acute viral hepatitis, confirmed primary biliary cholangitis or any cause of biliary obstruction on ultrasound. No genetic pre-screening for causal variants in known genes was applied before enrolment.

IRD (Inherited Retinal Disorders). IRD describes a phenotypically heterogeneous group of conditions consequent upon dysfunction and/or degeneration of the neural retina or retinal pigment epithelium, resulting in visual impairment. It is the most common cause of severe visual impairment. The participants were enrolled at five NHS hospitals, most at the Moorfields Eye Hospital, London. Most individuals had undergone some previous genetic testing using routine diagnostic approaches. The analysis of the WGS of a fraction of the participants in this domain has been reported previously[4].

LHON (Leber Hereditary Optic Neuropathy). LHON causes subacute sequential bilateral visual loss which is usually irreversible. Participants with a diagnosis of LHON and a positive test for one of three mitochondrial DNA (mtDNA) mutations in Europeans (m.11778G>A, m.3460A>G, m.81440T>C) were enrolled in this domain. These mutations are found in ~1 in 300 of the UK population, but the prevalence of blindness due to LHON mtDNA mutations is approximately 1 in 40,000. Environmental factors undoubtedly influence the clinical penetrance, but segregation analyses implicate an interaction between a nuclear modifier locus and the mtDNA mutation. The LHON domain was included to test the hypothesis that the blindness only occurs when both the nuclear and mtDNA risk alleles are present in the same individual.

<u>MPMT</u> (Multiple Primary Malignant Tumours). Participants in this domain were mostly (> 95%) recruited in the UK, through the NHS regional clinical genetics services. In each kindred there was a clinical suspicion of a cancer predisposition syndrome, but routine genetic assessment/testing had not identified a genetic cause at the time of enrolment. Most (95%) of cases analysed had developed MPMT (defined as ≥ 2 primaries by age 60 or ≥ 3 by 70) but a minority had developed a single primary and had a first-degree relative with MPMT. Tumours in the same tissue type and organ were considered separate primaries if, in the case of paired organs, they occurred bilaterally or if the medical record clearly denoted them as separate. The International Agency for Research on Cancer criteria for defining separate primaries were also used[5]. For 95% of pedigrees, only the DNA of a single family member was sequenced.

<u>NDD</u> (Neurological and Developmental Disorders). The participants in this domain were adults and children with a previously undiagnosed neurological condition. Referrals were from clinical specialists in paediatrics, adult neurology or clinical genetics. The cases enrolled had early or severe epilepsy, encephalopathy, dystonia, spasticity, intellectual disability or metabolic disorders. In many cases, pre-screening for known genetic aetiologies was performed but this was not systematic. The majority were singleton cases but some trios of proband, mother and father were included for paediatric conditions.

<u>NPD</u> (Neuropathic Pain Disorders). The NPD domain enrolled individuals with extreme neuropathic pain phenotypes (both sensory loss and gain) at secondary care clinics located in six NHS hospitals. In order to be recruited, cases had to be over 18 years of age with a history of life-style altering sensory disorder (either pain or loss of sensation) for greater than three months. Patients with a known underlying genetic cause of chronic pain (e.g. Fabry's disease [ORPHA:324] and *SCN9A* erythromelalgia [ORPHA:1956]), intellectual disability and/or autistic spectrum disorder so severe that they could neither give consent nor engage in additional pain phenotyping were excluded. The Neuropathic Pain Special Interest Group of the International Association for the Study of Pain grading for neuropathic pain was used to assess the participants[6]. Further details are available in **Appendix 1: Neuropathic pain disorders** and **Supplementary Table 1 (NPD Criteria – Diagnostic criteria, NPD Criteria – Outcome measures)**.

<u>PAH</u> (Pulmonary Arterial Hypertension). In PAH, adverse remodelling of the pulmonary vasculature causes narrowing and obliteration of the capillary arteries in the lung, resulting in elevated resting mean pulmonary artery pressure and right heart dysfunction. A mean pulmonary artery pressure of 25 mm Hg or above, with a pulmonary capillary wedge pressure of less than 15 mm Hg is indicative of PAH. The diagnosis of idiopathic PAH was based on the exclusion of other associated forms of PAH. Participants with idiopathic PAH were recruited at 10 NHS UK National Pulmonary Hypertension hospitals and four international hospitals. All recruits had a clinical diagnosis of idiopathic PAH, heritable PAH, drug-associated PAH, or pulmonary veno-occlusive disease/pulmonary capillary haemangiomatosis (PVOD/PCH) established by their expert centre. Analyses of this domain have been recently reported[7,8,9]. No

systematic genetic pre-screening for causal variants in previously established causal genes was applied before enrolment.

PID (Primary Immune Disorders). Cases were recruited by specialists in clinical immunology (either trained in paediatrics or internal medicine) at 21 NHS hospitals in the UK and a small number of hospitals in the Netherlands. The eligibility criteria included the following: clinical diagnosis of common variable immunodeficiency disorder (CVID) according to internationally established criteria[10], extreme autoimmunity, or recurrent (and/or unusual) severe infections suggestive of defective innate or cell-mediated immunity. Patients with known secondary immunodeficiency (e.g. due to cancer or HIV infection) were excluded. Genetic screening for known causes of PID prior to enrolment was encouraged but not applied systematically. Within a broad range of phenotypes, CVID is the most common disease category, comprising 46% of the cases. Participants in this domain consisted predominantly of singleton cases, but the DNA samples from additional affected and/or unaffected relatives of some of the patients were also sequenced. A manuscript describing analyses of this domain is under review (Thaventhiran *et al*, under review).

PMG (Primary Membranoproliferative Glomerulonephritis). PMG refers to kidney disease in which a biopsy shows increased glomerular mesangial matrix and cellularity with thickening of the capillary walls and there is an absence of an underlying infectious, neoplastic or autoimmune disorder. Participants in this domain were enrolled at 10 NHS paediatric renal units in the UK (64 patients) and 18 NHS adult renal centres (120 participants, of whom 21 had paediatric onset of disease). IC-PMG refers to PMG where there is deposition of immunoglobulins and complement C3 in the glomeruli, and 'C3 glomerulopathy' (C3G) is where there is C3 without significant immunoglobulins deposited. The C3G category is further subdivided by electron microscopic appearance into C3 glomerulonephritis (C3GN) and dense deposit disease (DDD). PMG has an estimated incidence of 3–5 in 1 million and has a poor renal prognosis[11]. Where available, kidney biopsies were reviewed centrally to confirm classification into IC-PMG, C3GN or DDD. No genetic pre-screening for causal variants in known genes was applied before enrolment.

SMD (Stem Cell and Myeloid Disorders). Cases were enrolled into this domain if they presented with an inherited bone marrow failure of unknown molecular aetiology in childhood or an inherited cytopenia (including erythroid lineage) for which non-genetic and autoimmune aetiologies could be excluded. In addition, patients presenting with a myeloproliferative phenotype and a positive family history of a related haematological disorder were eligible for enrolment. Recruitment into the domain was done by paediatric and adult haematologists from NIHR/National Cancer Research Network Primary Treatment Centres. In addition, cases and their close relatives were enrolled at centres in Canada, Denmark, Egypt, Israel, Norway, Sri Lanka, South Africa, Sweden, Turkey and the USA. The DNA of most probands had undergone previous genetic analysis using targeted sequencing with phenotype-specific gene panel tests and no pathogenic variants were identified.

SRNS (Steroid Resistant Nephrotic Syndrome). The prevalence of SRNS is estimated at 2–4 in 100,000 people. There are two major clinical subsets: primary SRNS, defined as no response to high dose steroid at four weeks in children or four months in adults, and secondary SRNS, where initial steroid sensitivity is lost and treatment resistance evolves as a secondary event either rapidly or over time. Participants in this domain were enrolled through the UK NephroS study at tertiary NHS paediatric nephrology centres and adult renal units across the UK. Patients with either sporadic or familial SRNS were eligible for enrolment. Cases with SRNS secondary to systemic disease or obesity were excluded from enrolment. The majority of patients had a histological renal biopsy diagnosis that allowed a second tier of stratification based on histology as well as drug response. The majority (70%) of the cases had undergone previous genetic testing by whole-exome sequencing and tested negative for causal variants in diagnostic-grade SRNS genes.

UKB (UK Biobank – Extreme Red Cell Trait Score). The UK Biobank is a biomedical cohort of approximately half a million participants recruited in the UK between 2006 and 2010[1]. The participants, 54% of whom are women, were aged between 37 and 73 years at their date of recruitment. Each participant underwent a baseline assessment at one of 21 centres distributed across Great Britain, during which EDTA anticoagulated blood was collected for full blood count (FBC) analysis[12]. A subset of UK Biobank participants likely to carry rare alleles modulating erythropoiesis or red cell survival/clearance mechanisms were selected for WGS by considering a univariate quantitative phenotype derived from the FBC-measured MCV and RBC# values. 383 and 381 participants from the extreme left and extreme right tails of the quantitative distribution passed quality control (QC) after being successfully sequenced. Further details of the approach used for selecting samples are given in **Appendix 2: Extreme red cell traits in UK Biobank**.

**Overview of the collection of participants**

Sex (derived from X and Y chromosome karyotypes), ethnicity and relatedness for all samples were inferred computionally from WGS data (as detailed below). The age of the participant on the date his/her DNA sample was collected was available for 97.9% of the 13,037 samples that underwent WGS and passed QC (see below). These variables highlight some of the differences in the disease presentation and the recruitment approaches between the domains (**Extended Data Figure 1b, Supplementary Table 1 (Domain metrics)**). The ICP domain comprised only female participants because the phenotype is related to pregnancy. The proportion of families for which WGS data were only generated for the proband was > 50% for all domains except GEL, CSVD and SMD, which largely comprised parent-child trios and mother-child duos. The overall ethnic breakdown for the study collection closely matched that of the 2011 UK census[13], suggesting equality of access (**Supplementary Table 1 (Domain metrics)**). Age at sampling was generally early in life for recruits with a paediatric onset (NDD, SMD), at childbearing age for ICP recruits, after the age of 50 for UKB recruits (due to UKB's enrolment criteria), and throughout adulthood for the other disorders (**Extended Data Figure 1b**).

**Clinical and laboratory phenotype data**

Staff at the hospitals responsible for enrolment were provided with the eligibility criteria for the domains for which they were recruiting patients. Different levels of pre-screening to exclude previously known disorders were used across the 15 domains. ICP, PAH, and PMG had virtually no genetic pre-screening. BPD, BPD, CSVD, EDS, IRD, MPMT, NDD, NPD, PID, SMD and SRNS had some pre-screening based on clinical presentation and the results of non-DNA based laboratory and imaging tests. HCM and LHON had extensive genetic screening in the relevant diagnostic-grade genes. The clinical and laboratory phenotype data were captured through case report forms (CRF) by paper questionnaires or by online CRF data capture applications and deposited in the NBR study database managed by the University of Cambridge. A web-based phenotype capture system allowed the reporting clinical staff to record Human Phenotype Ontology (HPO) terms[14,2,15] at the enrolling hospitals. Additional phenotypic information recorded on paper questionnaires were transformed into HPO terms by the NBR study coordination office. Free text entries were transformed into HPO terms where feasible. An overview of the HPO data obtained for the 15 domains is depicted in **Extended Data Figures 1c and 1d**.

**DNA sequencing**

DNA from 58.8% of enrolled participants was extracted at the receiving hospital or research laboratory and sent to Cambridge for plating. The DNA for the remaining samples was extracted from EDTA-treated whole bloods at a central DNA extraction and QC laboratory in Cambridge. Samples were tested for adequate DNA concentration (Picogreen), DNA degradation (gel electrophoresis) and purity (OD 260/280 QC(Trinean)) before selection for WGS. DNA samples were prepared at a minimum concentration of 30 ng/µl in 110 µl, visually inspected for degradation and had to have an OD 260/280 between 1.75 and 2.04. They were then prepared in batches of 96 and shipped on dry ice to the sequencing provider (Illumina Inc, Great Chesterford, UK).

Illumina measured the DNA concentration of each sample they received. A sample was excluded from WGS if its corresponding measurement was < 30 ng/µl. They also genotyped each sample using an Illumina Infinium Human Core Exome microarray. A sample was excluded from WGS if its genotyping call rate was < 0.99. Illumina also measured contamination by other human genomes by computing the fraction of homozygous alternate sites with at least one high quality read that disagrees with that variant call. Samples with a contamination measure > 0.05 were excluded. Finally, samples for which there was a mismatch between genetically inferred and clinician reported sex that could not be resolved by further investigation, or for which consent had been withdrawn, were also excluded from WGS. In total, 59 DNA samples sent to Illumina were excluded from WGS. After WGS, comparisons of the microarray genotyping and corresponding WGS data did not reveal any discrepancies.

To prepare each DNA sample for sequencing, 0.5 µg of DNA was fragmented using Covaris LE220 (Covaris Inc., Woburn, MA, USA) to obtain fragments with a mean length of 450 base pairs (bp). The fragmented DNA samples were processed using the Illumina TruSeq DNA

PCR-Free Sample Preparation kit (Illumina Inc., San Diego, CA, USA) on the Hamilton Microlab Star (Hamilton Robotics, Inc, Reno, NV, USA). The concentrations of the final libraries were measured using the Roche LightCycler 480 II (Roche Diagnostics Corporation, Indianapolis, IN, USA) with KAPA Library Quantification Kits (Kapa Biosystems, Inc, Wilmington, MA, USA).

Over the course of the project (February 2014 to June 2017) samples were sequenced at three different read lengths: 100bp (377 samples), 125bp (3,154 samples) and 150bp (9,656 samples). Samples sequenced with 100bp and 125bp reads utilised three and two lanes of an Illumina HiSeq 2500 instrument, respectively. Samples sequenced with 150bp reads utilised a single lane of a HiSeq X instrument. The changes in read length over the course of the project were imposed by the sequencing service provider. At least 95% of the autosomal genome had to be covered at 15X and a maximum of 5% of insert sizes had to be less than twice the read length. Following sample and data QC at Illumina, 13,187 sets of WGS data files were received at the University of Cambridge High Performance Computing Service (HPC) for further QC.

**Analysis of genotyping data**

*Overview of data processing pipeline*

WGS data for 13,187 samples were sent to Cambridge University by Illumina. There, they underwent a series of processing steps (**Extended Data Figure 2**), which are summarised in this paragraph and described in detail in the following sections. We estimated the karyotypes of the sex chromosomes and computed pairwise kinship coefficients between samples. This information was used to identify repeat participant submissions and DNA sample swaps. Additionally, four further QC procedures were applied to ensure the single nucleotide variant (SNV) and short (≤ 50bp) insertion/deletion (indel) calls for each sample were of a high standard. In total, the sequence data of 150 samples (1.1%) were removed, leaving a dataset of 13,037 participants for downstream analysis. The 13,037 individuals were assigned to one of the following ethnicity clusters: European, African, South Asian, East Asian or Other. Pairwise kinship coefficients adjusted for population stratification were then estimated. These were used to identify networks of closely related participants and to identify a maximal set of 10,259 unrelated individuals.

The variants called in the 13,037 individuals were left-aligned and normalised with bcftools[16], loaded into an HBase database[17] and filtered according to thresholds on their overall pass rates (OPR; defined below). The sex chromosome karyotypes, the ethnicities and the relatedness estimates were used, along with enrolment information, to annotate the samples and variants. Participants were annotated with:

- affection status (affected/unaffected),
- family membership indexed by proband identifier,
- an indicator for membership of the maximal unrelated participant set,
- ethnicity and
- sex chromosome karyotype.

Variants were annotated with:

- CellBase consequence predictions,

- HGMD information where available,
- and population-specific allele frequencies.

*Alignment and SNV/indel calling*

Reads were aligned with the Illumina Isaac aligner version SAAC00776.15.01.27[18] to the reference human genome build GRCh37[19]. This produced BAM files[20] with mean sizes approximately 95GB for 100bp samples, 60GB for 125bp samples and 65GB for 150bp samples. SNVs and indels were called from all BAM files using the Illumina Starling software version 2.1.4.2[21]. The variants were called separately for each sample and encoded as both regular VCF and genome VCF (gVCF) files[22]. The gVCF file format stores information on read coverage and alignment quality in homozygous reference regions as well as variant positions, which allows a PASS filter to be applied to all positions in the genome for each sample.

*Sample provenance checks*

As a means of identifying duplicate samples, a set of 8,872 common single nucleotide polymorphisms (SNPs) was chosen at the start of the project with which to compute an approximate pairwise kinship coefficient between samples as the data arrived at the Cambridge HPC. The SNPs were a random subset of those typed by Roche on their targeted microarray platforms. The kinship coefficients were computed using PLINK[23] after each data delivery. More precise relatedness estimation was performed at a later stage (see below). Any kinship coefficient > 0.99 triggered an investigation to determine whether there had been an accidental duplicate enrolment, a laboratory duplication or an enrolment of monozygotic twins. Where a pair of samples resolved to the same individual, only the higher quality WGS data were retained, where quality was determined firstly by read length and secondly by autosomal callability (defined below); 124 duplicates and 1 triplicate were found. The 22 pairs of monozygotic twins we identified were retained in the resource. Eight unresolved duplicates were excluded. Most duplicate samples were explained by submission of the same patient by different consulting clinicians recruiting into the same domain. However, one duplicate spanned two domains because a patient with a phenotype consistent with the recruitment criteria for two domains was enrolled by two clinicians with different specialties.

Participant-declared and genetically-determined sex were compared for concordance following each WGS data delivery. For each sample, we counted the number of heterozygous, homozygous and hemizygous QC-passing SNVs in the non-pseudoautosomal region of the X chromosome. We defined the H-ratio as the number of heterozygous variants divided by the number of homozygous and hemizygous variants. We made initial sex calls based on the H-ratio as follows: individuals with H-ratio < 0.02 were declared putatively male and the others were declared putatively female (**Extended Data Figure 3e**). Comprehensive sex chromosome karyotyping was performed at a later stage (see below). Based on the initial sex calls, two samples with discordance between declared and estimated sex were excluded from further analysis.

*Variant quality checks at the sample level*

We performed the following variant quality checks:

- We computed the proportion of positions in the non-N regions of the reference autosomes that passed quality filters in the gVCFs. This proportion is called the autosomal callability. Samples for which fewer than 95% of bases in the reference genome passed were removed (**Extended Data Figure 3a**).
- We identified a set of common SNPs by downloading and merging the genome and exome data from gnomAD[24] and computing the minor allele frequencies (MAFs) in each ethnically homogeneous cluster of study participants (except for the South Asian cluster, which is absent from the gnomAD genome set; see below for details of clustering by ethnicity). The set of SNVs that had a MAF > 5% in each population was identified. We then determined the proportion of the positions of these SNVs in each gVCF that failed the variant caller's internal QC. Samples for which this proportion exceeded 0.55% were excluded (**Extended Data Figure 3b**).
- Transition-transversion ratios (Ts/Tv) are commonly used to determine SNV calling accuracy[25]. We calculated Ts/Tv ratios for all samples and observed that the distribution of these ratios varied depending on the read length; samples with a Ts/Tv ratio more than four times the read length specific interquartile range plus the read length specific median were excluded (**Extended Data Figure 3c**).
- We estimated the degree to which a DNA sample was contaminated by any other DNA sample using verifyBamID[26]. Samples with an estimate of contamination (FREEMIX) exceeding 3% were excluded (**Extended Data Figure 3d**).

In total, 14 samples were excluded due to these four genotyping QCs.

*Coverage assessment*

The mean (over samples) of the mean autosomal coverage by unique reads was 41.4, 37.9 and 35.3, and the mean 10th percentile of coverage was 31.0, 25.7 and 26.2 for the 100bp, 125bp and 150bp read length batches, respectively (**Figure 1b**). The mean coverage was greater than 30X in all samples and 90% of the reference genome was covered at least to 19X in all samples. In addition, all samples were covered to at least 15X in at least 95% of the reference autosomes. The average read duplication rate was 1.5% for read pair inserts in the HiSeq 2500 (100bp and 125bp read lengths). However the rate was 18% for the HiSeq X-generated 150bp data, which meant that extra reads were required to make up a similar level of coverage.

*Computation of sex chromosome karyotypes*

Once the WGS dataset was complete, we estimated the sex chromosome karyotypes using the BAM and VCF files. For each sample and chromosome, we normalised the number of aligned reads by dividing it by the length of the chromosome in the reference genome, excluding positions with an unknown base. The X/Auto and Y/Auto ratios were defined as the normalised read counts on X and Y divided by the median of the normalised read counts on the autosomes (Auto). The median was used because it is robust to large copy number alterations. For the putative males and putative females (defined based on the H-ratio, as described above), the mean and standard deviation of X/Auto and Y/Auto were computed (**Extended Data Figure 3f**).

Let us define these values as $mean_z$(X/Auto), $sd_z$(X/Auto), $mean_z$(Y/Auto) and $sd_z$(Y/Auto), where $z \in \{m,f\}$ denotes whether the summaries were conditioned on the putative males or the putative females. We defined the following four gates to classify individuals according to their specific X/Auto and Y/Auto values, denoted x and y, respectively:

- XY gate (shown in blue in **Extended Data Figure 3f**):
  - $mean_m$(X/Auto) - 10 $sd_m$(X/Auto) < x < $mean_m$(X/Auto) + 10 $sd_m$(X/Auto)
  - $mean_m$(Y/Auto) - 10 $sd_m$(Y/Auto) < y < $mean_m$(Y/Auto) + 10 $sd_m$(Y/Auto)
- XYY gate (shown in green in **Extended Data Figure 3f**):
  - $mean_m$(X/Auto) - 10 $sd_m$(X/Auto) < x < $mean_m$(X/Auto) + 10 $sd_m$(X/Auto)
  - y > $mean_m$(Y/Auto) + 10 $sd_m$(Y/Auto)
- XX gate (shown in red in **Extended Data Figure 3f**):
  - $mean_f$(X/Auto) - 10 $sd_f$(X/Auto) < x < $mean_f$(X/Auto) + 10 $sd_f$(X/Auto)
  - $mean_f$(Y/Auto) - 10 $sd_f$(Y/Auto) < y < $mean_f$(Y/Auto) + 10 $sd_f$(Y/Auto)
- XXY gate (shown in purple in **Extended Data Figure 3f**):
  - $mean_f$(X/Auto) - 5 $sd_f$(X/Auto) < x < $mean_f$(X/Auto) + 5 $sd_f$(X/Auto)
  - $mean_m$(Y/Auto) - 5 $sd_m$(Y/Auto) < y < $mean_m$(Y/Auto) + 5 $sd_m$(Y/Auto)
- XXX gate (shown in orange in **Extended Data Figure 3f**):
  - 1.5 $mean_f$(X/Auto) - 5 $sd_f$(X/Auto) < x < 1.5 $mean_f$(X/Auto) + 5 $sd_f$(X/Auto)
  - $mean_f$(Y/Auto) - 5 $sd_f$(Y/Auto) < y < $mean_f$(Y/Auto) + 5 $sd_f$(Y/Auto)

One participant had an H-ratio of 0.01 and was thus declared initially to be a male. However, the participant was located within the XX gate. The two X chromosomes were almost identical in sequence to one another, explaining the low H-ratio, and further investigation showed that this was due to consanguinity. This and other anomalies are labelled in **Extended Data Figure 3g**. Where permitted by the study ethics, we approached the referring clinician to confirm the non-standard sex chromosome karyotypes (n = 5). The observed frequencies of non-standard sex chromosome karyotypes, which were present in 13 of the 13,037 individuals (5 XXY, 4 XYY, 2 XXX, 2 XO), were comparable with those reported by a previous study[27]. The genotypes for the samples with non-standard sex chromosome karyotypes (except for XXX) were recalled to take into account the correct ploidy in the sex chromosomes (**Extended Data Figure 2**).

*Ancestry and relatedness estimation*

We estimated the degree of relatedness between participants and clustered the participants by their nuclear ancestry into the ethnicity categories European, African, East Asian, South Asian and Other, as follows.

We selected a set of SNPs for these analyses by starting from the 292,878 autosomal SNPs typed by three widely used Illumina genotyping arrays (HumanCoreExome-12v1.1, HumanCoreExome-24v1.0 and HumanOmni2.5-8v1.1). Presence of a SNP on all three arrays indicates that it can be reliably measured by microarrays, a good prior that it will be reliably called by WGS. To ensure this, we removed SNPs with a missing WGS genotype in at least one individual or with an overall pass rate below 0.99. We also removed SNPs at genomic positions in which more than two distinct alleles had been observed in the 1000 Genomes Phase 3 dataset[28] or the NBR dataset to ensure that all genotypes could be coded unambiguously as a count of the number of copies of the unique alternative allele carried by the individual. We then removed SNPs with a MAF < 30% in our dataset. Finally, we pruned the SNPs using PLINK v1.9[23] so that no pair of variants had an $r^2 \geq 0.2$. The final set contained 32,875 SNPs.

We implemented a pipeline that partitions a given set of participants into sets of related individuals and derives a maximal set of unrelated individuals. The inputs to the pipeline are a matrix of genotypes for the 32,875 SNPs encoded in a VCF file and a list of individuals with a high prior of being unrelated (the *reference set*). The pipeline uses PC-AiR[29] to perform a principal component analysis (PCA) on the standardised genotypes of the reference set and to project the standardised genotypes of the individuals outside the reference set onto the fitted principal components (PCs). The PC-AiR output object is then passed to the PC-Relate function to compute a kinship matrix that accounts for the population structure captured by the leading 20 PCs. Finally, this kinship matrix is passed to the PRIMUS[30] function to obtain clusters of related participants and to derive a maximal set of unrelated individuals on the basis that pairs of individuals with a kinship coefficient > 0.09 are related.

We used the pipeline firstly to obtain a set of unrelated non-admixed individuals within the 1000 Genomes Phase 3 data. To achieve this, we first excluded the 347 participants labelled as “Admixed American”. We then defined a reference set of 2,110 putatively unrelated individuals within the remaining 2,157 participants using the snpgdsIBDKING function from the SNPRelate R package[31,32]. We used the pipeline secondly to compute the relatedness amongst the 13,037 individuals in our dataset. In this second run, we merged the genotypes of the 13,037 individuals in our dataset with those of the 2,110 individuals from the 1000 Genomes data. We specified the 2,110 individuals as the reference set. The pipeline identified a maximal set of 10,259 unrelated individuals (**Supplementary Table 1 (Domain metrics)**) and a set of 9,244 pedigree networks ranging from singletons to a network of 15 were computed (**Figure 1d**).

We partitioned the final set of unrelated individuals into the equivalence classes: non-Finnish Europeans, Finns, Africans, South Asians and East Asians using their 1000 Genomes population code annotations. We modelled the score vectors of the leading five PCs by a multivariate normal distribution within each class of the partition and estimated the mean vector and covariance matrix for each population. Subsequently, we projected the genotypes from each of the 13,037 participants onto the vector space spanned by the leading five components of the 1000 Genomes PCAs (**Extended Data Figure 3h**). We computed the likelihood of the projected data under the five multivariate normal models estimated from the 1000 Genomes scores and classified each individual as belonging to the population corresponding to the model that yielded the highest likelihood, provided it was greater than $10^7$. If no such population existed, the individual was classified as “Other” (**Extended Data Figure 3i**). The resource is 82% European. However, approximately 7% of the participants are of South Asian ancestry, a population for which limited whole-genome sequencing (WGS) reference data are presently publicly available through resources such as gnomAD (**Extended Data Figure 3j**).

*Variant normalisation and loading of variants into HBase*

Variants in the gVCF files were processed using the bcftools norm command with the -cs option, which left aligns and nomalises indels and sets/fixes incorrect or missing reference alleles[33]. The variants were then transformed from gVCF format into Google *proto* format[34] and stored as binary *proto* objects spanning 1000bp of contiguous sequence in the reference genome using OpenCGA[35]. During the transformation, the anchoring reference base of insertions and deletions were removed. The *proto* objects of each sample were then copied from the respective *proto* file to the HBase[36] *Archive* table using the 'PUT' operation of the HBase application programming interface. The resulting *Archive* table held the *proto* objects by sample in columns and grouped by position in rows. Header information from the original gVCF files was stored for each sample. Any multiple variant calls overlapping the same position within an individual were resolved by only retaining the variant with the highest genotype quality. The list of participants corresponding to each of the genotype calls and whether these calls had a PASS filter value were recorded for different genomic positions in an HBase *Allele count* table using the 'APPEND' operation. As homozygous reference and PASS values were the most frequent values of genotype and filter data, we stored data as differences from these. The variants were then transferred from the HBase *Allele count* table to the HBase *Analysis* table.

*SNV and indel annotations*

All variants called across the 13,187 samples were annotated with deleteriousness scores and conservation scores and with cohort summary statistics including allele count, allele number, genotype count, minor allele frequency, HWE statistics (obtained using HTSJDK[37]), call rate, pass rate and overall pass rate (described below). Subsequently, using the RESTful annotation service provided by CellBase[38], the variants stored in the HBase Analysis table were annotated with consequence terms, HGVS descriptions, deleteriousness scores such as the combined annotation dependent depletion (CADD) score[39], SIFT[40] and PolyPhen-2[41], and conservation scores such as genomic evolutionary rate profiling (GERP)[42], PhastCons[43] and PhyloP[44]. The genotype and variant annotation data were then exported to AVRO files. These files were modified to add additional annotation from external reference datasets, including 1000 Genomes Project Phase 3[45], UK10K project (version 2016-05)[46], the ExAC / gnomAD project (version r2.0.2)[47], Trans-Omics for Precision Medicine Program (TOPMed, freeze5)[48] and the Human Genome Mutation Database (HGMD, version PRO 2018.1)[49] using Apache Spark[50]. Since the TOPMed data are only available with reference to GRCh38, these data were first mapped back onto GRCh37 using CrossMap[51].

*SNV and indel quality control*

The HBase database described above holds records of whether a genotype could be called reliably by Starling from the aligned read data at each position of the reference genome in each sample. The call rate for a position is the proportion of samples in which calling was possible. The pass rate for a position is the proportion of samples with a genotype called, for which the call passed QC. We defined the overall pass rate (OPR) as the call rate multiplied by the pass rate. Only female and male samples were used to calculate the OPRs for variants on the X and the Y chromosome, respectively.

We generated histograms of *P*-values from tests of departure from the null hypothesis of HWE for common SNPs and indels (MAF > 5%) amongst 8,511 unrelated Europeans over different intervals of OPR ((0.97,0.98], (0.98,0.99], (0.99,1]) and MAF ((5%,6%], (6%,7%], (7%,8%], (8%,9%], (9%,10%], (10%,20%], (20%, 30%], (30%, 40%], (40%, 50%]). Variants with OPR > 0.99 were found to have genotype proportions generally consistent with HWE in all MAF bins for SNVs, short insertions and short deletions (**Extended Data Figures 4a, 4b and 4c**). For variants with OPR in (0.98,0.99], we observed approximately double the number of *P*-values < 0.05 relative to that expected under the null, uniformly across MAF bins, suggesting that deviation from HWE was present in a small proportion of variants in this interval of OPR. We thus only released variants with an OPR > 0.99 through the variant browser (described below). However, for rare variant analyses, where maintaining high sensitivity is important and potentially pathogenic variants are routinely confirmed by Sanger sequencing, we used a threshold of OPR > 0.98.

*Concordance in duplicates and twins*

We selected pairs of samples from the 124 duplicated samples and from the 22 monozygous twins to evaluate the reproducibility of sequencing results. From all resolved duplicates and twins, we chose pairs where both samples were sequenced with the same read length and the participant was one of the 13,037 participants included in the final data release (one pair sequenced with 100bp reads, 17 pairs sequenced with 125bp reads and 88 pairs sequenced with 150bp reads).

For each sample we selected all the autosomal SNVs and small insertions and deletions with OPR ≥ 0.99 and for which the genotype was either heterozygote for the reference and principal alternative alleles, or homozygote for the principal alternative allele. **Extended Data Figure 4d** illustrates all possible combinations of genotypes in a pair of samples. We use $d_{ij}$ (in the cases that $i \neq j$) to denote the number of variants in which sample 1 has the ith genotype and sample 2 has the jth genotype, where i and j take values in {0,1,2,3} representing respectively the homozygote reference genotype, the heterozygote genotype for the reference and principal alternative alleles, the homozygote genotype for the principal alternative allele and any other genotypes. We use $c_{ii}$, where i takes values in {1,2} to denote the number of variants in which both samples have genotype i. We use c and d to denote the total numbers of variants in the cells corresponding to concordant ($c_{11}+c_{22}$) and discordant calls ($d_{01}+d_{02}+d_{10}+d_{12}+d_{13}+d_{20}+d_{21}+d_{23}+d_{31}+d_{32}$) respectively. The mutual non-reference concordance can be defined as (c / (c + d)) * 100%. The distribution of this statistic is shown in **Extended Data Figure 4e**.

The probability of observing a heterozygous variant in sample 2 given it is heterozygous in sample 1 is $c_{11} / (d_{10} + c_{11} + d_{12} + d_{13})$. The probability of observing a homozygous alternative variant in sample 2 given it is homozygous alternative in sample 1 is $c_{22} / (d_{20} + d_{21} + c_{22} + d_{23})$ (**Extended Data Figures 4f and 4g**). Symmetrical formulae apply when the sample ordering is reversed. Hence, for each pair of duplicates/twins there are two probabilities of each type, the smaller of which is shown in red and the greater of which is shown in blue in **Extended Data Figures 4f and 4g**.

*Overview of SNVs and indels in the dataset*

We called 172,005,610 variants with an OPR > 0.99, of which 157,411,228 (91.5%) were SNVs and 14,594,382 (8.5%) were indels. We annotated the variants with Sequence Ontology (SO) terms based on their predicted consequences with respect to transcripts in Ensembl 75. A single primary consequence was selected for each variant following a series of filtering steps. Firstly, only consequences associated with transcripts annotated with a biotype in the most biologically significant biotype group (ranked by protein coding, pseudogene, long non-coding and short non-coding)[52] were chosen. Secondly, only consequences associated with transcripts with the best source of curation were retained (ranked by HGNC, VEGA, Ensembl, UniProt and miRBase/RFAM). Thirdly, only consequences with the strongest Ensembl-determined impact were retained[53] (ranked by HIGH, MODERATE, LOW and MODIFIER). Fourthly, only consequences with the greatest severity were retained (ranked by the order provided by Ensembl53). Fifthly, only consequences associated with transcripts annotated as being canonical by Ensembl were retained, if any existed. Finally, the transcript with the longest coding/cDNA sequence length was chosen.

On the basis of primary consequences, 61.7% of the variants were labelled genic (SO:0001564, gene_variant), while the remainder were labelled intergenic (SO:0001628, intergenic_variant), regulatory (SO:0001566, regulatory_variant) or other. 80.0% of the genic variants were intronic (SO:0001627, intron_variant) and only 4.4% were exonic (SO:0001791, exon_variant) or affected a splice site (SO:0001568, splicing_variant). The ratio of coding (SO:0001968, coding_transcript_variant) to UTR (SO:0001622, UTR_variant) variants was 45:55 for SNVs and 21:79 for indels. 33.7% of the 1,853,212 coding SNVs were synonymous (SO:0001819, synonymous_variant) and 64.2% were missense (SO:0001583, missense_variant). 38,886 SNVs (2.1% of coding SNVs) led to a start loss (SO:0002012, start_lost), stop loss (SO:0001578, stop_lost) or stop gain (SO:0001587, stop_gained). The majority (62.1%) of the 67,746 coding indels introduced a frameshift (SO:0000865, frameshift), while 34.5% introduced an inframe insertion or deletion (SO:0001817, inframe) and only 2,287 indels (3.4% of coding indels) led to a start loss (SO:0002012, start_lost), stop loss (SO:0001578, stop_lost) or stop gain (SO:0001587, stop_gained) (**Extended Data Figure 5**).

48.6% and 59.2% of the SNVs and indels, respectively, were absent from the following databases: 1000 Genomes, UK10K, gnomAD, TOPMed (lifted back to GRCh37) and HGMD. 54.8% of the variants were observed in only one member of the maximal unrelated set. 82.6% of these variants were novel. 36.7% of the variants observed in two members of the maximal unrelated set were novel. A small proportion (1.90% of SNVs and 9.01% of indels) of the variants observed in more than two members of the maximal unrelated set were novel (**Figure 1e**). Thus, common variants are well represented in genetic databases but the majority of genetic variants are very rare and are mostly absent from the available databases.

In coding regions, indels that introduce frameshifts are, on average, selected against due to their deleterious effects on reproductive fitness[54], and our analysis corroborated this notion (**Extended Data Figure 3k**). In contrast, in non-coding regions of the genome, indel lengths were biased towards a multiple of two bases (**Extended Data Figure 3l**). We have made a similar observation in the gnomAD dataset (data not shown) to ensure this is not a technical artefact specific to our bioinformatic pipeline. The pattern is specific to repetitive regions of the genome (**Extended Data Figure 3m**), suggesting it can in part be attributed to tandem duplications.

**Integrative Variant Analysis (IVA) web application**
The SNVs and indels have been loaded into the IVA web application[55]. The data can be browsed freely online at[56].

**Large deletion calling and quality control**
Two software packages were used to call deletions larger than 50bp in each sample: Manta version 0.23.1[57] and Canvas version 1.1.0.5 (PMID: 27153601). Manta calls deletions on the basis of split read alignments and larger-than-expected insert sizes of read pair alignments. Canvas calls deletions on the basis of sustained reductions in coverage in a region of the reference genome. Manta is optimised for calling deletions of 50bp–10Kb while Canvas is optimised for calling deletions > 10Kb.

For each sample, deletion calls were merged across the two methods if the reciprocal overlap between them (the smaller of the two values obtained by dividing the length of the intersection of the two deletion calls by the length of each of the two calls) was > 0.7. Merged deletions were annotated with the Manta breakpoints as they are more precise than Canvas breakpoints. After merging, each deletion call in each sample was assigned to one of the three QC categories "Fail", "Possible" or "Confident". These categories are defined according to the rules below.
"*Fail*"
Any of the following criteria are met:

- the length of the call is > 50Mb;
- the genotype is heterozygous and the call is on the non-PAR of chromosome X in a male;
- the call is on the mitochondrial or the Y chromosome;
- the call was made by Canvas only and CANVAS_QUAL < 10;
- the call was made by Manta only and MANTA_FILTER is not "PASS".

"*Confident*"
The call does not meet the "Fail" criteria and none of the following additional criteria are met:

- > 70% of the call overlaps a flagged region (see below);
- the call was made by Canvas only and SNPS_DENSITY_HET_PASS > 0.5;
- the call was made by Canvas only and the number of Canvas calls in the sample is greater than the mean plus five times the standard deviation across all samples (as such

extreme values are probably due to generalised and artefactual non-uniformities in coverage leading to inflation in the number of false positive calls);
- the call was made by Canvas only, the sample had more than 50 Canvas deletions and the proportion of the genome which was deleted was greater than the 99.7% percentile across all samples;
- the call was made by Manta only and MANTA_QUAL < 20th batch-specific percentile;
- the call was made by Manta only and MANTA_GQ < 20th batch-specific percentile;
- the call was made by Manta only and MANTA_IMPRECISE = “TRUE”;
- the call was made by Manta only and length > 1Mb;
- the call was made by Manta only and DUKE0_START = “TRUE” or DUKE0_STOP = “TRUE”;
- the call was made by Manta only and SNPS_N_HET_PASS > 0;
- the call was made by Manta only and the genotype was heterozygous and READS_MEAN_MAPQ < 45.

The parameters in capitals above were generated by the Manta and Canvas software, except for DUKE0_START, DUKE0_STOP, SNPS_N_HET_PASS, SNPS_DENSITY_HET_PASS and READS_MEAN_MAPQ, which were generated using custom code. DUKE0_START and DUKE0_STOP evaluate to “TRUE” if a deletion call's start or end positions lie within 10bp of the breakpoints of a Duke non-unique region (see definition below); SNPS_N_HET_PASS is the number of heterozygous SNPs with OPR ≥ 0.99 within the boundaries of a deletion call; SNPS_DENSITY_HET_PASS is the ratio of SNPS_N_HET_PASS to the length of the deletion in Kb; READS_MEAN_MAPQ is the mean mapping quality of the read alignments within a deletion's boundaries. The flagged regions in the reference genome were the following:
- low mappability regions: DAC Blacklisted Regions (Encode, Accession: wgEncodeEH001432) and Human Mappability Blacklist (Encode, Accession: wgEncodeEH000322),
- centromeres[58],
- telomeres[58],
- IG loci (NCBI, Gene IDs: 50802, 3492, 3535),
- HLA loci (NCBI, Accession: NC_000006),
- segmental duplications[59,60],
- Duke non-unique regions (Encode, Accession: wgEncodeDukeMapabilityUniqueness35bp).

“*Possible*”
The deletion call does not meet either the “Fail” or the “Confident” criteria. By visual inspection of the reads around 100 deletion calls from the Possible and Confident deletion sets using Integrative Genomics Viewer (IGV)[61], we estimated the false call rates to be 24% and 10% respectively. To control the rate of false calls, whilst retaining calls in the Possible class which have a high chance of being real due to overlap with calls in the Confident set, any deletion in the Possible set which did not have a reciprocal overlap of at least 0.7 with any of the Confident deletions was removed.

In order to select a set of rare deletions for use in the downstream association analyses, the deletion calls required annotating with internal cohort allele frequencies. The deletion calls from all samples were partitioned into groups corresponding to distinct latent deletions. The partitioning was performed as follows. First, we used the ‘hclust’ function in R to construct a dendrogram of all deletion calls. One minus the reciprocal overlap was treated as the distance between each pair of calls, and the complete linkage option was used (i.e. the distance between two clusters corresponds to the maximum distance between two elements chosen from the two clusters). The dendrogram was cut at the position corresponding to a reciprocal overlap of 0.7 to create the partition, thus ensuring that all pairs of deletion calls within the same cluster had a reciprocal overlap of at least 0.7. To boost the number of real deletions retained, the partitioning algorithm was applied to the Confident deletions alone, and Possible deletions were subsequently assigned to the same groups as the Confident deletions with which they shared the highest reciprocal overlap. Applying this procedure resulted in 201,986 distinct latent deletions.

Some sets of overlapping Manta deletion calls, which were probably generated by the same underlying deletion, had very different variances of left-hand and right-hand breakpoints across samples. This caused the partitioning procedure to fail to assign such calls to the same latent deletion. To address this, we employed the following remedy. First, we constructed a dendrogram using hierarchical clustering as described above on the 201,986 latent deletions. The start and end points of each latent deletion were estimated by taking the mean of the start points and the mean of the end points respectively across the Manta calls that mapped to the latent deletion, or if there were no Manta calls mapped, by taking the means across the Canvas calls. For each node in the dendrogram for which the mean reciprocal overlap of its descending latent deletions exceeded 0.5, we evaluated whether the spatial densities of start points or end points were significantly greater than expected by chance. We did this by performing a binomial test under the null that the probability of observing the start or end point of a deletion at any given base was the same in the interval as in the entire chromosome. If all *P*-values (one for each internal node in the dendrogram) were greater than a threshold, the procedure was halted, otherwise the deletions descending from the node with the lowest *P*-value were merged, and the procedure was then repeated. We selected a threshold of 0.05 which retained high specificity whilst removing a large number of highly overlapping clusters which appeared on manual inspection to represent the same latent deletions. Application of this merging step reduced the number of deletions by 24,436, resulting in 177,550 distinct deletions being retained for final analysis.

Retained deletion calls were then annotated with cohort allele frequencies based on the number of other deletions in their groups. We computed *P*-values under the null hypothesis of HWE for all common deletions as QC. The distribution of *P*-values was highly skewed (82% of deletions in unrelated Europeans with a cohort allele frequency of at least 0.05 had a $P$-value $< 0.01$) and we therefore concluded that the calling of common deletion genotypes was unreliable but retained rare deletions for downstream analysis (see below).

**Clinical reporting**

Diagnostic multi-disciplinary team (MDT) meetings reported pathogenic or likely pathogenic variants to referring clinicians for cases in all 18 domains except EDS, CNTRL, GEL, HCM, LHON and UKB. The reasons that no reports were made for these domains were as follows: the MDTs for EDS did not identify any likely pathogenic or pathogenic variants in any of the 26 EDS samples; LHON cases were all known to carry pathogenic mitochondrial variants prior to enrollment, as the scientific motivation for recruitment was to identify disk-modifying nuclear DNA variants; HCM enrolled patients who had already been found negative in a screening of HCM diagnostic-grade genes; CNTRL and UKB participants did not include rare disease patients; clinical reporting for GEL participants via the NHS Genomic Laboratory Hubs is ongoing and the results will be reported separately in due course.

*Gene lists and transcript selection*

For each of the 15 rare disease domains (i.e. all domains except CNTRL, GEL and UKB) a list of genes was created by domain-specific experts. Only genes with an established causal role (more than three independent families reported with causal variants in a gene, or two families reported plus additional functional studies and/or a mouse model) were included in these lists. The lists for the different domains overlapped (**Figure 2b**). The 2,478 gene/domain pairs, encompassing 2,073 distinct genes across all domains, were manually curated and annotated with the relevant RefSeq and/or Ensembl transcript identifiers to support variant reporting. Transcripts were selected based on, by order of priority: community input, presence in the Locus Reference Genomic (LRG) resource[62] and designation as canonical in Ensembl (**Supplementary Table 2 (Diagnostic-grade genes)**). The genes in the 15 lists were submitted to LRG, whose role is to determine the most relevant transcripts for clinical reporting. The gene lists were also loaded into the Sapientia™ web application (Congenica Inc, Cambridge, UK) for use by the MDTs reporting causal variants to referring clinicians. During the course of the project, gene lists were updated and reversioned when new evidence caused genes to meet the threshold for inclusion. A final re-analysis of all samples was performed using gene lists that were fixed at the time of the data freeze. We refer to the genes in the final lists as 'diagnostic-grade genes'.

*Variant filtering*

Variants (SNVs, indels) were shortlisted if (i) their MAF in control populations[24] was < 1/1,000 for putative novel causal variants and < 25/1,000 for variants listed as disease-causing in HGMD, (ii) their predicted impact according to the Variant Effect Predictor (VEP)[63] was "HIGH" or "MODERATE" or if the consequences with respect to the designated transcript included one of "splice_region_variant" or "non_coding_transcript_exon_variant" if the variant was in a non-coding gene, and (iii) the variant affected a gene with a known aetiological role in the patient's disease. Variants with more than three alternate alleles or an internal MAF ≥ 10% were discarded to guard against errors in repetitive regions and remove potential systematic artefacts, respectively. The above filtering criteria were applied universally to all domains, except for ICP, which adopted a higher MAF threshold of 3% for both novel and previously reported variants. The higher threshold accounted for causal variants being present in the male and non-child bearing female population. This strategy reduced the number of variants for review by the MDT from about 4 million per individual to fewer than 10, while retaining known regulatory or moderately common pathogenic variants.

*Web application for variant assessment*

For each case with prioritised variants, the variant calls, HPO-coded phenotype and the relevant metadata (unique study numbers; referring clinician and hospital; self-declared and genetically inferred sex, ancestry, relatedness to other participants, and consanguinity level) were transferred to Congenica for visualisation in the Sapientia™ web application during MDT meetings. Sapientia™ displays variant information such as predicted effect, location in the protein, MAFs in reference cohorts (e.g. ExAC, UK10K, and ESP[24,46,64], conserved regions, splice sites, and links to external resources (e.g. HGMD[49], ClinVar[65], OMIM[66] and PubMed[67], as well as showing patient data such as phenotype information in the form of HPO terms[14]. Sapientia™ also allows MDTs to annotate each variant with its likely level of pathogenicity, its contribution to the disease phenotype (complete, or partial, the latter meaning the variant was either one of two in compound heterozygosity causal of a recessive disorder or was causal of only some of the phenotypic abnormalities), and to generate tailored research reports for the referring clinicians.

*Variant interpretation in MDT meetings*

MDTs brought together experts from different hospitals across the UK and overseas, and typically consisted of an experienced clinician with domain-specific knowledge, a scientist with experience in clinical genomics, a clinical bioinformatician and a member of the reporting team. Assignment of the level of pathogenicity to variants followed the American College of Medical Genetics (ACMG) guidelines[68] and variants were marked in Sapientia™ as pathogenic, likely pathogenic or of unknown significance (VUS). Only pathogenic and likely pathogenic variants were systematically reported and VUSs were reported at the MDT's discretion. As per the REC-approved study protocol, secondary findings (e.g. pathogenic variants in *BRCA1* in patients not presenting with a relevant cancer phenotype) were not reported. Deletions involving domain-relevant diagnostic-grade genes were reviewed by the MDT and reported if deemed likely pathogenic or pathogenic. Complex rearrangements were analysed only in IRD and NDD participants[69]. The reports recommended clinical grade confirmation in an accredited laboratory before feedback of the results to the patient or modification of clinical management. Although Sanger sequencing was not performed systematically to confirm the > 1000 reports issued, no failures to confirm the variants were reported back by the referring clinicians. In addition, 177 of the variants reported in cases in the IRD domain were confirmed by Sanger sequencing (SNVs, indels or structural variants) or microarray (structural variants only)[4].

*Sensitivity of WGS to detect known pathogenic variants*

The mean depth of coverage by WGS was 36.07X across all samples. In order to assess the relative sensitivity of WGS and WES, we compared the coverage of WGS and WES at the sites of 116,449 variants in the autosomal diagnostic-grade genes catalogued as "DM" or "DM?" in

HGMD Pro 2018.1, or as likely pathogenic/pathogenic in ClinVar version 2018-07-29, excluding variants with at least one “benign” interpretation. We also assessed the sensitivity of WES at the 938 distinct SNVs and indels included in our pertinent findings reports. Only indels of length ≤ 50bp were considered. We computed coverage statistics for 1,000 representative male samples (i.e. with an XY sex chromosomal karyotype) from our WGS 150bp read data and 1,000, 1,000, 248 and 553 representative male samples respectively from each of: UK Biobank[70], INTERVAL[70] and the Columbia University exome sequencing study for chronic kidney disease (in two datasets, IDT xGen Exome Research Panel V1 samples and Roche samples)[71]. Specifically, we computed mean coverage and the 1st and 99th percentile of coverage over each set of samples. Because UK Biobank and INTERVAL sequencing reads had been aligned to human genome GRCh38, we realigned them to human genome GRCh37 with BWA mem[72] and marked duplicate reads using Picard MarkDuplicates[73]. We computed coverage using samtools after discarding secondary alignments, reads not passing quality controls, duplicates and supplementary alignments. Only bases with a base call quality ≥ 10 and reads with a mapping quality ≥ 20 were counted. The coverage of a deletion was set to the mean coverage of the deleted bases and the coverage of an insertion was set to the mean coverage of the two bases flanking the insertion breakpoint. A nominal cutoff on the mean coverage of 20X was used to distinguish between detectable and undetectable variants to diagnostic standard.

The results of our analysis showed that the number of undetectable SNVs and indels ranged from 119 to 318 and 47 to 78 respectively, across the WGS dataset and the four WES datasets. 56–72% and 42–53% of these variants, respectively, map to a small number of genes (*HBA1*, *HBA2*, *VWF* for SNVs; *HBA1*, *HBA2* for indels). 1,997–9,063 of the causal variants (1.71–7.78%), were undetected in WES datasets but only 470 (0.4%) were undetected in the WGS dataset. In keeping with this observation, 25–99 (2.67–10.55%) of the 938 SNVs and indels reported by the MDT showed insufficient mean coverage in the WES datasets and are thus liable not to have been detected by WES (**Extended Data Figure 7b**). The WES data had a much greater variance in coverage across individuals than the WGS data and, consequently, a larger proportion of individuals tended to have coverage < 20X in WES datasets than in WGS datasets (even after controlling for mean coverage): 3.60–9.75% of SNV–individual pairs and 4.07–9.31% of indel–individual pairs had insufficient coverage at causal SNVs and indels, respectively, while, by WGS, those values were only 1.63% and 1.36%, respectively (**Extended Data Figure 7c and 7d**).

*Overview of diagnostic results*

In total, 1,138 reports were generated by the MDTs, listing 1,103 distinct causal variants (731 SNVs, 264 indels, 102 large deletions, 6 others), with 266 (26.7%) of the SNVs and indels being absent from HGMD Pro2018.1 and absent from the set of ClinVar variants with a likely pathogenic/pathogenic assignation but lacking benign assignations (**Supplementary Table 2 (SNV and indel list**, **Large deletion list)**). The reported variants were observed in 303 unique diagnostic-grade genes. Halfthe reported variants were in 21 genes and a quarter in three genes (**Figure 2d**).

The smallest number of genes needed to explain at least 50% of reports ranged between one and eight across the domains (excluding NDD). For example, *BMPR2* featured in 80% of the reports issued in the PAH domain and one of *SPTA1*, *JAK2*, *NF1* and *SEC23B* featured in 50% of the reports issued in the SMD domain. The NDD domain had a more even distribution of reports over genes, with 31 genes needed to explain 51% of reports. The reports not discussed so far were distributed across between one and 91 genes, depending on the domain (**Extended Data Figure 6a**). Most of the reported autosomal SNVs and indels were absent from gnomAD and TOPMed and the lower their MAF the more likely they were to be autosomal dominant (**Extended Data Figure 6b**). Interestingly, 21 causal variants were reported for diagnostic-grade genes shared between two domains (**Extended Data Figure 6c**). For example, two BPD cases with thrombocytopenia, platelet function abnormality accompanied by bleeding and an NDD case with brachycephaly, microcephaly and global developmental delay, carried likely pathogenic missense variants in the phosphatase encoded by *PTPN11*, a gene known to harbour variants causal of Noonan syndrome[74,75]. Six MPMT cases, two SMD cases and one NDD case carried a protein-truncating variant in *NF1* causative of type 1 neurofibromatosis, manifesting variously as neoplasm of the nervous system or small intestine (MPMT), dilatation and neurofibromas (NDD), and failure to thrive, broad philtrum, xanthomatosis, cafe-au-lait spot, hypotelorism, juvenile splenomegaly, enlarged kidney, hepatomegaly, monocytosis, myelomonocytic leukemia, thrombocytopenia and anaemia (SMD) and dilatation and neurofibromas (NDD). Conversely, some patients had phenotypes caused by variants in several genes. For example, the seven patients with a causal variant in *NMNAT1* or *RPE65* all have retinal dystrophy, as expected, but two also had intellectual disability, likely caused by variant(s) in other genes.

Reports implicating likely pathogenic or pathogenic variants were issued for 1,138 affected individuals. The domain specific diagnostic yield (the percentage of diagnosed patients) ranged from 0% (for PMG) to 53.9% (for IRD) (**Figure 2c**). The wide range in yield can be attributed to variation in the extent of genetic screening before enrolment (see Enrolment section), variation in the genetic architecture of disease and variation in the depth of public domain knowledge of the genetic aetiologies of diseases. For example, the high yield for the IRD domain, in which more than half of the affected individuals received a conclusive report, can be attributed to detailed retinal image phenotyping, a lack of genetic pre-screening and a predominantly autosomal recessive mode of inheritance. In SRNS, extensive genetic pre-screening by WES[76] and the apparent contribution of a polygenic component to SRNS risk[77] led to a relatively low diagnostic yield (11.4%). In BPD, extensive phenotypic pre-screening was performed (e.g. full haemostasis testing). However, certain phenotypic traits, such as bleeding, are strongly influenced by environmental exposures (e.g. trauma, including surgery), and the genetic architecture of BPDs is diverse, resulting in an overall yield of 12.7%. The reason that no causal variants were identified in PMG patients remains unclear, but there is emerging evidence that most cases of PMG can be explained by polygenicity. (Levine *et al*, manuscript under review).

Deletion calling by WES remains challenging, particularly the calling of short deletions encompassing only one exon or part of an exon and the calling of deletions of exons with poor coverage[78]. WGS allows more reliable calling of large deletions. 177,550 distinct deletions were retained for analysis after filtering. We excluded deletions with an allele frequency greater than 1 in 4,000 in the cohort of 10,259 unrelated individuals and deletions that did not overlap any diagnostic-grade genes for a case's domain. We visually reviewed the aligned read data of the 524 remaining unique deletions using IGV. Nearly 85% (444) of these deletions were deemed confirmed or likely to be real after this visual review and were presented to the relevant MDTs. Of these, 23.42% (104) were labelled as likely pathogenic or pathogenic (**Supplementary Table 2 (Large deletion list)**). The length of the 104 distinct reported deletions ranged between 203bp and 16.80Mb (mean 786.33Kb; median 15.91Kb) and nearly half (43) were absent from ClinVar.

A total of 28 heterozygous deletions encompassing autosomal recessive genes were returned to the referring clinician, of which 25 were reported in compound compound heterozygosity with an SNV or an indel on the other haplotype. For example, one SRNS case carried a rare 14.55Kb deletion which removed the last 7 exons of *NPHS1* in compound heterozygosity with a splice donor variant. These two variants explain the patient's phenotype. This is the first report of a large deletion in *NPHS1* causing SNRS.

15 of the reported deletions were called as homozygotes. Of these, nine were carried by probands born to consanguineous parents. In another case the homozygosity was illusory due to an overlap between two distinct heterozygous deletions originating from the unrelated parents. In the remaining five cases, the rare homozygous deletions were confirmed by visual inspection in IGV and by additional genotyping, and uniparental disomy was excluded as a possible cause. Overall, 103 of the 1,038 reports issued (9.9%) included a large deletion, confirming the clinical importance of this category of variants and the value of using accurate deletion calling algorithms. In addition, for the IRD and NDD domains, we performed an additional analysis for complex structural variants (cxSV) and identified two cases harbouring a cxSV in a pertinent diagnostic-grade gene[69].

The penetrances of causal genetic variants differed by domain. In general, causal variants in diagnostic-grade genes for IRD, NDD, BPD and SMD have high penetrance (conditional on a second high penetrance causal allele being present on the other haplotype in the case of autosomal recessive diseases). In contrast, causal rare variants in *BMPR2*, which are present in approximately 80% of patients with familial PAH and in 20% of patients with idiopathic PAH, have been reported to have a penetrance as low as 14% in males[8,79]. It is thought that environmental triggers are important in precipitating the disorder. In NPD, penetrance is also modulated by environmental exposures, such as low temperature, which is a causal risk factor for non-freezing cold injury.

*Upload of reported variants to public repositories*

The reported alleles and their clinical interpretations have been deposited in ClinVar and are available under the study names "NIHR_Bioresource_Rare_Diseases_13k",

“NIHR_Bioresource_Rare_Diseases_MYH9”, “NIHR_Bioresource_Rare_Diseases_PID” and “NIHR_Bioresource_Rare_Diseases_Retinal_Dystrophy”, and also in DECIPHER.

*Reported variants informing changes in clinical management*

We did not systematically collect information about whether the MDT reports led to changes in clinical management or the repurposing of existing drugs. There have, however, been striking examples of how the latter has changed the care path of patients enrolled in the project. Firstly, 27 patients with early-onset dystonia were shown to carry causal variants in the histone methyltransferase gene *KMT2B*, and many were treated successfully by deep brain stimulation[80]. Secondly, gain-of function variants in *DIAPH1* cause macrothrombocytopenia and deafness[81]. We have recently shown that the treatment of these patients with Eltrombopag, an FDA-approved drug for the treatment of autoimmune thrombocytopenia, increases the platelet count to a safe level in the perioperative setting[82], thereby reducing the need to use donor platelet concentrates. Finally, we identified a pedigree with a p.E527K gain-of-function variant in the kinase SRC resulting in juvenile myelofibrosis and severe thrombocytopenia, further complicated by osteoporosis. After discovering that this variant was pathogenic, the proband (case 27 of the published pedigree) was successfully cured of her thrombocytopenia by an allogeneic haematopoietic stem cell (HSC) transplant from her HLA compatible sister, who was negative for the causal variant[83].

The identification of causal variants has improved the stratification of patient care, including the frequency of clinic visits. We showed that haploinsufficiency of *NFKB1* is the most frequent cause of primary immune deficiency. Patients with loss-of-function *NFKB1* mutations have recurrent and severe infections accompanied by autoimmunity and unexplained splenomegaly and an increased risk of oncological manifestations. These new findings have a direct impact on the care of this genetically defined category of PID patients[10]. Similarly, we identified 27 BPD cases with isolated thrombocytopenia caused by variants in *ANKRD26*, *ETV6* or *RUNX1*. These genes encode DNA-binding proteins and the pathogenic variants are associated with an increased risk for haematological cancers, which is particularly severe for patients with variants in *ETV6*[84,85] and *RUNX1*[86]. Hence, frequent follow-up clinic visits are warranted and allogeneic HSC transplants need to be considered as they can be curative. In contrast, the clinicians of the 19 patients with thrombocytopenia due to variants in *ACTN1, CYCS* or *TUBB1* could reassure their patients that their conditions were benign[87]. Such patients do not require regular follow-up but haematology consultation is required at times of haemostatic challenges (e.g. childbirth or surgical procedures, including dental ones). Finally the identification of four new genes (*ATP13A3, AQP1, GDF2, SOX17*) for PAH led to improved diagnostic sensitivity for this condition. The genetic findings have also confirmed that mutations in *BMPR2*[88] and *EIF2AK4* are associated with a poorer prognosis. In particular, patients carrying causal variants in *EIF2AK4* should be referred early as potential lung transplant recipients[7].

**Structural variants in patients with null protein phenotypes**

We recorded comprehensive intermediate phenotype data on most BPD patients. This included measurements relating to haemostasis, including of coagulation factor levels (Factor I (fibrinogen; *FGA, FGB, FGG*), II (*F2*), VII (*F7*), IX (*F9*), XI (*F11*), von Willebrand Factor (*VWF*)), platelet receptor expression levels e.g. of the fibrinogen receptor (GPIIbIIIa; *ITGA2B*/*ITGB3*) and red cell Rh grouping information. These data allowed us to identify 15 unexplained cases who had a complete or near absence of expression of one of these proteins. We refer to such an absence of protein expression as a 'null phenotype.' Genetic defects relating to null phenotypes tend to have recessive inheritance. We identified:

- eight unsolved cases with null phenotypes for coagulation factors;
- six unsolved cases of possible Glanzmann's thrombasthenia (diagnosed from observing impaired platelet aggregation in response to at least two of the five agonists arachidonic acid, collagen, epinephrine, thromboxane analogue and TRAP, and observing normal agglutination in response to ristocetin);
- a patient with a mild bleeding disorder accompanied by an unexplained haemolytic anaemia, who had no Rh group proteins on her red cell membranes.

Three of the 15 cases with null phenotypes had a rare coding variant on one haplotype of the relevant gene while the other haplotype appeared to be wild-type according to the standard variant calls. Through visual inspection of the reads in IGV, we were able to resolve two of these three cases.

- The first case had Glanzmann's thrombasthenia and carried a variant in *ITGB3* which encodes a premature stop at amino acid position 242. Visual inspection of the gene body revealed an excess of improperly mapped reads in intron 9, primarily due to alignment of paired-end reads facing away from, rather than towards, each other (**Extended Data Figure 8a**). We counted the number of improperly mapped read pairs in the region in our collection after excluding the GEL domain participants (as we do not hold their phenotypic data), and found this case to have the greatest number of improperly mapped reads out of 6,656 individuals whose samples were sequenced using 150bp chemistry (**Extended Data Figure 8b**). In a separate dataset of 304 WGS samples sequenced with 150bp chemistry, we identified two further individuals (a Glanzmann's thrombasthenia case and his mother) with a similar excess in intron 9, primarily due to read mates aligning to different chromosomes (**Extended Data Figure 8c,d**). The proband's platelets were devoid of GPIIbIIIa and the mother had a ~50% reduction in expression of GPIIbIIIa (data not shown). The proband carried a rare paternally inherited missense variant encoding a change from threonine to proline at amino acid position 456 of *ITGB3*. Sanger sequencing of the *ITGB3* cDNA obtained by reverse transcription of the RNA from the proband's platelets showed exclusive expression of the proline coding allele and provided no evidence of an alternatively spliced transcript (data not shown). In order to resolve the genetic sequence of intron 9, we performed Oxford Nanopore-based sequencing of long-range PCR-amplified target DNA as previously described[69]. The flow cell ran for 3 hours, and the mean coverage was 863,986X**.** We identified an insertion of an SVA (Alu, SINE-VNTR-Alu) retrotransposon element of 2,270bp at position 45369041 of chromosome 17, predicted to induce nonsense-mediated decay. SVA elements are present throughout the reference genome, explaining the short reads' mates aligning to various chromosomes

(**Extended Data Figure 8e**). To date, roughly 130 pathogenic variants caused by retrotransposon elements have been documented in the literature[89], but none of them have been implicated in Glanzmann's thrombasthenia. Here we report the first such example and demonstrate the utility of long-range sequencing reads for identifying complex causal variants suggested by short-read WGS analysis.

- The second case had red cells which serotyped negative for all common antigens of the Rh blood group system. Upon further investigation, the red cells were shown to lack both the RHD and RHCE proteins and also the Rh-associated glycoprotein RHAG. This very unusual $Rh_{null}$ red cell phenotype is known to cause a syndrome characterised by chronic haemolytic anaemia of varying severity[90]. The most frequent genetic explanation for this rare condition is the presence of loss-of-function variants on both *RHAG* haplotypes, because the RHAG protein is essential for the expression of the RHD and RHCE proteins on the red cell membrane[91]. This case was found to carry a known causal loss-of function splice donor acceptor variant in intron 2 of *RHAG*[92]. The second mutation was identified by visual inspection of the reads of the *RHAG* locus, as a heterozygous tandem duplication spanning exons 2–7.

**Genetic association between rare variants and rare diseases**

*Statistical approach*

We used the BeviMed statistical method[93] to identify genetic associations with rare diseases in our dataset. Each run of BeviMed requires selection of a subset of unrelated study participants partitioned into cases and controls and a set of rare variants to test for association. To achieve adequate power, the cases should be chosen to maximise the chance that they share a common genetic aetiology (e.g. by identifying a group of individuals with similar phenotypes) and the rare variants should be chosen to maximise the chance that they are aetiologically exchangeable (e.g. by identifying variants predicted to have a similar effect on a gene product). BeviMed computes posterior probabilities over the hypotheses:

- no association between any variant and case/control status,
- dominant association between a pathogenic subset of variants and case/control status,
- recessive association between a pathogenic subset of variants and case/control status.

BeviMed also reports the posterior probability that each variant is pathogenic conditional on the mode of inheritance. We impose a prior correlation structure on the probabilities that variants are pathogenic which reflects prior belief about which class of variant is disease causing. This correlation structure derives from groupings of variants by their VEP consequences. The class of variant responsible for causing disease is inferred by BeviMed, which can suggest a molecular aetiology.

*Case/control groupings using phenotypic tags*

A set of phenotypic 'tags' were defined for each domain that determined the case/control groupings for BeviMed. Cases shared a particular tag if their phenotypic characteristics were considered compatible with a similar genetic aetiology. Some tags were set using logical rules applied to the HPO terms and other data and others were set manually. The full set of tags and corresponding numbers of cases and controls can be found in **Supplementary Table 3**

**(Phenotypic Tags)**. Given a particular tag and a set of rare variants, the corresponding case/control groupings were obtained as follows:

1. cases: unrelated (i.e. at most one per family) affected individuals with the tag who have not been diagnosed by an MDT with a variant outside the set;
2. controls: members of the maximal set of unrelated individuals who do not have the tag and who are not related to any of the cases.

*Selection of variants affecting transcript sequences*

For each gene, we selected SNVs/indels for inclusion in the BeviMed analyses. Only variants with a within sample MAF in unrelated individuals less than 0.002 were used. We defined $PMAF_X$ for each variant as the probability that $Y \geq C$, where $Y$ is a random minor allele count for a variant with MAF 1/X and C is the fixed observed minor allele count. Variants had to have a $PMAF_{500}$ greater than 0.05 in several subsets of participants: European probands, non-European probands, European members of the maximal unrelated set and non-European members of the maximal unrelated set. SNVs/indels were retrieved from the HBase variant database and were filtered as follows:

- minimum OPR ≥ 0.98 for each of the three batches;
- unless a variant was in HGMD with class "DM" or "DM?", it had to have $PMAF_Z$ greater than 0.05 in every gnomAD population (only gnomAD males were used to compute the $PMAF_z$ on variants in the non-PAR of X). We used Z = 1,000 for recessive association analyses and Z = 10,000 for dominant association analyses;
- SNVs with a CADD phred score < 10 were excluded;
- variants in the non-PAR of X that appear only as heterozygotes in males were excluded;
- multi-allelic variants for which the reference allele was the minor allele were excluded;
- unless a variant was in HGMD with class "DM" or "DM?", it had to have a CellBase-predicted consequence of type transcript_ablation, splice_acceptor_variant, splice_donor_variant, stop_gained, frameshift_variant, stop_lost, start_lost, transcript_amplification, inframe_insertion, inframe_deletion, missense_variant, protein_altering_variant, regulatory_region_ablation, non_coding_transcript_exon_variant or 5_prime_UTR_variant on a transcript of type lincRNA, miRNA, misc_RNA, Mt_rRNA, Mt_tRNA, protein_coding, rRNA, snoRNA, snRNA, TR_C_gene, TR_D_gene, TR_J_gene, TR_V_gene, IG_C_gene, IG_D_gene, IG_J_gene or IG_V_gene ("retained biotypes").

Large deletions were subjected to the same $PMAF_{500}$ filters as the SNVs and indels.

In order to identify groups of variants that had high a prior probability of being aetiologically exchangeable, we grouped variants into gene specific (non-mutually exclusive) classes labelled "5' UTR", "moderate" and "high-impact":

- "5' UTR:" there were no VEP consequences with a HIGH or MODERATE impact and at least one transcript was annotated with the VEP '5_prime_UTR_variant' consequence;

- “moderate”: at least one transcript was annotated by VEP with a MODERATE or HIGH impact consequence or with the VEP ‘non_coding_transcript_exon_variant’ consequence
- “high-impact”: at least one transcript was annotated by VEP with a HIGH impact consequence or the variant was a large deletion that overlapped an exon of a transcript with a retained biotype.

Thus, all variants in the high-impact class were also in the moderate class, while variants in the 5’ UTR class were not present in any other classes. The rationale for combining missense variants and variants with a HIGH impact consequence label is that both types of variant are capable of inducing loss of function. The conditional prior probabilities of the models that treated each of these three groupings of variants as disease causing were 0.01, 0.495, and 0.495 for the “5’ UTR”, “moderate” and “high-impact” variant classes, respectively. All priors in the BeviMed model were set at the default values values, including the prior probability of association accumulated across all association models, which was set to 0.01.

*Post-processing of BeviMed results*

We applied three post-processing steps to the BeviMed results.

1. Allele frequency datasets are much smaller for non-European ancestry populations than for the European ancestry population. Consequently, the false positive rates of BeviMed association analyses relying on non-European cases can be inflated if variants which are moderately rare in non-Europeans are unintentionally included. To guard against this, when we obtained a BeviMed posterior probability of association (PPA) > 0.75 for a gene–tag pair, we re-ran BeviMed excluding all variants:
    - present in heterozygosity in at least two individuals from the same non-European ancestry (e.g. South Asian or African) group (homozygosity is likely to indicate consanguinity) and
    - absent from all Europeans.
    - If the new BeviMed PPA was < 0.25, we flagged the result as being dependent on variants that may be moderately rare in a non-European population. This should be taken into account in follow-up functional studies.
2. Genomic regions spanning more than one gene that are deleted in multiple cases can potentially induce multiple correlated genetic associations with a particular disease tag. In order to guard against false positive associations due to these correlations, we post-processed the results to ensure that no two genes having a PPA with a given tag > 0.75 both depended on the same set of deletions to obtain a PPA > 0.25. This was done as follows. For each tag, if at least two genes had a PPA > 0.75, we looped through each pair of such genes, and marked the gene with the lower PPA for removal if, when deletions shared between the genes were excluded, its PPA dropped below 0.25. After completing the loop, the associations corresponding to marked genes were removed.
3. The results with PPA > 0.75 may contain multiple associations between a gene and correlated phenotypic tags. In order to remove such redundant information, we retained only one association per gene, corresponding to the tag for which the highest PPA was obtained.

*BeviMed results for genes*

BeviMed was run for each tag/gene pair. For each pair, the total PPA, summing over all modes of inheritance and classes of variant (5' UTR, moderate and high-impact) was recorded. A total PPA > 0.75 was considered good evidence of a causal relationship. The 95 genetic associations that remained after post-processing are shown in **Figure 3** and in **Supplementary Table 3 (BeviMed Association Tags)**.

*Observed false positive rate*

Let *TPR* and *FPR* denote the observed true positive rate and the observed false positive rate, respectively. Note that for arbitrary *C* > 0, *D* > 0, if $\frac{A}{C} < \frac{B}{D}$ , then $\frac{A}{C} < \frac{A+B}{C+D}$ . Consider the *n* tests that are not true positives. Let us suppose that *np* of these tests correspond to gene–tag pairs for which the gene truly mediates the aetiology of the tag and that *n*(1 − *p*) correspond to gene-tag pairs for which this is not the case (i.e. to the null hypothesis). Put

$$\begin{aligned} A &= 16 - np \cdot TPR \\ B &= np \cdot TPR \\ C &= n(1-p) \\ D &= np \end{aligned}$$

The statement $\frac{A}{C} < \frac{B}{D}$ is equivalent to 16/*n* < *TPR*. This holds if we assume *FPR* < *TPR*, for then $FPR = \frac{16 - np \cdot TPR}{n(1-p)} < TPR$, which implies that $16 - np \cdot TPR < TPR \cdot n(1-p)$ and thus $16 < TPR \cdot n$.

Consequently,

$$FPR = \frac{16 - np \cdot TPR}{n(1-p)} = \frac{A}{C} < \frac{A+B}{C+D} = \frac{16}{n}.$$

In total, we have tested 23,090 genes for association with 29 tags (i.e. we performed 669,610 statistical tests), of which 102 are confirmed true positives. (Note that only 79 of the 102 confirmed associations are reported in Figure 3 because for each gene we presented just the tag with the strongest posterior evidence for association.) Thus, we can set *n* = 669,610 - 102 and infer an upper bound on the observed false positive rate of 16/*n* = $2.4 \times 10^{-5}$.

*Positive predictive value*

We performed a comprehensive analysis of the study-wide positive predictive value (equivalent to 1 minus the false discovery rate). We present three alternative approaches to making inferences about the study-wide positive predictive value (PPV).

<u>Lower bound on observed PPV.</u> Only 16 of the 95 genetic associations we report are unconfirmed, which sets a lower limit on the observed PPV of 83%.

<u>Posterior PPV.</u> The mean posterior probability for the reported associations gives a posterior PPV of 93%.

Permutation PPV. We estimated the study-wide expected number of gene–tag associations with a PPA > 0.75 under the assumption that all tests are null. This estimate, divided by the observed number of gene–tag associations with a PPA > 0.75, gives an estimate of the FDR (equivalently, 1-PPV) which is conservative because in practice a proportion of tests are not null. Our approach follows closely that proposed by Tusher, Tibshirani and Chu[94].

In order to estimate the expected number of gene–tag associations with a PPA > 0.75 under the null, we generated 100 permuted datasets and ran the entire association analysis procedure on each dataset.

We used the genotype data from the maximal unrelated set of participants and outcome data from participants selected within each ancestry group according to the following procedure:

1. Let $k_g$ be the number of participants in the maximal set of unrelated individuals who belong to ancestry group $g$.
2. Create a participant (rows) by tag (columns) matrix $M_g$ from the real outcome dataset for participants in ancestry group $g$, in which elements are coded:
    - 1 to indicate that the participant is a case for the tag,
    - 0 to indicate the participant is a control for the tag,
    - NA to indicate that the participant is to be excluded from analyses for that tag due to relatedness.
3. Row sort the matrix on the row count of 1s, breaking ties by sorting on the row count of 0s.
4. Select the leading $k_g$ rows as outcome data.

On each iteration of the permutation procedure, for each ancestry group $g$, we randomly assigned the $k_g$ rows of the outcome matrix $M_g$ to the genotype data of the $k_g$ unrelated participants. We applied BeviMed to the ancestry unified dataset of $\sum_g k_g$ rows. For each gene–tag combination, we relabelled cases as controls in the outcome data if they were so relabelled in the actual analysis (because they had been genetically explained by variants in a different gene). We applied the study-wide post-processing filters described above.

The entire process was repeated 100 times. This process, without permutation, generated 95 results with a PPA > 0.75. Over 100 repetitions, the mean number of PPAs > 0.75 was 20.0, giving a permutation PPV of 79%. Increasing the PPA threshold from 0.75 to 1.0 increased the PPV approximately linearly from 79% to 100% (not shown).

*Estimated false positive rate*

Following the procedure described above to compute the permutation PPV, we estimated the false positive rate as the mean proportion of PPAs > 0.75 before retaining only the top associated tag for each gene. In total, we tested 23,090 genes over 29 tags (i.e. we performed

669,610 statistical tests), of which on average 20.0 had PPA > 0.75, giving an estimated FPR of $3.0 \times 10^{-5}$.

**Polygenic and rare variant associations with phenotypic extremes in the UK Biobank cohort**

*Selection of participants*

A genomewide association study (GWAS) of FBC traits measured in approximately 173,000 European ancestry participants in the UK Biobank cohort (*n*~133,000) and the INTERVAL study (*n*~40,000)[70] has previously identified 582 genetic variants independently associated with quantitative properties of mature red blood cells[95]. Most of these variants were common: only 40 had an in sample MAF lower than 1%. The GWAS design had limited power to identify associations with rare variants, due to the imprecision of genotype imputation at low allele frequencies (**Appendix 2: Extreme red cell traits in UK Biobank**). In order to identify rare variants associated with full blood count measured properties of red cells, we attempted to derive a univariate quantitative phenotype with high rare-variant heritability. To do this, we used 65 previously reported GWAS associations between variants with MAF < 1% and phenotypes of red cells as a model for the effect of rare variants on mature red cell phenotypes (**Figure 4a**). 384 individuals were selected for WGS from each tail of the quantitative phenotype, of which 383 from the left tail and 381 from the right tail were successfully sequenced and passed QC (**Figure 4b**). The left tail of the score (BeviMed tag "Left") included individuals with lower than average RBC# and higher than average MCV, while the right tail (BeviMed tag "Right") included individuals with higher than average RBC# and lower than average MCV (**Figure 4c**). A more detailed explanation of the technicalities of the selection procedure can be found in **Appendix 2: Extreme red cell traits in UK Biobank.**

*Polygenic contribution to tail status*

We first sought to understand the polygenic variant contribution to the risk of being extreme for the quantitative phenotype, by using GWAS variants with published associations with RBC# or MCV to construct a polygenic predictor of the quantitative phenotype. 448 of the 468 variants reported to be associated with RBC# or MCV in[95] had well called genotypes (overall pass rate > 0.8) in the WGS data. We identified well called high linkage disequilibrium proxy variants for 16 of the remaining 20 variants. 6 of these 16 variants had $r^2 = 1.0$ with their proxy variant in the European component of the 1000 Genomes data, 9 had $r^2 > 0.95$,14 had $r^2 > 0.9$ and all had $r^2 > 0.7$. The four variants for which no proxy could be found jointly explain 0.04% of the variance of the phenotype in 116,689 European ancestry participants included in the interim release (May 2015) of the UK Biobank imputed genotype data. We regressed the quantitative selection phenotype jointly on the 464 well called/proxy variants using 298,607 European participants from the full release of the UK Biobank imputation data (July 2017) who were not included in the interim release and were not WGSed for this study. The polygenic score formed by the linear predictor of this regression explained 22% of the variance in the phenotype in sample ($R^2 = 0.219$).

We identified 508 unrelated European participants in study domains other than UKB who can be considered a population sample with respect to their polygenic score values. These are rare disease patients whose pathology has been fully explained by rare variants declared as "likely pathogenic" or "pathogenic" in an MDT, or healthy individuals with a patient relative, the phenotype of whom has been fully explained in the same way. We call this group 'unselected' because they have not been selected with respect to their polygenic red cell genotype. The distribution of the polygenic score in the two UK Biobank selected tails and in the unselected category is shown in the box plots in **Figure 4d**. There is strong statistical evidence for an increasing trend in the mean value of the score with an ordinal variable for category membership (0 = left tail,1 = unselected controls, 2 = right tail; *P*-value = $2.4 \times 10^{-96}$, linear regression).

The central limit theorem implies that the distribution of the polygenic score should be approximately Gaussian in the unselected control group. Under the assumptions 1) that the quantitative selection phenotype (**Figure 4b**) has a Gaussian distribution and 2) that the mean of the phenotype depends linearly on the polygenic score, the expected distribution of the score in each tail group can be derived, conditional on the quantitative trait tail selection thresholds and conditional on the population proportion of phenotypic variance explained by the polygenic score (see **Appendix 2: Extreme red cell traits in UK Biobank** for details). The probability densities of these distributions are displayed as violin plots underlying the box plots in **Figure 4d**. The selection thresholds chosen correspond to the quantitative phenotype value for the least extreme individual selected from the corresponding tail. (This is a conservative approach, because the selection procedure involved some recalibration of the phenotype for age and sex in the tails of the score). Comparison of the expected distributions with the observed data suggests that tail status is less well predicted by the polygenic score than the Gaussian variance components modelling predicts. A number of factors could explain the incongruence between theory and observation. The tails of the quantitative phenotype are heavier than Gaussian and the excess density in the tails could be due to rare events unrelated to polygenicity: for example, haematology analyser measurement error. The excess may also be partly explained by the effects of rare or extremely rare alleles. This motivates analyses designed to search for such alleles and the genes mediating their associations.

*Whole genome sequencing GWAS*

We performed a WGS GWAS with the aim of identifying genetic variants (particularly moderately rare genetic variants) associated with the ordinal phenotype indicating group membership. We analysed the 11,147,508 genetic variants with an overall pass rate > 0.8 and an in-sample MAF > 0.25% in the 1,272 individuals in the two tails and the unselected control category. For each variant, we regressed the ordinal outcome on the alternative allele count using adjacent categories logistic regression, under the assumption that the log-odds for the right tail *vs* the unselected controls is the same as the log-odds for the unselected controls *vs* the left tail. Our analysis identified no novel associations at a significance threshold of $1 \times 10^{-7}$. However we did replicate two well known common variant red cell associations in the MYB-HBS1L (*P*-value = $1.4 \times 10^{-10}$) and ODF3B-TYMP-SCO2 (*P*-value = $1.4 \times 10^{-8}$) loci[96].

*BeviMed analysis*
The participants from each tail who passed QC were treated as a rare disease case group in a BeviMed analysis. We also performed a combined analysis in which the participants in both tails were merged into a single case group.

The inferred posterior probabilities for the genes with the strongest evidence for association (posterior probability greater than 0.4) for the two tail group analyses are displayed in **Figure 4e**. The strength of evidence (respective posterior probabilities 0.997 and 0.575) that rare variants in *HBB* and *TFRC* are associated with the low MCV/high RBC# phenotype (the right tail of the phenotype) is entirely consistent with biological knowledge (**Supplementary Table 3 (BeviMed Association UK Biobank)**) and these associations can be considered 'positive controls'. Similarly, prior biological knowledge suggests that the association identified with the transcription factor *CUX1* is very plausible while the associations identified with *ALG1*, *ZNF407* are plausible. The weight of evidence from a detailed examination of the alleles implicated in the other genes exhibiting a BeviMed PPA > 0.4, together with the expression profiles of the genes in blood cells and other tissue types is less compelling. A list of all variants in the associated genes, for which the posterior probability of pathogenicity is greater than 0.5, is given in **Supplementary Table 3 (BeviMed Variants UK Biobank).** In conclusion, our analysis demonstrates that genomic loci carrying rare alleles causing large deviations in quantitative traits can be identified by applying the BeviMed analysis approach. Since the extreme tails of any heritable quantitative trait are likely to be under negative selection, mutations in such genes may cause rare diseases or be otherwise clinically relevant. The forthcoming WGS of the full UK Biobank cohort offers a further opportunity to search for rare variant associations with other biomedically relevant quantitative intermediate traits, including blood cell traits not explored here.

**Matchmaker Exchange**
We used GeneMatcher[97], which is part of Matchmaker Exchange[98], to identify additional patients, external to the cohort, carrying rare variants in candidate disease genes identified through a manual review process informed by the literature. We were able to find matches for three patients:

1. A BPD patient with a complete lack of serotonin storage in his platelet dense granules had symptoms including hypotonia, mental disability, epilepsy, uncontrolled movements and gastrointestinal problems. The patient had consanguineous parents and was homozygous for a rare allele in *SLC18A2,* which codes for VMAT2, a serotonin transporter[99]. Consequently, the patient was deficient for VMAT2. We identified a matching child with a similar clinical phenotype using GeneMatcher. We subsequently became aware of a case report of a patient with VMAT2 deficiency exhibiting a similar neurological phenotype[100].
2. A BPD patient and his sister with a platelet function defect and mild thrombocytopenia had symptoms including psychomotor retardation and epilepsy. Both siblings inherited a rare allele in *MADD* from each of their parents. The siblings matched with 13 patients

from 10 different families with similar neurological phenotypes using GeneMatcher (manuscript in preparation). However, platelet studies in these other patients have not yet been performed.

3. An NDD patient with intellectual disability, autistic features and seizures was matched to three unrelated patients with a similar phenotype using GeneMatcher. In addition, a fourth patient was matched by personal correspondence[101]. In all five patients, *WASF1* had been independently identified as a strong candidate because it is constrained against loss-of-function variants in ExAC (probability of loss-of-function intolerance (pLI) = 0.91) and is highly and specifically expressed in the adult human brain[102]. All variants were *de novo*, predicted to induce a loss of function, and absent from 1000 Genomes, ExAC and gnomAD. Interestingly, they clustered around the WASP-homology 2 (WH2) domain, in the highly conserved C-terminal actin-binding WCA region.

**Regulome analysis**

*Ethics*

Blood samples for generating open chromatin and histone modification data for activated CD4+ (aCD4) T-cells were collected with written and signed informed consent, approved by the East of England – Cambridgeshire and Hertfordshire REC (reference 05/Q0106/20). The other epigenetic data described in this section have been released as part of previous studies. Information about the consent under which these other data were generated can be found in the publications cited below.

*Definition of cell type specific regulomes*

Given that WGS provides variant calls across the entire genome, we sought to identify rare variants that exert their effect on phenotype through disruption of non-coding (possibly non-exonic) regulatory elements, such as enhancers. Genetic variation in these elements has not been systematically assayed in previous genetic studies of rare disease, which have largely relied on WES or other targeted sequencing. Regulatory elements are largely cell type specific. We sought to identify regions occupied by regulatory elements in cell types relevant to the biology of a subset of the rare disease domains (**Supplementary Table 3 (Regulome Cell Data)**). We call the set of regulatory elements corresponding to a particular cell type a 'regulome'. We determined the regulomes for aCD4 cells, B cells (B), erythroblasts (EB), megakaryocytes (MK), monocytes (MON) and resting CD4+ T cells (rCD4).

For each cell type, we applied the RedPop method (see below) to identify regulatory elements from open chromatin data (ATAC-seq) and histone modification data (H3K27ac). Additionally, we had access to the following transcription factor (TF) chromatin immunoprecipitation sequencing (ChIP-seq) data, which we used to call binding sites (see below) to expand the regulomes: FLI1, GATA1, GATA2, MEIS1, RUNX1, TAL1 and CTCF in MK; GATA1, KLF1, NFE2 and TAL1 in EB; and CTCF in MON and B.

For each cell type, the regulome build process proceeded as follows:

1. call regulatory elements from ATAC-seq/DNAse-seq and H3K27ac-seq data using RedPop,
2. call TF binding peaks using ChIP-seq data if available and obtain enrichment scores,
3. discard TF regions with an enrichment score < 10 unless they overlap between at least two different TFs,
4. collapse overlapping features to obtain a single genomic track,
5. merge features within 100bp of each other.

Each regulome feature was assigned a *target gene* label using either gene annotations from Ensembl 75 or a compendium of previously published promoter capture Hi-C (pcHi-C) data[103] as follows:

1. assign a feature to a gene if the feature overlaps the gene or the region up to 10,000bp either side of the gene body,
2. assign a feature to a gene if the feature overlaps the gene's pcHi-C "blind spot" (the region spanning the *Hind*III restriction fragment overlapping the gene's transcription start site and the two adjacent *Hind*III restriction fragments),
3. assign a feature to a gene if the feature overlaps a linked promoter interacting region identified using pcHi-C in the same cell type.

**Regulatory element detection using patterns of peaks (RedPop)**

We sought to identify regulatory elements in cell types of relevance to the rare disease domains in this project, in order to search for possible pathogenic variants within them. It is important to locate these elements because they are much more likely to harbour pathogenic variants than other non-coding regions of the genome. Open chromatin data alone do not provide sufficiently high spatial resolution and transcription factor ChIP-seq restricts to regions bound by proteins targeted by specific antibodies. Open chromatin and histone modification (H3K27ac) sequencing data together, on the other hand, can be used to detect regulatory elements with high resolution and in an unbiased fashion. Open chromatin around binding sites typically results in a broad, low-resolution peak of elevated ATAC-seq/DNAse-seq coverage. The surrounding nucleosomes of a regulatory element are typically acetylated, leaving two peaks in H3K27ac coverage, spaced a few hundred bp apart. By combining the genomic coverage tracks of an open chromatin and an H3K27ac assay, regulatory elements can be detected with high precision. We developed an algorithm for regulatory element detection using patterns of peaks (RedPop) (manuscript in preparation) that utilises these patterns.

First, ATAC-seq/DNAse-seq and H3K27ac ChIP-seq reads were aligned to the genome using the BWA[72] aln command. Open chromatin regions were then called using F-Seq[104]. The called open chromatin regions were extended upstream and downstream symmetrically until their lengths were at least 3.2Kb. Any adjacent overlapping segments were merged. Each merged segment was then considered individually. For each segment, the following sets of vectors were generated.

1. *Globally* normalised coverage tracks of open chromatin and H3K27ac. These tracks are position specific read count vectors, which are normalised by the mean coverage genome-wide and smoothed by averaging consecutive segments of 40bp.
2. *Locally* normalised coverage tracks of open chromatin and H3K27ac. These tracks are computed as the globally normalised coverage divided by the mean coverage in the surrounding 800bp region.
3. The covariance track. This track contains the covariance between the open chromatin and the H3K27ac globally normalised coverage tracks, computed in 800bp sliding windows and smoothed by averaging consecutive segments of 800bp.

Any positions for which the value of the smoothed covariance was < -1 and less than the values in the surrounding 160bp were declared local minima. At each position declared a local minimum, the nearest contiguous segment for which the locally normalised open chromatin coverage exceeded the locally normalised H3K27ac coverage, and and any other contiguous segments within 100bp of the local minimum, were recorded and extended symmetrically to 400bp. These segments were discarded unless the globally normalised open chromatin coverage track exceeded 47X (a default value obtained through a sensitivity/specificity study of ChIP-seq data from MKs). Any overlapping undiscarded segments were merged and recorded as the locations of regulatory elements.

*Transcription factor binding peak calling*

We applied the BLUEPRINT protocol for ChIP-seq data analysis[105]. H3K27ac histone modification sequence reads were mapped to human genome GRCh37 with BWA[72] aln method. Low-quality reads (-q 15), multi-mapped reads and duplicate reads were marked and removed with samtools and Picard[73]. We performed QC using deepTools plotFingerprint[106]. Files containing aligned read data corresponding to the same cell type were merged. The resulting file was downsampled in proportion to the number of aligned reads. H3K27ac peaks were called by MACS2[107] using the 'narrow' option.

*Identifying possibly causal deletions of elements regulating diagnostic-grade genes*

In order to identify regulatory elements disrupted in patients with a rare disease, we examined rare deletions called in cases that overlapped regulatory elements called using RedPop or using ChIP-seq analysis. We filtered these deletions to retain only those which delete an element

- with a target gene label that is diagnostic-grade for a recessive disorder encompassed by the domain of the carrier and
- which is called in a cell type relevant to the domain of the carrier (**Supplementary Table 3**).

Furthermore, the deletion, had to

- be called homozygous or hemizygous by Manta or Canvas, or
- delete an element the target gene of which contained a rare allele of 'moderate' impact (as described in the *Genetic association testing in genes* section) in the carrier.

Our approach is illustrated in **Figure 5a**. The filtering resulted in a list of only four deletions: a heterozygous deletion overlapping the 5' UTR region of the *ARPC1B* found in a PID patient who also carries a frameshift variant in the same gene (Thaventhiran *et al*, under review); a hemizygous deletion of a *GATA1* enhancer in a patient with thrombocytopenia, described below; a homozygous deletion of a CTCF binding site in the first intron of *LRBA* in a PID patient, described below; and a deletion which on manual inspection proved to have been called in error.

*Deletion of GATA1 enhancer and HDAC6 open reading frame*

The *GATA1* enhancer, called in MK and EB, was removed by a 4108bp deletion (X:48,659,245-48,663,353) together with the first four exons of the histone deacetylase 6 gene (*HDAC6*), including its ATG start codon (**Figure 5b**, **Extended Data Figure 9a**). The patient was a 9-year old boy with macrothrombocytopenia, bleeding symptoms and mild intellectual disability (ID) with an autism spectrum disorder (ASD) (**Figure 5c**). His parents are healthy, although mild asymptomatic thrombocytopenia is present in the mother (**Figure 5c**). The patient was hemizygous for the deletion while the mother was a heterozygous carrier (**Figure 5d**, **Extended Data Figure 9b**). Platelets of the proband express no HDAC6 protein(**Figure 5e**). In contrast, HDAC6 expression in platelets from the mother was comparable to expression levels in platelets from unrelated controls and from the father. HDAC6 is the major deacetylase responsible for removing the acetyl group from Lys40 of α-tubulin, which is located in polymerized microtubules[108]. Indeed, absence of HDAC6 expression in platelets was accompanied with very high expression levels of acetylated α-tubulin while non-acetylated α-tubulin expression levels were similar to levels in controls (**Figure 5e**). Platelets from the mother but not the father contain at least some platelets with hyperacetylated α-tubulin levels (**Figure 5e**). HDAC6 is extremely intolerant to LOF variants (pLI = 1) and the proband is the only patient in our cohort with a hemizygous high impact variant. **Figure 5b** shows that the deleted region also contains binding sites for several transcription factors important for megakaryopoiesis[109]. In addition, this region overlaps with a recently identified regulatory element upstream of HDAC6 that was shown to strongly control GATA1 expression in myeloid K562 cells[110]. Interestingly, GATA1 protein was significantly decreased in platelets from the proband and from the mother **(Figures 5f** and **5g**). GATA1 is an important transcription factor that regulates platelet formation, hemizygous mutations of which result in variable degrees of macrothrombocytopenia and dyserythropoiesis or anemia[111].

The proband had an elevated mean platelet volume (MPV) and platelet distribution width (PDW) and the mother had a normal MPV but an elevated PDW (**Figure 5c**). Electron microscopy (EM) analysis of platelets from the proband showed his platelets were larger and rounder than usual and had a lower than usual density of alpha granules, which is typical of the platelets of GATA1 deficient patients[112] (**Extended Data Figure 9c–e**). All the red blood cell parameters (red blood cell count, haemoglobin concentration, haematocrit, mean corpuscular haemoglobin, mean corpuscular haemoglobin concentration and red cell distribution width) of the proband were normal, although there were mild signs of dyserythropoiesis. Blood smear analysis revealed the presence of anisocytosis of red blood cells (+), moderate poikilocytosis (++), and slight hypochromasia (+). An elevated LDH level was detected (404 U/L; Normal values < 300), which has been described in GATA1 deficient patients[113]. The red blood cell parameters of the parents of the patient were normal.

HDAC6 deficiency has never been described in humans. Hdac6 knockout mice have no gross defects[114] except for altered emotional behaviour[115] and enhanced platelet spreading due to hyper-acetylated microtubules. However, their bleeding tendency has not been evaluated[116]. Structured illumination microscopy (SIM) analysis of acetylated tubulin in combination with F-actin (**Extended Data Figure 9f**) was performed on platelets of the proband under basal and activated conditions. Non-activated platelets showed disturbed marginal bands which were hyperacetylated (**Extended Data Figure 9g**). Quantification of platelet spreading on fibrinogen showed enhanced spreading for the proband while platelets from the parents were similar to those of the control (**Extended Data Figure 9h**). Hdac6 knockout mice have normal platelet counts and exhibit normal megakaryopoiesis. This is in contrast to the results of in vitro studies of human MKs depleted of HDAC6 using shRNA or the HDAC6 inhibitor Ricolinostat, which resulted in defective proplatelet formation[117]. We differentiated peripheral haematopoietic stem cells from the proband, his mother and an unrelated control and differentiated them to MKs. We observed a significant reduction in proplatelet-forming MKs amongst the MKs of the proband and only a mild reduction in the MKs of the mother (**Extended Data Figure 9i**). All MKs from the patient and some cells from the mother showed absent HDAC6 expression (**Extended Data Figure 9j**). In contrast to MKs treated with HDAC6 inhibitors, we detected obvious microtubule detects in the MKs of the patient and the mother (**Extended Data Figure 9k**).

Proplatelet formation defects have also been observed in the MKs of GATA1 deficient patients[112]. Consequently, it is difficult to distinguish the contribution of GATA1 and HDAC6 deficiency to the observed MK defect. In contrast, the platelet phenotypes observed in the proband seem to combine the GATA1 and the HDAC6 defects:

- low GATA1 expression leads to dyserythropoiesis, macrothrombocytopenia and reduced density of platelet alpha granules, and
- HDAC6 deficiency causes hyper-acetylation of microtubules, resulting in enhanced platelet spreading.

*Details on the functional analysis of the GATA1 enhancer/HDAC6 deletion*

<u>PCR and Sanger sequencing to validate the HDAC6 deletion</u>. Genomic DNA was extracted from peripheral leukocytes. PCR was performed with primers flanking the deletion (HDAC6-F: 5'-catcttcaagaggatcagagg and HDAC6-R: 5'- catagctagacactggtt), generating a PCR fragment of 358bp when the deletion is present. Sanger sequencing of PCR fragments was performed using the same primer sets.

<u>Antibodies</u>. The following antibodies were used: rabbit HDAC6 (clone D2E5, Cell Signaling technology, Danvers, MA, USA), mouse anti-acetylated tubulin antibody (clone 6-11B-1, Sigma, St Louis, MO, USA), mouse anti-alpha-tubulin (A11126, Thermo Fisher Scientific, Waltham, MA,

USA), rabbit VWF (Dako, Aligent Technologies, Leuven, BE), mouse CD63 and rat GATA1 N6 (Santa Cruz Biotechnology, Dallas, TX, USA), rabbit GATA1 (NF that was produced against recombinant N-terminal zinc finger[118], rabbit GAPDH (14C10, Cell Signaling) and integrin beta3 (sc- 14009; Santa Cruz Biotechnology).

Electron and fluorescent microscopy of platelets and immunoblot analysis. Electron microscopy for platelets from the proband was performed as described[111]. Immunostaining of resting and fibrinogen spread platelets was performed for platelets from the proband, parents and an unrelated healthy control as previously described[119]. Platelet imaging was performed using a structured illumination microscope (SIM, Elyra S.1, Zeiss, Heidelberg, D.E). Images were analyzed with ZEN Black (Zeiss, Heidelberg, DE). Images were analyzed with ImageJ software (National Institutes of Health, Bethesda, Maryland, U.S.A) using the 'Analyze Particles' plugin for automated analysis. Total protein lysates were obtained from platelets as described in a previous publication[120]. Protein fractions were resolved by SDS–polyacrylamide gel electrophoresis, and blots were incubated with the indicated antibodies. Membranes were next incubated with HRP-conjugated secondary antibody, and staining was performed with the ECL detection reagent (Life Technologies). Chemiluminescent blots were imaged with the ChemiDoc MP imager, and the ImageLab software version 4.1 (Bio-Rad) was used for image acquisition.

Hematopoietic stem cell differentiation assay. CD34+ hematopoietic stem cells (HSC) were isolated by magnetic cell sorting (Miltenyi Biotec) from peripheral blood from the proband, his parents and an unrelated control. The recovered (differentiation day 0) CD34+ stem cells were cultured in StemSpan SFEM medium with StemSpan CC100 ensuring strong expansion of HSC for 3 days (Stem Cell Technologies, Vancouver, C.A). Differentiation was initiated by adding 50ng/ml thrombopoietin (TPO), 25ng/ml stem cell factor and 10ng/ml interleukin 1β (Peprotech, Rocky Hill, New Jersey, U.S.A). Proplatelet formation counting (on total differentiation day 12) was performed as described in previous publications[119,83]. For immunostaining MK were seeded for 4 hours on fibrinogen-coated coverslips and stained cells were photographed at 63x magnification with a confocal microscope (AxioObserver.Z1, Zeiss, Heidelberg, D.E).

Statistical analysis. All data were analysed using GrahpPad Prism7. The details of sample sizes, statistical methods, and *P*-values are listed in the figures or figure legends.

*Homozygous deletion of CTCF binding sites in the first intron of LRBA*

*LRBA* is a member of the family of genes encoding BEACH-domain containing proteins and it has recently been identified as a novel diagnostic-grade gene for the PID domain[121]. Homozygous coding loss of function mutations in *LRBA* cause a syndrome characterized by early onset hypogammaglobulinemia and autoimmunity. We identified an unresolved PID case carrying a homozygous deletion of a CTCF binding site in an element proximal to the *LRBA* promoter. The patient presented with a mild pancytopenia, characterised by neutropenia and autoimmune haemolytic anaemia, occasionally complicated by periods of thrombocytopenia. The clinical features of this PID case are compatible with reduced LRBA function and it is thus plausible that the deleted CTCF-binding element is causally implicated in the patient's pathologies.

**Identifying causal SNVs in elements regulating diagnostic-grade genes**

We sought to identify rare disease-causing non-coding SNVs. To achieve this, we examined rare SNVs with a CADD phred score > 20 overlapping a regulatory element of a diagnostic-grade gene associated with a recessive disorder. Both the cell type in which the element was called and the patient had to be labelled with the same domain (**Supplementary Table 3**). Furthermore, the patient had to carry a high-impact rare allele such as a deletion or premature stop in the body of the diagnostic-grade gene. Application of this procedure yielded two candidate SNVs in elements targeting *AP3B1* and *MPL*. The SNV in the *MPL* element was followed up for further analysis.

*An SNV in the promoter of MPL, combined with deletion of exon 10 of MPL*

*MPL* encodes the receptor for the MK growth and development factor thrombopoietin[122]. Homozygote or compound heterozygote coding loss-of-function mutations of *MPL* cause chronic amegakaryocytic thrombocytopenia (CAMT)[123]. CAMT is categorised as type 1 or type 2 according to the severity of the thrombocytopenia and of the ensuing bone marrow aplasia. The bioinformatical approach outlined above highlighted a thrombocytopenic 10-year-old male carrying a single exon heterozygous deletion of *MPL* (chr1:43,814,723-43,815,177) and a heterozygous SNV with a CADD score of 21.8 that is absent from gnomAD. The SNV lies in a strong MK-specific regulatory element (**Extended Data Figure 10a**). Motif analysis using MatInspector[124] predicted binding of HIF1 to the wild type sequence (GGAC**G**TGGGGCT) through the well-characterised recognition site “RCGTG”, but not to the mutant sequence (GGAC**A**TGGGGCT).

The patient presented in the first months of life with a rash and a platelet count of 45 x $10^9$/L (reference range 150–450 x $10^9$/L). This low count was thought to be secondary to viral infection at the time it was measured and no further investigations were performed. At the age of 4 years, however, a full blood count (FBC) following a consultation for his attention deficit hyperactivity disorder, which at that time was assumed to be due to a delivery-related trauma, revealed that the patient's thrombocytopenia was chronic. A bone marrow aspirate showed a lower than usual number of MKs, adequate erythroid precursor cells, plentiful myeloid precursor cells and no signs of myelofibrosis. A clinical diagnosis of a CAMT-like condition was made but it could not be genetically confirmed because a second coding allele in *MPL* could not be found. The mother of the proband carried the large deletion (**Extended Data Figure 10b-d**). She and the father are healthy and have FBC results within normal ranges. The *MPL* regulatory SNV (chr1:43803414 G>A) is in trans of the large deletion because it is absent in the mother and therefore was inherited from the father or is a *de novo* variant.

A luciferase reporter assay was performed to characterise the activity of an *MPL* promoter fragment (chr1:43,803,336-43,803,488) containing the wild type G allele (MPL-SNV-G) relative to that of containing the variant A allele (MPL-SNV-A). The presence of the wild type MPL promoter fragment resulted in enhanced luciferase expression that was approximately 50% reduced in the presence of the A allele (**Extended Data Figure 10e)**. Measurement of the abundance of MPL on platelets of the proband and of the mother by flow cytometry using a specific monoclonal antibody showed markedly reduced levels in the proband compared to levels in controls and in the mother (**Extended Data Figure 10f)**.

In conclusion, absence of the MPL protein due to coding loss-of-function variants on both *MPL* alleles causes type 1 CAMT. The case reviewed above had a chronic thrombocytopenia but the other blood cell lineages seemed unaffected. This clinical phenotype is compatible with the hypothesis that the residual abundance of MPL in this case is sufficient to prevent haematopoietic stem cell exhaustion, which is the hallmark of type 1 CAMT.

*Details of the luciferase reporter assay*
The *MPL* promoter fragments chr1:43,803,336–43,803,488 containing the G- and A-allele (MPL-SNV-G and MPL-SNV-A), respectively, were cloned in pGL3-luciferase (Promega, Madison, WI, USA). Co-transfection assays in K562 cells were performed with pGL3-empty, pGL3-MPL-SNV-G or MPL-SNV-A in combination with pEGFP (Clontech, Mountain View, CA, USA) using the Amaxa electroporation system according to the manufacturer's instructions (method X-01; Lonza AG, Cologne, Germany). Cell extracts were prepared after 48 hours using 1x reporter lysis buffer and 40µl of cell extract was mixed with 50µl of luciferase assay reagent (Promega, Madison, WI, USA) to determine luciferase activity in a luminescence counter (EG&G Berthold). Each plasmid was assayed in four separate transfection experiments. The firefly luciferase activity was standardized with GFP expression as internal control. The data were analysed using GrahpPad Prism7.

**Alternative variant datasets for versions 37 and 38 of the human reference genome**
In order to migrate the 13,187 whole genomes to the human genome reference GRCh38, the Genalice high performance NGS secondary analysis suite[16] was deployed. The sequencing reads were extracted from the bam files delivered by Illumina and for comparison mapped against both assemblies, GRCh37 and GRCh38, using Genalice Map (v2.5) using default parameters[125]. The mapped reads were stored in a proprietary file format, called Genalice Aligned Reads (GAR). SNVs and indels were subsequently called using the Genalice Population calling tool in single sample mode. The variants were collated together in a single Genalice Variant Map (GVM) with blocks of reference matching positions and quality metrics (i.e. calling quality, genotype likelihoods, etc.). All genomes were processed against either of the assemblies in less than 14 days (read mapping: 12 days; genotyping: 2 days) using 10 compute nodes (Intel(R) Xeon(R) Gold 6142 CPU, 32x 2.60GHz). Variants were then exported in AVRO and standard VCF file format for variant annotation and further downstream analysis.

### Appendix

*Appendix 1: Neuropathic pain disorders*

Neuropathic pain arises as a consequence of a disease or lesion in the somatosensory nervous system[126]. A number of extreme neuropathic pain phenotypes, caused by rare high impact genetic mutations have recently been described[127]. Identification of such mutations has implications for diagnosis, genetic counselling and in some cases personalised treatment[128]. In broader terms such mutations help us understand the pathophysiology of neuropathic pain with implications for more common acquired neuropathic pain disorders, such as painful diabetic neuropathy[129]. Loss of sensation can be caused by inherited sensory neuropathies with sensory loss restricted to pain (congenital insensitivity to pain) but it may also include it may also include large fibre modalities such as touch (hereditary sensory neuropathy). Mutations in ion channels are increasingly recognised as causes of functional disorders of somatosensation[127]. For instance, homozygous loss-of-function mutations in *SCN9A*, the gene that encodes the sodium channel ($Na_v$) 1.7, have been shown to cause congenital insensitivity to pain[130]. Conversely, heterozygous gain of function variants are associated with a number of inherited pain disorders that include inherited erythromelalgia (IEM)[131] and paroxysmal extreme pain disorder (PEPD)[132]. *SCN9a* variants have also been linked to idiopathic small fibre neuropathy[133]. Some of these variants are relatively common in the general population and their penetrance may depend on environmental factors.

Our goals were to aid genetic diagnosis of patients with NPD within the UK, to determine the prevalence of known mutations associated with NPD in relation to distinct clinical presentations and finally to discover novel mutations causing NPD. The aim of our study was to determine whether genes previously implicated in neuropathic pain caused their clinical presentation. We recruited singleton individuals with extreme neuropathic pain phenotypes (both sensory loss and gain), all within the UK, from secondary care clinics located in Oxford, London, Salford, and Newcastle. We included participants older than 18 years of age with a proven history of life-style altering sensory disorder (either pain or loss of sensation) for greater than three months. The criteria for case definition for different clinical presentations are shown in **Supplementary Table 1 (NPD Criteria – Diagnostic Criteria)**. We excluded patients with a known underlying genetic cause of chronic pain, e.g. Fabry's disease and *SCN9A* congenital erythromelalgia, although genetic pre-screening for these disorders was not mandatory. Patients with a learning disorder or/and autistic features sufficient to prevent the giving of consent or participation in additional pain phenotyping were also excluded. The outcome measures used for patient phenotyping are shown in **Supplementary Table 1 (NPD Criteria Outcome Measures)**. The Neuropathic Pain Special Interest Group (NeuPSIG) of the International Association for the Study of Pain (IASP)'s grading for neuropathic pain[6] was used to grade neuropathic pain for all study participants recruited.

A total of 193 study participants with WGS data underwent neuropathic pain grading. We excluded five participants because they were unaffected family members and one participant for whom phenotypic data were unavailable.

1. **No Neuropathic pain** – 8 (4.1%) participants did not report neuropathic pain.

2. **Neuropathic pain unlikely** – 1 (0.5%) the participant's history and pain distribution was not consistent with neuropathic pain.
3. **Possible Neuropathic pain** – 11 (5.7%) participants reported an appropriate history of a relevant lesion or disease, AND pain with a distinct neuroanatomically plausible distribution.
4. **Probable Neuropathic pain** – 56 (29.0%) participants satisfied criteria for possible neuropathic pain AND had clinical signs in the neuroanatomical distribution of neuropathic pain.
5. **Definite Neuropathic pain** – 111 (57.5%) participants satisfied criteria for probable neuropathic pain AND a diagnostic test confirmed a lesion of the somatosensory nervous system.

*Appendix 2: Extreme red cell traits in UK Biobank*

Selection of participants for sequencing. The UK Biobank is a biomedical cohort of approximately half a million participants, recruited in the UK between 2006 and 2010[1]. The participants, 54% of whom are women, were aged between 37 and 73 years at their date of recruitment. Each participant underwent a baseline assessment at one of 21 centres across Great Britain, during which 4 ml of EDTA treated peripheral whole blood was collected for FBC analysis[12]. These blood samples were stored at 4 degrees centigrade and transported overnight in temperature controlled shipping boxes to the UK Biocentre laboratory in Stockport, Greater Manchester, UK, where FBCs were measured using a bank of four Beckman Coulter LH-700 instruments.

A GWAS of the FBC traits based on approximately 173,000 of the European ancestry participants in the UK Biobank cohort (n~133,000) and the INTERVAL trial (n~40,000)[70] has previously identified 582 genetic variants independently associated with quantitative properties of mature red blood cells[95]. Most of these variants were common, with only 40 having an in sample MAF lower than 1%. The GWAS design had limited power to identify associations with rare variants for three reasons. Firstly, it relied on the imputation of rare alleles from the UK10K/1000 Genomes reference panels[46,28], which are too small to contain a large proportion of the rare haplotypes carried by the hundreds of thousands of GWAS participants. Secondly, in an attempt to inhibit spurious associations due to statistical model misspecification, participants with extreme phenotype data were deliberately excluded from the association analyses. A genetic association with a rare variant can only be detected with high probability if its effect size is large, which implies that carriers of rare alleles exhibiting detectable associations were more likely than typical study participants to have been excluded from the GWAS analyses. Thirdly, the GWAS relied on univariable genetic analyses to identify allelic associations, and these can have less power than methods such as BeviMed, which are able to model jointly the association of multiple rare variants in a DNA sequence element[93].

We sought to identify a subset of UK Biobank participants likely to carry rare alleles modulating properties of peripheral red blood cells, which could plausibly be identified by WGS. Our method was to construct a univariable composite quantitative phenotype from the UK Biobank baseline mature red cell FBC measurements, which we thought likely to have high rare-variant heritability. We then selected participants for sequencing from each tail of the distribution of the phenotype.

To construct the phenotype, we used the 65 variants with MAF < 1% that were reported to be significantly ($P < 8.31 \times 10^{-9}$) associated with at least one of twelve quantitative properties of (mature or immature) red cells by[95], as a model for the likely effect of rare alleles on the baseline UK Biobank FBC. **Figure 4a** shows the pairwise relationships between the estimated effect sizes of these variants on the red cell FBC parameters MCV, RBC#, HGB and RDW. This subset of parameters is minimal in the sense that the other mature red blood cell FBC parameters can be calculated deterministically from it. The estimated effect sizes were reported by[95] as per allele additive differences in the mean of the rank-inverse standard unit normalised trait and are therefore given in units which are comparable across traits. In general, MCV and RBC# exhibit a greater range and variance in absolute effect size than HGB and RDW. Astle *et al.*[95] reported that, of all the traits they studied, MCV has the highest estimated common variant heritability and that it yielded the second largest number of associated variants with MAF < 1%. It seems reasonable to conjecture from this, that MCV also has a relatively high *rare* variant heritability. There is a strong inverse correlation between the effect sizes of alleles perturbing MCV and RBC#, suggesting that the effect sizes may measure aspects of the same underlying biological mechanism, perhaps the control of the total blood volume proportion of red cells (haematocrit). Since we could not identify any other precise systematic relationship between aspects of the joint distribution of the four estimated red cell trait effect sizes, we decided to restrict our attention to the marginal joint distribution of MCV and RBC# effect sizes (highlighted by the red square, **Figure 4a**). We used Deming regression to estimate the approximate linear relationship:

$$\beta_{MCV} = -1.69 \times \beta_{RBC\#} \qquad (1)$$

between the effect sizes for the two traits. This linear relationship is shown by a red line in **Figure 4a.**

We took the baseline UK Biobank MCV and RBC# parameters and adjusted them to remove the effect of various sources of technical and biological variation. A detailed description of the adjustments can be found in the STAR methods section of[95]. In brief, the phenotypes were firstly adjusted to remove differences between instruments, to remove time dependent instrument drifts and to remove the effect of delay time between venepuncture and measurement. Data acquired on days where the instrument mean was an outlier for the corresponding trait were removed. Subsequently, participants who had a hysterectomy or who had a self-report or medical history containing a record of myelofibrosis, lymphoma, leukemia, malignant lymphoma, multiple myeloma, multiple myelofibrosis or myelodysplasia, chronic lymphocytic leukemia, chronic myeloid leukemia, acute myeloid leukemia, polycythemia vera, polycythemia, a myeloproliferative disorder, essential thrombocytosis, a haematological cancer histology report, an unspecified lymphatic or general haematological neoplasm, a myelodysplastic syndrome, or an unspecified heme malignancy, monoclonal gammopathy, an unspecified hereditary haematological disorder, haemochromatosis, thalassaemia, haemophilia, sickle cell anaemia, neutropenia, lymphopenia or pancytopenia were excluded from analysis. Finally, the phenotypes were adjusted in a second stage to remove the effects of sex, age, menopause status, the interaction of sex and menopause status with age, height, weight and for the effects of history and current habits of smoking and alcohol consumption.

We excluded all participants who did not self report their ancestry as one of “British”, “White”, “Irish” or “Any other white background” in the UK Biobank baseline assessment questionnaire. We also excluded individuals whose genotypes appeared in the interim 2015 genetic data release and who were identified as having non-European ancestry by the principal components approach reported in[95].

We defined the quantitative selection phenotype $Q_i$ for the $i$th UK Biobank participant as the (signed) Euclidean distance in $R^2$ between the origin and the point $P(f_i,g_i)$, where $P$ is the orthogonal projection onto the line:

$$g = -1.69 \times f, \ (2)$$

where $f_i = f(RBC\#_i)$ and $g_i = g(MCV_i)$ for functions $f$ and $g$ that Box-Cox transform, standardise and centre the technically and biologically adjusted traits *RBC#* and *MCV* respectively, in the UK Biobank participants not hitherto excluded. In[95] the covariate adjusted traits were rank inverse normalised against $N(0,1)$ before the genetic association analyses. However, here we preferred to work with Box-Cox transformed traits in order to adjust the central part of the data towards a Gaussian without over shrinking outliers, which might reduce power. The selection phenotype can be expressed as:

$$Q_i \equiv -1.69 \times g(MCV_i) + f(RBC\#_i)$$

and its, centered and standardised distribution in male and post menopausal female UK Biobank participants is shown in sub-panel bounded by a dotted line in **Figure 4b.**

We excluded pre-menopausal females, as candidates for sequencing because of their high prevalence of anemia and because of the additional component of non-genetic variation in each red cell parameter that is induced by the menstrual cycle. We also excluded individuals with a UK Biobank report of an insufficient DNA quantity (less than 4.7ug) to generate a working stock of 130ul of 36ng/ul (TRINEAN measured concentration < 36ng/μl, PicoGreen measured concentration < 36ng/μl) or with a UK Biobank report of inconsistency between genetic and self reported sex. Finally, we excluded participants with a platelet count below $75 \times 10^9$/l, a white blood cell count below $0.5 \times 10^9$/l or a white blood cell count above $13 \times 10^9$/l.

We partitioned the 316,739 male and post-menopausal female participants with a computed phenotype value into six sex specific age groups thresholding at 53.3 years and 62 years. (These age groups divide those participants with a phenotype value, including the pre-menopausal women, into three groups, each of approximately 125,000 participants). Within each of these six sex-age groups, we ranked the study participants according to the value of the phenotype. We selected a total of 384 individuals from each tail of the phenotype, stratifying the selection by sex-age group so that the final selection for each tail sampled each group in proportion to its size. This additional stratification was necessary despite the adjustment of the mean of each trait for age and sex to ensure reasonable age and sex balance in the tails. The full blood count of each selected participant was reviewed by an expert panel of haematologists to exclude any text-book non-genetic or somatic pathologies such as bone-marrow failure, polycythemia vera or essential thrombocytopenia, which might explain the extreme value of the selection phenotype. A small sample of DNA was screened by the Cambridge Blood and Stem-Cell Biobank for the *JAK2* mutation V617F, a common cause of somatic myeloproliferative disorders. Any participants failing the FBC or DNA screen were replaced by the next most extreme individual in the same sex-age subgroup.

DNA samples from a total of 416 male and 352 female UK Biobank participants were retrieved from the central sample archive and sent to Illumina sequencing. The main panel of **Figure 4b** shows the distribution of the quantitative phenotype of the selected individuals in each tail, while **Figure 4c** is a bivariate scatter showing the distribution of RBC# and MCV (after adjustment for technical but not biological variation) in the two tails. Of the 768 individuals sent for WGS one individual from each tail failed Illumina sequencing quality control and two distinct individuals from the right tail failed in-house checks for DNA contamination.

Derivation of distribution of polygenic score in tail selected participants. Assume that the mean centred, standardised quantitative phenotype Q takes an $N(0,1)$ distribution in the population and that it may be decomposed as:

$$Q = S + U$$

where *S* is the polygenic score and *U* is the part of Q not explained by the score. The central limit theorem suggests that in sufficiently large samples the distribution of *S* should be $N(\alpha, \phi^2)$ distributed for some $\alpha,\ \phi^2$. Consequently *U* should also be normally distributed. An assumption of independence between *S* and *U* implies that $0 < \phi^2 < 1$ and that:

$$Q|S = s \sim N(s - \alpha, 1 - \phi^2)$$

Consequently,

$$\begin{aligned} p(q < \tau, s) &= \int_{-\infty}^{\tau} p(q,s)dq \\ &= p(s) \int_{-\infty}^{\tau} p(q|s)dq \\ &= \tfrac{1}{\phi}\varphi(\tfrac{s-\alpha}{\phi})\Phi(\tfrac{\tau-s+\alpha}{\sqrt{1-\phi^2}}) \end{aligned}$$

where $\varphi$ and $\Phi$ are respectively the PDF and CDF of $N(0,1)$. This implies that,

$$p(s|q < \tau) = \frac{1}{\phi}\varphi(\frac{s-\alpha}{\phi})\frac{\Phi(\frac{\tau-s+\alpha}{\sqrt{1-\phi^2}})}{\Phi(\tau)} .$$

By a symmetrical argument,

$$p(s|q > \tau) = \frac{1}{\phi}\varphi\left(\frac{s-\alpha}{\phi}\right)\frac{1-\Phi\left(\frac{\tau-s+\alpha}{\sqrt{1-\phi^2}}\right)}{1-\Phi(\tau)} .$$

Affiliation numbering continues from the main paper

**NIHR BioResource - Rare Diseases Collaborators:** Zoe Adhya[168], Maryam Afzal[42], Irshad Ahmed[169], Saeed Ahmed[228], Jayanthi Alamelu[44], Raza Alikhan[104], Louise Allen[8,83,229], Arif Alvi[108], Gautam Ambegaonkar[230], Ariharan Anantharachagan[35,231], Jamie Anderson[41], Gururaj Arumugakani[232], Rita Arya[233], Efi Athieniti[41], Steve Austin[44], Yesim Aydinok[234], Waqar Ayub[235], Mohsin Badat[33], Trevor Baglin[35], Jonathan Barratt[236], John Baski[118,119], Rachel Bates[33], Gareth Baynam[237,238,239], Claire Bethune[240], Neha Bhatnagar[107], Shahnaz Bibi[46], Preetham Boddana[241], Claire Booth[46], Angela Brady[242], Annette Briley[16], Richard Brown[243], Christine Bryson[1,2], Jackie Buck[244], Gary Campbell[245], Natalie Canham[141,242], Jenny Carmichael[35], Dahlia Castle[41], Elizabeth Chalmers[170], Melissa V Chan[246], Anita Chandra[35], Sam Chong[153], Emma M Clement[46], Virginia Clowes[242], Victoria Cookson[46], Amanda Creaser-Myers[247], Rosa Da Costa[119], Sophie Davies[35], Sarah Deacock[248], Patrick B Deegan[26], John Dempster[168], Michael Desborough[33], Lisa A Devlin[163], Anand Dixit[117], Rainer Doffinger[158], Helen Dolling[1,2], Natalie Dormand[119], Tariq El-Shanawany[172], Tony Elston[249], Ingrid Emmerson[117], Henry Farmery[6], Helen Firth[3,48], Nick Fordham[33], Bruce Furie[105], Alice Gardham[242], H Bobby Gaspar[30], Johanna Gebhart[250], Neeti Ghali[251], Rohit Ghurye[168], Rodney D Gilbert[252,253], Lionel Ginsberg[54,106,161], Joanna C Girling[254], Paul Gissen[46,54], Kathleen M Gorman[147,148], Alan Greenhalgh[255], Sian Griffiths[256], Yisu Gu[33], Robert D M Hadden[257], Csaba Halmagyi[1,2], Tracey Hammerton[1,2], Lorraine Harper[157], Claire Harrison[122], Shivaram Hegde[256], Robert H Henderson[46], Anke Hensiek[35], Yvonne M C Henskens[258], Muriel Holder[122], Sean Hughes[231], Stephen Hughes[259], Anna E Huis in 't Veld[187], Jane A Hurst[46], Val Irvine[188], Praveen Jeevaratnam[260], Mark Johnson[261], Bryony Jones[262], Caroline Jones[263], Yousuf Karim[180,248], Mahantesh Karoshi[264], David Keeling[107], Fiona

Kennedy[188], Sorena Kiani[168], Andrew King[33], Sally Kinsey[265], Alison Kirkpatrick[180], Nigel Klein[46], Ellen Knox[266], Deepa Krishnakumar[35], James Laffan[168], Sarah H A Lawman[267], Sara E Lear[35,245,268], Melissa Lees[46], Andrew Lewington[269], James Liang[270], Ri Liesner[102], Silvia Lucato Hadeler[1,2], Malcolm Macdougall[117], Rajiv D Machado[271,272], Lucy H Mackillop[33,273], Robert MacLaren[49], Laura Magee[274], Mohamed Mahdi-Rogers[121], Mike Makris[201,247], Ania Manson[35], Adnan Manzur[46], Patrick B Mark[178,275], Larahmie Masati[66], Vera Matser[1,2], Anna Maw[35], Elizabeth M McDermott[162], Simon J McGowan[27,29], Coleen McJannet[1,2], Amy McTague[147,148], Sharon Meehan[66], Catherine L Mercer[88], Anna C Michell[31,33], David Milford[276], Anoop Mistry[232], Jason Moore[277], Valerie Morrisson[35], Sai H K Murng[177,178], Elaine Murphy[106], Joanne Ng[147,148], Adeline Ngoh[147,148], Muna Noori[262], Eric Oksenhendler[278], Albert C M Ong[165,201], Shokri Othman[66], Yasmin Panchbhaya[41], Antonis Pantazis[119], Apostolos Papandreou[147,148,279], Alasdair P J Parker[35], Georgina Parsons[33], K John Pasi[280], Chris Patch[122], Jeanette H Payne[281], David Perry[69], Bartlomiej Piechowski-Jozwiak[121], Fernando Pinto[170], Gary J Polwarth[159], Mark J Ponsford[282,283], Sanjay Prasad[118,119], Waseem Qasim[30,46], Ellen Quinn[46], Isabella Quinti[284], Sanjay Raina[285], Lavanya Ranganathan[66], Julia Rankin[277], Karola Rehnstrom[1,2], Evan Reid[8,55], Mary M Reilly[54,153], Shoshana Revel-Vilk[286], Mike Richards[287], Emma E Richards[111], Matthew T Rondina[288], Elisabeth Rosser[46], Peter Rothwell[289], Jennifer G Sambrook[1,2], Richard Sandford[55], Saikat Santra[276], Gwen Schotte[187], Harald Schulze[290], Suranjith L Seneviratne[51,161], Fiona Shackley[165], Momin Shah[41], Pankaj Sharma[291], Hassan Shehata[171,292], Deborah Shipley[255], Manish D Sinha[18,19,122], Linda Sneddon[116], Aman Sohal[276], Laura Southgate[272,293], Miranda Splitt[117], Hans Stauss[161], Cathal L Steele[294], Penelope E Stein[121], Sophie Stock[1,2], Matthew J Stubbs[22,23], Emily Symington[69], Gordon B Taylor[295], Jecko Thachil[296], Dorothy A Thompson[46], Sarah Trippier[274], Rafal Urniaz[26], Marijcke W M Veltman[1,2], Julie Vogt[139], Ajay Vora[297], Minka J A Vries[258], Emma L Wakeling[242], Roddy Walsh[118,119], Ivy Wanjiku[66], Timothy Warner[168], Evangeline Wassmer[276], Henry G Watson[295], Dean Waugh[113], Nick Webb[259], Angela Welch[112], David Werring[106], Lisa Willcocks[35], David J Williams[106], Henna Wong[33], Sarita Workman[161], Nigel Yeatman[168]

[1]Department of Haematology, University of Cambridge, Cambridge Biomedical Campus, Cambridge, UK. [2]NIHR BioResource, Cambridge University Hospitals NHS Foundation, Cambridge Biomedical Campus, Cambridge, UK. [3]Wellcome Sanger Institute, Wellcome Genome Campus, Hinxton, Cambridge, UK. [6]MRC Biostatistics Unit, Cambridge Institute of Public Health, University of Cambridge, Cambridge, UK. [8]Department of Medical Genetics, Cambridge Institute for Medical Research, University of Cambridge, Cambridge Biomedical Campus, Cambridge, UK. [16]Women and Children's Health, School of Life Course Sciences, King's College London, London, UK. [18]King's College London, London, UK. [19]Department of Paediatric Nephrology, Evelina London Children's Hospital, Guy's & St Thomas' NHS Foundation Trust, London, UK. [22]Department of Haematology, Hammersmith Hospital, Imperial College Healthcare NHS Trust, London, UK. [23]Centre for Haematology, Imperial College London, London, UK. [26]Department of Medicine, School of Clinical Medicine, University of Cambridge, Cambridge Biomedical Campus, Cambridge, UK. [27]MRC Molecular Haematology Unit, MRC Weatherall Institute of Molecular Medicine, University of Oxford, Oxford, UK. [29]NIHR Oxford Biomedical Research Centre, Oxford University Hospitals Trust, Oxford, UK. [30]UCL Great Ormond Street Institute of Child Health, London, UK. [31]Department of Cardiovascular Medicine, Radcliffe Department of Medicine, University of Oxford, Oxford, UK. [33]Oxford

University Hospitals NHS Foundation Trust, Oxford, UK. [35]Addenbrookes Hospital, Cambridge University Hospitals NHS Foundation Trust, Cambridge, UK. [41]Illumina Cambridge Limited, Chesterford Research Park, Little Chesterford, Saffron Walden, Essex, UK. [42]Bristol Renal and Children's Renal Unit, Bristol Medical School, University of Bristol, Bristol, UK. [44]Department of Haematology, Guy's and St Thomas' NHS Foundation Trust, London, UK. [46]Great Ormond Street Hospital for Children NHS Foundation Trust, London, UK. [48]East Anglian Medical Genetics Service, Cambridge University Hospitals NHS Foundation Trust, Cambridge, UK. [49]Moorfields Eye Hospital NHS Foundation Trust, London, UK. [51]Institute of Immunity and Transplantation, University College London, London, UK. [54]University College London, London, UK. [55]Department of Clinical Genetics, Addenbrookes Hospital, Cambridge University Hospitals NHS Foundation Trust, Cambridge, UK. [66]Department of Medicine, Imperial College London, London, UK. [69]Department of Haematology, Cambridge University Hospitals NHS Foundation Trust, Cambridge, UK. [83]Department of Renal Medicine, Addenbrookes Hospital, Cambridge University Hospitals NHS Foundation Trust, Cambridge, UK. [88]Southampton General Hospital, University Hospital Southampton NHS Foundation Trust, Southampton, UK. [102]Department of Haematology, Great Ormond Street Hospital for Children NHS Foundation Trust, London, UK. [104]The Arthur Bloom Haemophilia Centre, University Hospital of Wales, Cardiff, UK. [105]Beth Israel Deaconess Medical Centre and Harvard Medical School, Boston, USA. [106]University College London Hospitals NHS Foundation Trust, London, UK. [107]Oxford Haemophilia and Thrombosis Centre, The Churchill Hospital, Oxford University Hospitals NHS Trust, Oxford, UK. [108]Glasgow Royal Infirmary, NHS Greater Glasgow and Clyde, Glasgow, UK. [111]Department of Neurology, Sheffield Teaching Hospitals NHS Foundation Trust, Sheffield, UK. [112]Institute of Neuroscience and Psychology, University of Glasgow, Glasgow, UK. [113]Department of Neurology, Leeds Teaching Hospital NHS Trust, Leeds, UK. [116]Newcastle University, Newcastle upon Tyne, UK. [117]Newcastle upon Tyne Hospitals NHS Foundation Trust, Newcastle upon Tyne, UK. [118]National Heart and Lung Institute, Imperial College London, London, UK. [119]Royal Brompton Hospital, Royal Brompton and Harefield NHS Foundation Trust, London, UK. [121]King's College Hospital NHS Foundation Trust, London, UK. [122]Guy's and St Thomas' Hospital, Guy's and St Thomas' NHS Foundation Trust, London, UK. [139]West Midlands Regional Genetics Service, Birmingham Women's and Children's NHS Foundation Trust, Birmingham, UK. [141]Department of Clinical Genetics, Liverpool Women's NHS Foundation, Liverpool, UK. [147]Developmental Neurosciences, UCL Great Ormond Street Institute of Child Health, London, UK. [148]Department of Neurology, Great Ormond Street Hospital for Children NHS Foundation Trust, London, UK. [153]The National Hospital for Neurology and Neurosurgery, University College London Hospitals NHS Foundation Trust, London, UK. [157]University Hospitals Birmingham NHS Foundation Trust, Birmingham, UK. [158]Division of Clinical Biochemistry and Immunology, Cambridge University Hospitals NHS Foundation Trust, Cambridge, UK. [159]Royal Papworth Hospital NHS Foundation Trust, Cambridge, UK. [161]Royal Free London NHS Foundation Trust, London, UK. [162]Nottingham University Hospitals NHS Trust, Nottingham, UK. [163]Regional Immunology Service, The Royal Hospitals, Belfast, UK. [165]Sheffield Teaching Hospitals NHS Foundation Trust, Sheffield, UK. [168]Barts Health NHS Foundation Trust, London, UK. [169]Birmingham Heartlands Hospital, University Hospitals Birmingham NHS Foundation Trust, Birmingham, UK. [170]Royal Hospital for Children, NHS Greater Glasgow and Clyde, Glasgow, UK. [171]Epsom & St Helier University Hospitals NHS Trust, London, UK. [172]Immunodeficiency Centre for Wales, University Hospital of Wales, Cardiff, IUK. [177]Gartnavel General Hospital, NHS

Greater Glasgow and Clyde, Glasgow, UK. [178]Queen Elizabeth University Hospital, Glasgow, UK. [180]Frimley Park Hospital, NHS Frimley Health Foundation Trust, Camberley, UK. [187]Department of Pulmonary Medicine, VU University Medical Centre, Amsterdam, The Netherlands. [188]Golden Jubilee National Hospital, Glasgow, UK. [201]Department of Infection, Immunity & Cardiovascular Disease, University of Sheffield, Sheffield, UK. [228]Department of Renal Medicine, Sunderland Royal Hospital, Sunderland, UK. [229]Department of Ophthalmology, Addenbrookes Hospital, Cambridge University Hospitals NHS Foundation Trust, Cambridge, UK. [230]Child Development Centre, Addenbrookes Hospital, Cambridge University Hospitals NHS Foundation Trust, Cambridge, UK. [231]Lancashire Teaching Hospital NHS Foundation Trust, Lancashire, UK. [232]The Leeds Teaching Hospitals NHS Trust, Leeds, UK. [233]Warrington and Halton Hospitals NHS Foundation Trust, Warrington, UK. [234]Ege University Hospital, Department of Paediatric Hematology-Oncology, Izmir, Turkey. [235]University Hospitals Coventry and Warwickshire, Coventry, UK. [236]Infection, Immunity and Inflammation, University of Leicester, Leicester, UK. [237]School of Paediatrics and Child Health, University of Western Australia, Perth, Australia. [238]Genetic Services of Western Australia, Western Australian Register of Developmental Anomalies and Office of Population Health Genomics, Public and Aboriginal Health Division, Western Australian Department of Health, Perth, Australia. [239]Genetic and Rare Diseases Program, Telethon Kids Institute, Perth, Australia. [240]University Hospitals Plymouth NHS Trust, Plymouth, UK. [241]Gloucestershire Royal Hospital, Gloucestershire Hospitals NHS Foundation Trust, Gloucester, UK. [242]North West Thames Regional Genetic Service, London North West University Healthcare NHS Trust, Harrow, UK. [243]Department of Neurology, Addenbrookes Hospital, Cambridge University Hospitals NHS Foundation Trust, Cambridge, UK. [244]NHS, NHS Trust, UK. [245]Norfolk and Norwich University Hospitals NHS Foundation Trust, Norwich, UK. [246]Blizard Institute, Barts and The London School of Medicine & Dentistry, Queen Mary University of London, London, UK. [247]Royal Hallamshire Hospital NHS Foundation Trust, Sheffield, UK. [248]Royal Surrey County Hospital NHS Foundation Trust, Guildford, UK. [249]Colchester Hospital University NHS Foundation Trust, Colchester, UK. [250]Medical University of Vienna, Vienna, Austria. [251]National Ehlers–Danlos Syndrome Diagnostic Service, Northwick Park Hospital, London, UK. [252]Southampton Children's Hospital, University Hospital Southampton NHS Foundation Trust, Southampton, UK. [253]Faculty of Medicine, University of Southampton, Southampton, UK. [254]West Middlesex University Hospital, Chelsea and Westminster Hospital NHS Foundation Trust, London, UK. [255]Freeman Hospital, The Newcastle upon Tyne Hospitals NHS Foundation Trust, Newcastle upon Tyne, UK. [256]University Hospital of Wales, Cardiff, UK. [257]Department of Neurology, King's College Hospital NHS Foundation Trust, London, UK. [258]Maastricht University Medical Centre, Maastricht, The Netherlands. [259]Royal Manchester Children's Hospital, Manchester University NHS Foundation Trust, Manchester, UK. [260]Lister Hospital, East and North Hertfordshire NHS Trust, Stevenage, UK. [261]Chelsea and Westminster Hospital NHS Foundation Trust, London, UK. [262]Queen Charlotte's and Chelsea Hospital, Imperial College Healthcare NHS Trust, Du Cane Road, London, UK. [263]Alder Hey Children's Hospital, Liverpool, UK. [264]Barnet General Hospital, Royal Free London NHS Foundation Trust, London, UK. [265]Leeds General Infirmary, Leeds Teaching Hospitals NHS Trust, Leeds, UK. [266]Birmingham Women's Hospital, Birmingham Women's and Children's NHS Foundation Trust, Birmingham, UK. [267]Sussex Kidney Unit, Royal Sussex County Hospital, Brighton and Sussex University Hospitals, Brighton, UK. [268]Addenbrooke's Treatment Centre, Addenbrooke's Hospital, Cambridge University Hospitals NHS Foundation Trust, Cambridge,

UK. [269]Renal Medicine, Leeds Teaching Hospitals NHS Trust, Leeds, UK. [270]Middlemore Hospital, Auckland, New Zealand. [271]School of Life Sciences, University of Lincoln, Lincoln, UK. [272]Molecular and Clinical Sciences Research Institute, St George's University of London, London, UK. [273]Nuffield Department of Women's and Reproductive Health, Oxford University Hospitals NHS Trust, Oxford, UK. [274]St George's University Hospitals NHS Foundation Trust, London, UK. [275]Institute of Cardiovascular and Medical Sciences, University of Glasgow, Glasgow, UK. [276]Birmingham Children's Hospital, Birmingham Women's and Children's NHS Foundation Trust, Birmingham, UK. [277]Royal Devon and Exeter NHS Foundation Trust, Exeter, UK. [278]Department of Clinical Immunology, Hopital Saint-Louis, Assistance Publique-Hopitaux de Paris, University Paris Diderot, Sorbonne Paris Cite, Paris, France. [279]UCL MRC Laboratory for Molecular Cell Biology, London, UK. [280]Barts and The London School of Medicine and Dentistry, Haemophilia Centre, The Royal London Hospital, London, UK. [281]Dept of Haematology, Sheffield Children's Hospital NHS Foundation Trust, Sheffield, UK. [282]Cardiff University, Cardiff, UK. [283]Immunodeficiency Centre for Wales, Heath Hospital, Cardiff, UK. [284]Department of Molecular Medicine, Sapienza University of Rome, Rome, Italy. [285]The Princess Alexandra Hospital NHS Trust, Harlow, UK. [286]Shaare Zedek Medical Center, affiliated with Hebrew-University Medical School, Jerusalem, Israel. [287]Leeds Children's Hospital, The Leeds Teaching Hospitals NHS Trust, Leeds, UK. [288]Department of Internal Medicine, Eccles Institute of Human Genetics, University of Utah Health Sciences Center, Salt Lake City, USA. [289]Stroke Prevention Research Unit, University of Oxford, Oxford, UK. [290]Experimental Biomedicine, University Hospital Würzburg, Würzburg, Germany. [291]Institute of Cardiovascular Research Royal Holloway University of London (ICR2UL), London, UK. [292]Epsom General Hospital, Epsom, UK. [293]Faculty of Life Sciences and Medicine, King's College London, London, UK. [294]Ninewells Hospital and Medical School, NHS Tayside, Dundee, UK. [295]Aberdeen Royal Infirmary, NHS Grampian, Aberdeen, UK. [296]Haematology Department, Manchester Royal Infirmary, Central Manchester University Hospitals National Health Service Foundation Trust, Manchester Academic Health Science Centre, Manchester, UK. [297]Sheffield Children's Hospital NHS Foundation Trust, Sheffield, UK
